# Computational pipeline reveals nature’s untapped reservoir of halogenating enzymes

**DOI:** 10.64898/2026.01.20.700248

**Authors:** Judit Szenei, Ashleigh Burke, Anne Liong, Aleksandra Korenskaia, April L. Lukowski, Nadine Ziemert, Pablo I. Nikel, Pedro N. Leão, Bradley S. Moore, Tilmann Weber, Kai Blin

## Abstract

Microbial halogenated natural products (hNPs) hold ecological, agricultural, and biomedical relevance. The hNP-producing potential of the organism can be assessed by the precise prediction of biosynthetic enzymes, yet the detailed annotations of halogenases are often missing from genomic and metagenomic data. We created a manually curated database (https://halogenases.secondarymetabolites.org/) containing information on the halide-specificity, role, and position of verified catalytic residues and results of the mutagenesis studies of more than 120 experimentally validated or *in silico* inferred halogenases. The collection of experimental data supports a computational pipeline that allows the family-, substrate-, and halide-scope-level annotation of halogenating enzymes by relying on catalytic residues, conserved motifs, and profile Hidden Markov Models (pHMMs). Our analysis with sequence similarity networks (SSNs) highlighted several underexplored clusters in the UniRef50 database. Such finding was a halogenase from *Rhodopirellula baltica* (*Rhoba*VHPO) previously labelled as a hypothetical chloroperoxidase, which clustered apart from the known chloroperoxidases and bromoperoxidases, but accepted chloride and preferred bromide. Our database and workflow provide extensive and scalable solutions for the systematic and precise annotation of halogenating enzymes in genomic and metagenomic data. The in-depth categorization of halogenases will improve the chemical structure prediction of microbial hNPs, supporting ecological assessments and natural product discovery.

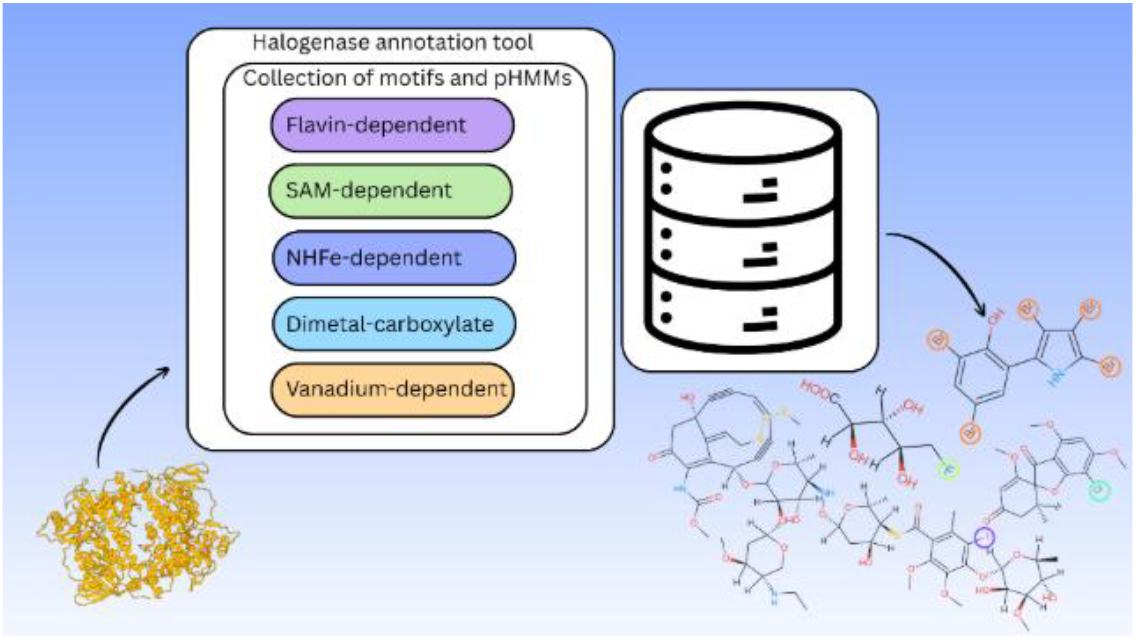

## INTRODUCTION

Microbes possess a large palette of halogenating enzymes, making them the driving factor of halogen cycling in marine and terrestrial environments [1,2]. Leaning on the precise activity prediction of halogenases helps us assess the variety of potential halogenated natural products (hNPs) which enter the biogeochemical cycles in these habitats. The activity prediction can be aided by identifying known functionally important residues in the enzyme sequence that reflect the versatile catalytic mechanisms among halogenases. Halogenating enzymes install chlorine, bromine, iodine or fluorine through electrophilic, nucleophilic or radical substitution reactions [3,4] utilizing different cofactors.

### Electrophilic substitution

Halogenation through electrophilic substitution happens with the help of a heme-oxo-Fe(IV) species, reduced flavin or a vanadate-center by heme-dependent, flavin-dependent, or vanadium-dependent enzymes [3], respectively. The heme-dependent family provided the first described halogenase, produced by *Caldariomyces fumago* [4], that remained the only representative of this group with available crystal structure [5]. In comparison, flavin-dependent halogenases (FDHs) are widespread across the kingdom of Fungi, Bacteria and Archaea with crystal structures available for more than 20 enzymes in the Protein Data Base [6]. Two-component FDHs are most common in sequence and structure databases [7], requiring a flavin-reductase partner, but recent studies revealed single-component members of the family which function without a coenzyme [8,9]. The substrate scope of FDHs extends to tryptophan, phenol and pyrrole [10], which can be free-standing (variant A enzymes) or tethered to a carrier protein (variant B enzymes) [3, 11]. A handful of FDHs accept aliphatic substrates [12, 13], but whether it is enabled through electrophilic substitution or radical mechanism [14] is debated and we see electrophilic reaction happening mostly on aromatic compounds, also in the case of vanadium-dependent halogenases (VHPOs) [15, 16]. While FDHs are the most known for their regio-and substrate-selectivity, VHPOs also show selective nature [16].

### Nucleophilic substitution

The group of nucleophilic halogenases contains attractive biocatalysts, the S-Adenosyl-L-Methionine (SAM)-dependent enzymes [3], the only known family so far which catalyzes fluorination in nature [17]. The C-terminal with RNAA motif sets the fluorinases apart from members of the family which cannot utilize fluoride. Actinomycetes hold the most fluorinating enzymes [17, 18], but in silico mining yielded an additional fluorinase from Archaea with a YYGG motif at the C-terminal [19].

### Radical substitution

The group of radical halogenases contains several families capable of acting on unactivated carbon. Non-heme iron alphaketoglutarate (NHFe)-dependent members are the most covered within the group of radical halogenases, including both variant A [20, 21] and variant B members [22]. Even though the distribution of dimetal-carboxylate halogenases exceeds the NHFe-dependent ones in cyanobacteria [23], the family gained attention only in the past years, mostly from cyanotoxin biosynthetic pathways [24]. It is the group most abundant in enzymes acting on fatty acids [25], while from the NHFe-dependent family only HctB from hectachlorin biosynthesis is known for that substrate scope [26, 27].

The group of radical halogenases expanded recently with the discovery of ApnU, a copper-dependent enzyme from atpenin biosynthesis [28], polyhalogenating unactivated carbon atoms, like the iron-dependent families.

### Annotation of halogenases

Halogenases showcase a wide variety of cofactor requirements which lead to distinct catalytic cycles, recognizable based on the functionally important amino acids. Relying on catalytic residues provides support for detailed predictions such as inferring the substrate scope, halide scope and selectivity. While the bioacatalyst mining tool EnzymeMiner [29] allows the specification of functionally important residues, annotation tools such as BLAST [30], or solely profile Hidden Markov Model (pHMM)-based [31] solutions, like HHPred [32] and InterPro profiles [33], rely only on homology. Even though homology can guide us to the substrate scope e.g. for flavin-dependent tryptophan halogenases [34], pHMMs integrated in InterPro are often too broad to infer substrate scope or selectivity. Despite the abundant information on catalytic residues and conserved motifs from experimental studies, there was no tool which integrated this knowledge across several halogenase families and neglecting catalytic information often leads to ambiguous or incorrect annotation.

### Improved halogenase annotation

We aimed to close the gap in the differing annotation tool availability for underexplored halogenase families and improve the annotation by collecting the catalytic residues and motifs and by constructing activity-specific pHMMs. We demonstrated the potential of the computational pipeline by applying it on UniRef50 [35]. Furthermore, a haloperoxidase from *Rhodopirellula baltica* (*Rhoba*VHPO) caught our attention due to its clustering with VHPOs that possess characteristic iodoperoxidase catalytic residues. Monochlorodimedone (MCD) assay confirmed its annotation as chloroperoxidase and uncovered its bromide preference that was five-fold greater to that of chloride. The developed computational pipeline and database support the in-depth annotation of halogenating enzymes across six enzyme families, enabling the more accurate prediction of the chemical structure of hNPs and the mining of novel biocatalysts.

## RESULTS AND DISCUSSION

### Halogenase Data Base

To address the annotation gap among halogenating enzymes, we manually curated the Halogenase DB, which holds information of experimentally validated or *in silico* inferred halogenases. The first layer of the DB contains the accession, producing organism, protein name and which halides can or cannot be utilized. The second layer of the database includes the residues relevant to the structure or the catalytic cycle of the enzyme. Furthermore, the position of these residues in the sequence, as well as in the pHMM suitable for the given enzyme is found also in the second layer.

The JSON format of the DB allows easy access to the data and smooth scalability as new halogenases are being explored.

The open-source Halogenase DB (https://halogenases.secondarymetabolites.org/) consists of 125 entries, more than double the halogenase entries in SwissProt. Copper-dependent (1 sequence), dimetal-carboxylate (7 sequences), SAM-dependent (7 sequences), NHFe-dependent (22 sequences), FDHs (65 sequences), VHPOs (22 sequences) are represented in different distributions depending on the available information in literature. Validation details are available for 94 enzymes and mutagenesis, or structure analysis studies were available for 50 enzymes.

The Halogenase DB serves as first-line information when someone starts working with a target enzyme and highlights the pitfalls that might arise in the experimental process. While in the case of VHPOs, the naming follows a systematic convention where it indicates the most electronegative halide they accept, this standardization is missing in other families. It is also worth noting that despite the rule-of-thumb naming of VHPOs, there are examples in *Acaryochloris marina* (*Am*VBPO) and in *Ascophyllum nodosum* (*An*VBPO), where the VHPOs labelled as bromoperoxidase form chlorinated compounds [36, 37]. The chlorination activity is vastly different in the two species as *An*VBPO tetrachlorinates phenolsulfonephthalein [38], while *Am*VBPO required such high chloride concentrations in the MCD assay that the kinetic studies couldn’t be completed [39].

Therefore, the database includes information about the tested halides and the halide preference if this information is available. Out of 125 entries only around 10 enzymes were tested for fluoride.

### Collection of catalytic residues and conserved motifs

The computational pipeline provides a transparent method for in-depth halogenase annotation while leveraging the results of experimental studies. The conserved motifs and catalytic residues inferred through *in silico* analysis and experimental validation support the family-level categorization for both FDHs and dimetal-carboxylate enzymes. The annotation of flavin-dependent enzymes is based on WxWxI, a motif which prevents the monooxygenation activity and Fx.Px.Sx.G, a motif which marks the tunnel in the protein structure. In the case of the dimetal-carboxylate family, the annotation paradigm considers the ExxQExxH and HxxDExxH motifs (Table 1).

**Table 1.** Small motifs and catalytic residues. *Flavin-dependent* GxGxxG marks the flavin-binding site in FDHs, followed by WxWxI(P) [3] in two-component halogenases. The second motif might be responsible for preventing monooxygenation activity [3]. A broader pattern, Fx.Px.Sx.G takes place towards the C-terminal of the sequence [3]. This motif marks the tunnel between the bound cofactor and the site of halogenation [3]. In the single-component members of the family, the GXGXXA motif at the N-terminal seems to be conserved (SI) ***NHFe-dependent*** The presence of HXG/A, also referred to as catalytic triad indicates halogenation reaction [41]. ***Dimetal-carboxylate*** They are mined based on two motifs containing catalytic Glu and Asp residues [42]. ***SAM-dependent*** The C-terminal RNAA (in bacteria) and YYGG (in archaea) motifs (SI) indicate fluorination activity [19]. ***Vanadium-dependent*** There are no established patterns to mine for VHPO(s). However, the presence of a catalytic Lys and Ser residue indicates selective nature and helps differentiate from close non-halogenase relatives [40]. ***Copper-dependent*** Two HXXHC motifs indicate the copper-binding site [28]. The two Cys residues (Cys173 and Cys200) probably do not participate in the metal-binding site formation but create one of the two intramolecular disulfide bridges in the structure [28]. **Curated pHMMs.** After generating MSAs for each of the targeted halogenase families, we curated the pHMMs. The number of pHMMs differs for each group of halogenases, depending on the available sequence data and the similarity between enzymes with distinct substrate-and halide-scope (Supplementary Table 1). We did cross-validation to define thresholds which can be used as guiding points to exclude enzymes with different activity compared to the target-group of the pHMM.

| Enzyme family | Motif/Catalytic residue | Role |
| --- | --- | --- |
| Flavin-dependent | GXGXXG | Flavin-binding site |
| Flavin-dependent | GXGXXA | Conserved motif in two-component FDHs |
| Flavin-dependent | WXWXI | Prevents monooxygenation activity |
| Flavin-dependent | FX.PX.SX.G | Key residues in the tunnel between the flavin binding site and the site of halogenation both in two-and single-component FDHs |
| NHFe-dependent | HXG/A | Catalytic triad |
| Dimetal-carboxylate | EXXQEXXH | Contains the catalytic Gln specific to halogenases |
| Dimetal-carboxylate | HXXDEXXH | Contains the putative active site |
| SAM-dependent | RNAA | C-terminal motif in bacterial fluorinases |
| SAM-dependent | YYGG | C-terminal motif in archaeal fluorinase |
| Vanadium-dependent | K324; S427 (in NapH1) | Characteristic to regio/stereo-selective chloroperoxidases |
| Copper-dependent | HXXHC; HXXHC | The two motifs indicate the metal-binding site |

The family-level categorization of NHFe-dependent, SAM-dependent and vanadium-dependent enzymes is homology-based with options to search for specific residues in validated enzymes. A C-terminal motif supports the identification of potential fluorinases both for bacteria and for archaea in the SAM-dependent family and a smaller motif marking the catalytic triad helps the differentiation of halogenases and dioxygenase/hydroxylase homologs of NHFe-dependent enzymes (Table 1). There are no conserved motifs for family-level annotation in case of VHPOs, but the presence of a catalytic Lys and Ser residue indicates the stereo/regio-selectivity of our target enzyme [40] (Table 1).

The prediction of the number of halogenation events for FDHs is supported for pyrrole halogenases (Supplementary Information (SI)). For VHPOs, the prediction of potential iodination activity and the selectivity of the enzyme rely on the residue collection (SI).

As the catalytic residues for halogenation activity among the copper-dependent enzymes are not known yet, we cannot set them apart from the hydroxylase homologs, but family-level annotation is possible by searching for two HXXHC motifs [28].

For the SAM-dependent family, the pHMMs predict fluorination and non-fluorination activity. Non-fluorinating enzymes are often labelled as chlorinases, stemming from the fact that they are not routinely tested for bromide or iodide. The only known example of a SAM-dependent halogenase tested for all four halides is SaIL, a halogenase from *Salinospora tropica*, which accepts all halides, except fluoride [43]. Using the labels “fluorinase” or “non-fluorinase” helps to describe the halide-utilization more clearly.

Categorization in the NHFe-dependent family branches into variant A and variant B enzymes. For variant A enzymes, the workflow includes substrate-scope predictions by relying on nucleotide-, indole alkaloid-, and small amino acid-specific pHMMs. The pHMM-based substrate prediction is feasible for variant A enzymes as they cluster based on substrate-scope on the phylogenetic tree (Supplementary Figures (SF)). The workflow does not consider catalytic residues or conserved motifs, and we recommend using the features of the workflow where the user can compare the sequences to validated enzymes in which the catalytic residues were inferred. As the residues and the structural motifs are unknown for deciding between variant A and variant B groups, the pHMMs provide a new layer of categorization. Due to the limited number of experimental studies, the in-depth categorization of dimetal-carboxylate halogenases is not possible. Therefore, the workflow for diiron halogenases leans on a general pHMM built from the MSA of the currently known dimetal-carboxylate enzymes and the two motifs containing the catalytic Asp and Glu residues. As the ExxQExxH motif is missing in AurF [EC: 1.14.99.68] (4-aminobenzoate N-oxygenase) and its homologs, the workflow can give indication of oxygenation activity as well. AurF did not hit the halogenase pHMM during cross-validation, therefore a cutoff is not necessary to differentiate from the oxygenase homologs. However, searching for the conserved motif containing the active site can strengthen confidence in the categorization.

While a pHMM from copper-dependent ApnU homologs has been curated, we recommend relying on the two HXXHC motifs, rather than homology-based annotation. The workflow can predict copper-dependent enzymes, but it is not suitable for distinguishing between halogenases and hydroxylases (SF) due to the lack of experimental data and close homology between the members of this family.

Known flavin-dependent monooxygenases and a dialkyldecalin synthase hit the general profile for FDHs with low bitscores. A phenol brominase, Bmp5 [EC: 1.14.19.55] from pentabromopseudilin biosynthesis [44] was not picked up during cross-validation, as this was the only representative of a single-component FDH with a GxGxxG, but without the conventional WxWxI and Fx.Px.Sx.G motif when building the general, family-specific FDH model. However, the final model does include Bmp5 and therefore detects homologous sequences. The general model can also detect other single-component FDHs (e.g. VemK [45], PezW [46]) with GxGxxG motif, but not sequences with GxGxxA motif even though the MSA used for building the model was cut after the flavin-binding site. For the general FDH profile, using cutoffs is not recommended and we suggest relying on the conserved motifs instead of the bitscore. The general model is suitable for identifying conventional members of the family with the Fx.Px.Sx.G and with or without the WxWxI motif.

Furthermore, there are substrate-specific models for tryptophan, pyrrolic and phenol acting halogenases. In the case of tryptophan-halogenases the models are suitable for differentiating between tryptophan 5 [EC: 1.14.19.58] and tryptophan-6/7 halogenases [1.14.19.59/1.14.19.9]. It is worth noting that while the tryptophan-5 profile does pick up MibH, a RiPP halogenase from the microbisporicin BGC [47], MibH-like sequences were underrepresented when building the model. Two separate pHMMs serve the differentiation between enzymes acting on tyrosine or orsellinic-acid like compounds. While these profiles work for the known sequences, the consideration of catalytic residues and conserved motifs are needed to make confident predictions.

The VHPO workflow consists of a general model and halide-acceptance specific models. These models follow the previously described naming conventions and do not predict halogen-preference. While the profiles set apart known or canonical sequences, as we will see later in the Sequence Similarity Networks (SSNs), there are unexplored spaces where a sequence hits multiple pHMMs. Furthermore, we try to give information about the selectivity of the enzyme if it matches the selective-chloroperoxidase profile, by searching for the catalytic Lys and Ser residues described in NapH1 [1.11.1.10] from napyradiomycin biosynthesis [40]. It also helps filter out NapH3-or MarH2-like sequences, which are close non-halogenase VHPO homologs 40].

Using pHMMs with the recommended cutoffs (Supplementary Table 1.) helps strict categorization but is only suitable for canonical enzymes and might exclude unique sequences that still show halogenation activity or the same substrate-specificity that the profiles e.g. for tryptophan or pyrrole FDHs indicate.

In contrast to that, the exploration of large families, like the FDHs and VHPOs, cutoffs help narrow down the option for candidates for experimental work.

### Workflow of the computational pipeline

The computational workflow categorizes the six mentioned enzyme families and the comparison of enzyme(s) of interest to known halogenases. It is set up as a literate programming tool, implemented in a python notebook to enable the user to upload their sequence data and guide them through the general categorization workflow (Figure 1).

**Figure 1.**
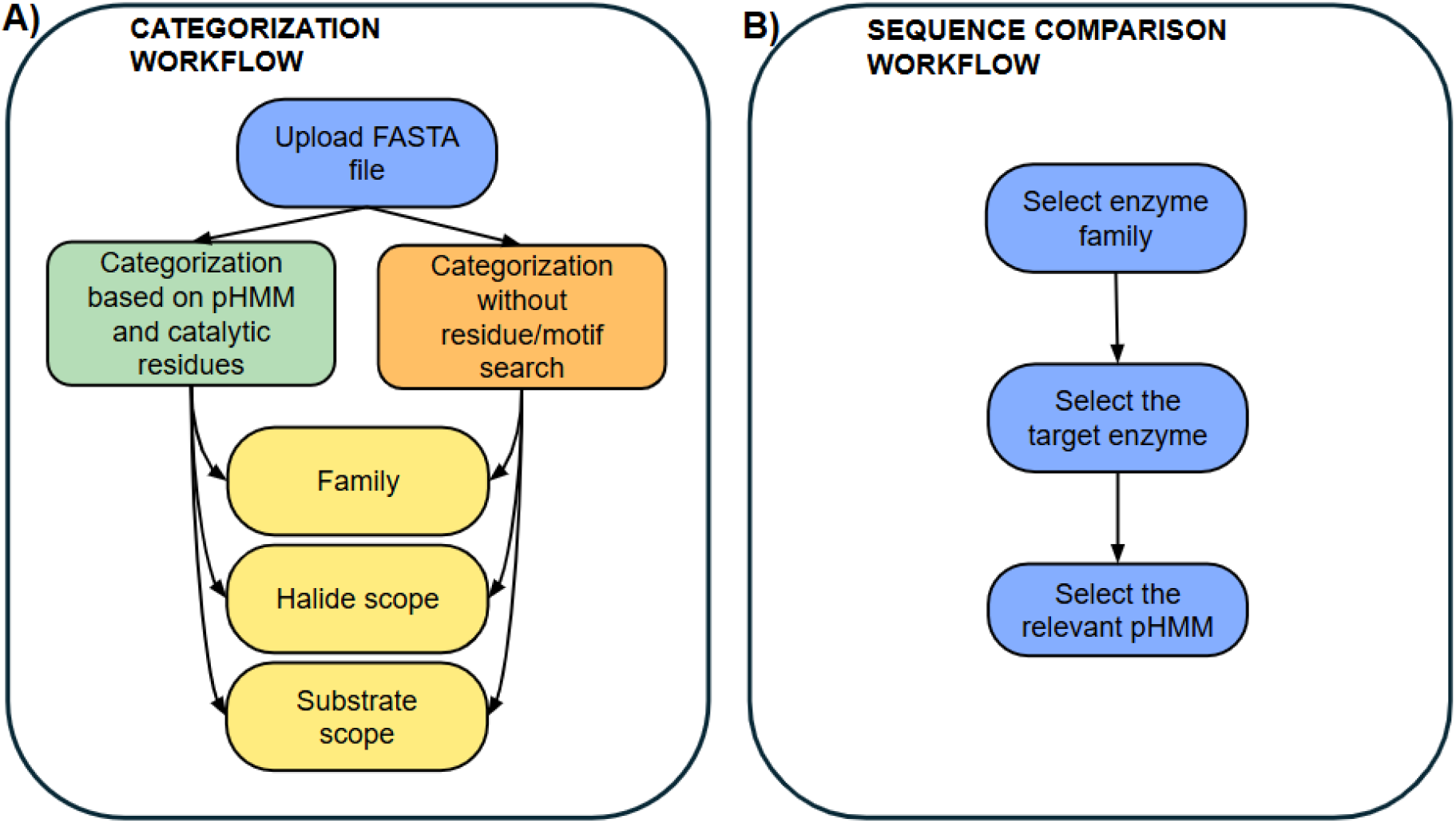
Process of sequence comparison. The workflow allows the comparison of target sequence(s) to known halogenases (**A**) and the search for validated or putative catalytic residues (**B**). To execute the comparison, the user needs to define the family of interest. After defining the enzyme family, the user chooses the target enzyme from a dropdown list. The database holds information on the given enzyme and the role of catalytic residues. The positions that the workflow searches for in the pHMM-alignment are pHMM-specific. Therefore, the user needs to choose the right pHMM for the comparison. To mine novel enzymes with a focus on known catalytic residues, the user chooses the enzyme family of interest. Each enzyme family has several options for known enzymes the user can compare their target sequence(s) to. For each enzyme the positions of catalytic residues in the pHMMs are collected. However, these positions are pHMM-specific. The user can choose the appropriate pHMM from a dropdown list. (Figure 1) The database holds information about what catalytic residue the position in the pHMM corresponds to and the role of that residue.

**Figure 2.**
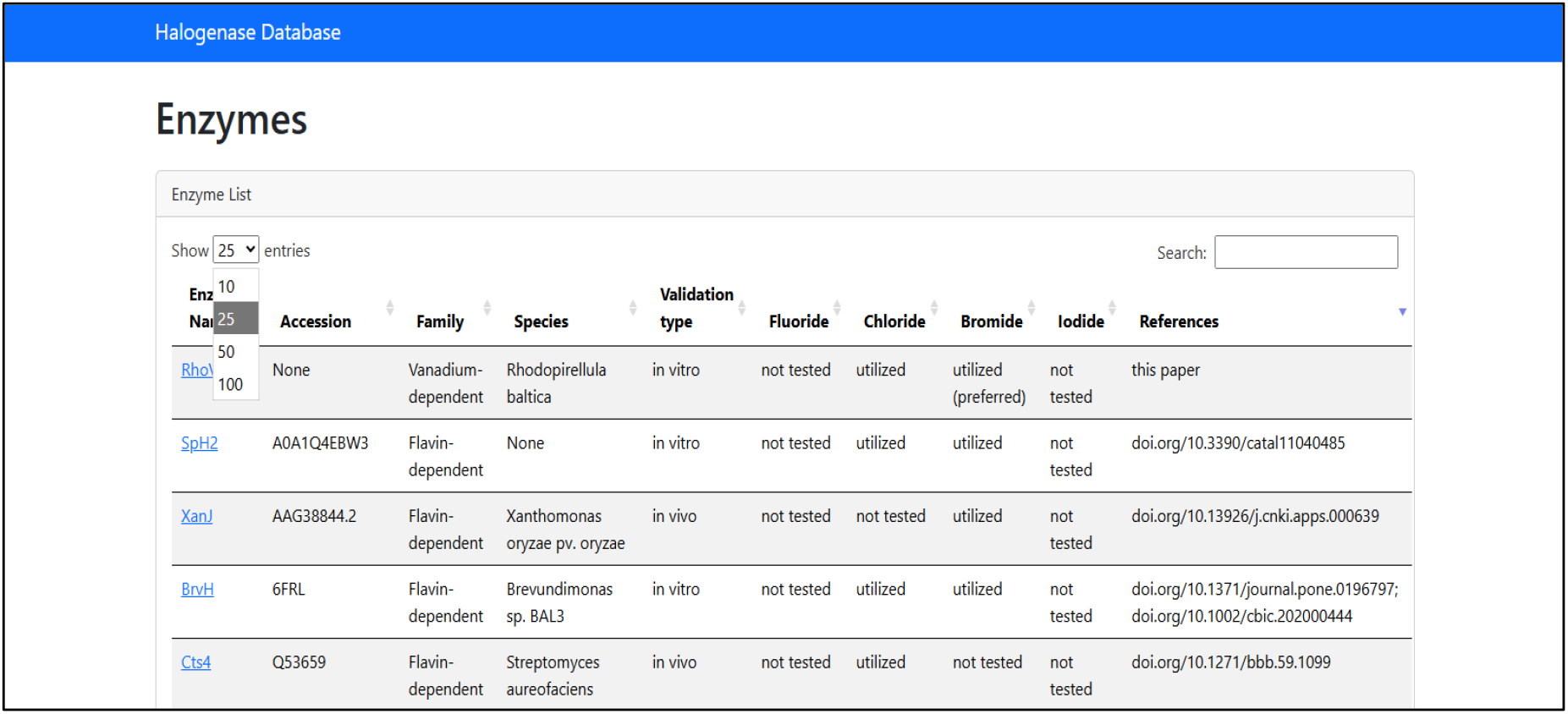
Halogenase DataBase. The first page of the database contains information on the name, accession which can be RefSeq ID, UniProt ID or PDB ID, the family the enzyme belongs to, the producing organism, its halide-acceptance and references to the studies of experimental validation, structure analysis and discovery.

**Figure 3.**
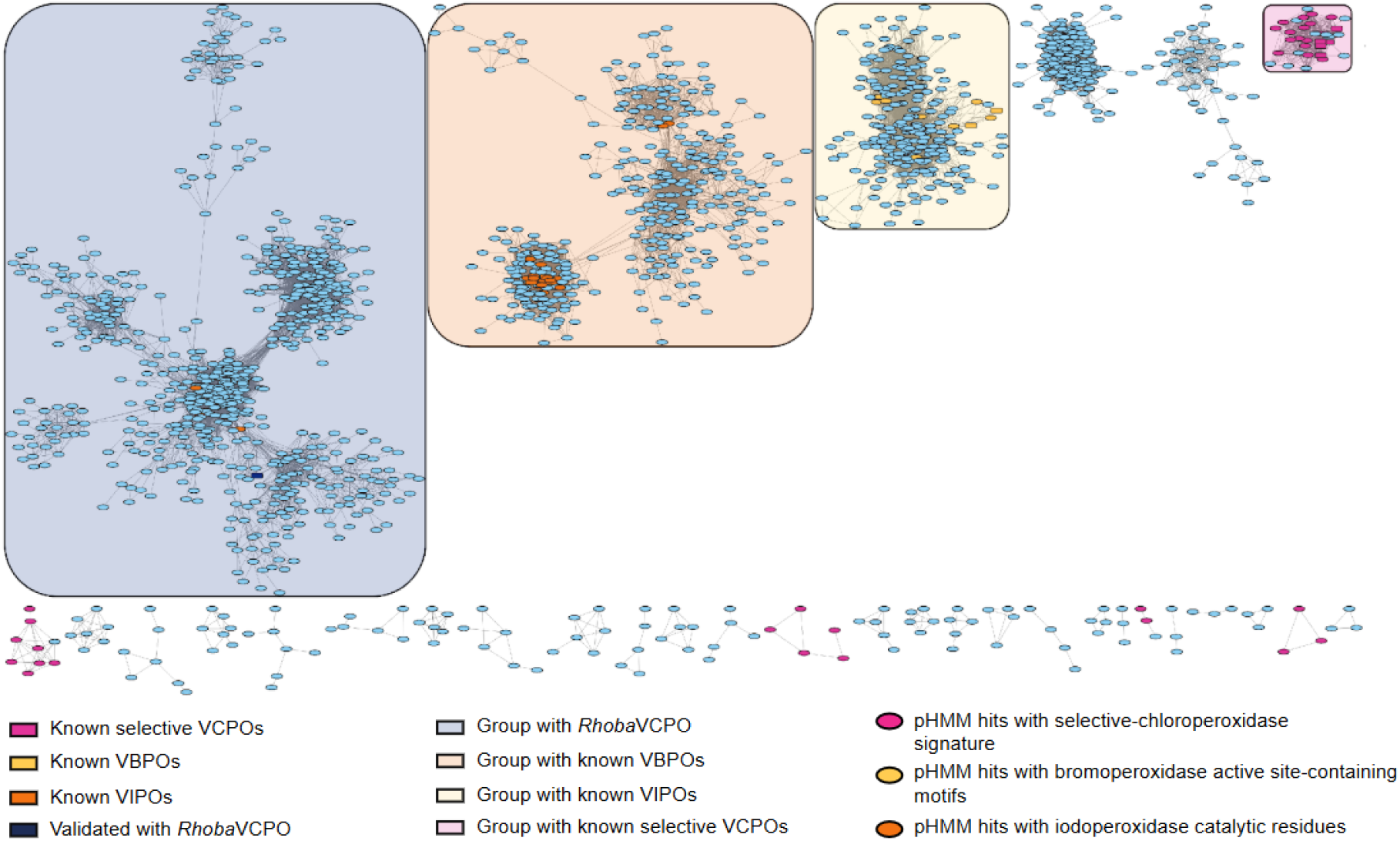
SSN of VHPOs. The built SSN contains hits from the selective-and non-selective chloroperoxidase pHMMs, as well as from the bromoperoxidase and iodoperoxidase pHMMs and reveals the space of VHPOs.

Users can upload FASTA file(s) with protein sequences. Following the upload, the name of the file that is to be analyzed needs to be written in the text box. After filling out the textbox and running the cell, the general categorization workflow can be executed.

The categorization for each family branches into two paths (Figure 1). It can be based on the presence of catalytic residues and conserved motifs or rely solely on the hit to certain pHMMs and the bitscore. The notebook showcases these two features. The recommended cutoffs are indicated in the notebook but are not applied during the workflow. The code will return all hits to the specific profile, and the user decides whether they consider the threshold. When the general workflow is finished, the user runs the family-specific tasks. These workflows differ in depth depending on the available experimental information about the family (SI).

### Computational workflow highlights novelty

The analysis of UniRef50 with the computational pipeline highlighted several unexplored groups across the covered halogenase families (SI) and the VHPO analysis caught our attention. Among the VHPOs, the known selective chloroperoxidases formed their own cluster that includes several sequences with the catalytic Lys and Ser residues. Unconnected to this cluster, smaller groups were revealed by the selective VCPO profile which also possesses these amino acids in the same column of the alignment. Two other clusters without the catalytic residues separate the VBPO cluster with the known sequences from the VCPOs. Close to them lies the VIPO cluster with the validated sequences and several undescribed sequences containing the same catalytic residues as the iodoperoxidase from *Zobellia galactivorans*. While they cluster tightly with the known sequences, they are connected to a separate part of the cluster that has no validated proteins yet. These catalytic residues are also present in two sequences in an unconnected group which contains a VHPO from *Rhodopirellula* (Q7UVW2), previously described as hypothetical VCPO [48] which clusters with VIPOs on the phylogenetic tree. Seeing the distance between this sequence from the known VCPOs and the presence of putative VIPOs made us question the predicted halide-acceptance of this sequence.

### VHPO from *Rhodopirellula* is a chlorinase with bromine preference

The SSN revealed the *Rhoba*VHPO in a cluster separate from the known chloroperoxidases. The positioning of the *Rhoba*VHPO was intriguing as it was indirectly connected to two sequences with the catalytic residues characteristic to iodoperoxidases. While *Rhoba*VHPO was previously annotated as a hypothetical chloroperoxidase, there was no experimental study to confirm this categorization or explore its halide scope. We conducted the monochlorodimedone (MCD) assay (Figure 4) to explore its chlorination and bromination potential (Figure 5).

**Figure 4.**
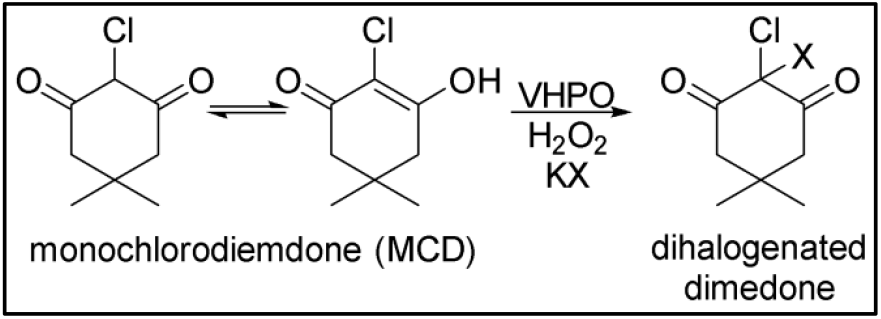
Reaction in the presence of KBr and KCl during MCDassay. Halogenation of MCD by VHPO in the presence of hydrogen peroxide and KX (X=Cl, Br).

**Figure 5.**
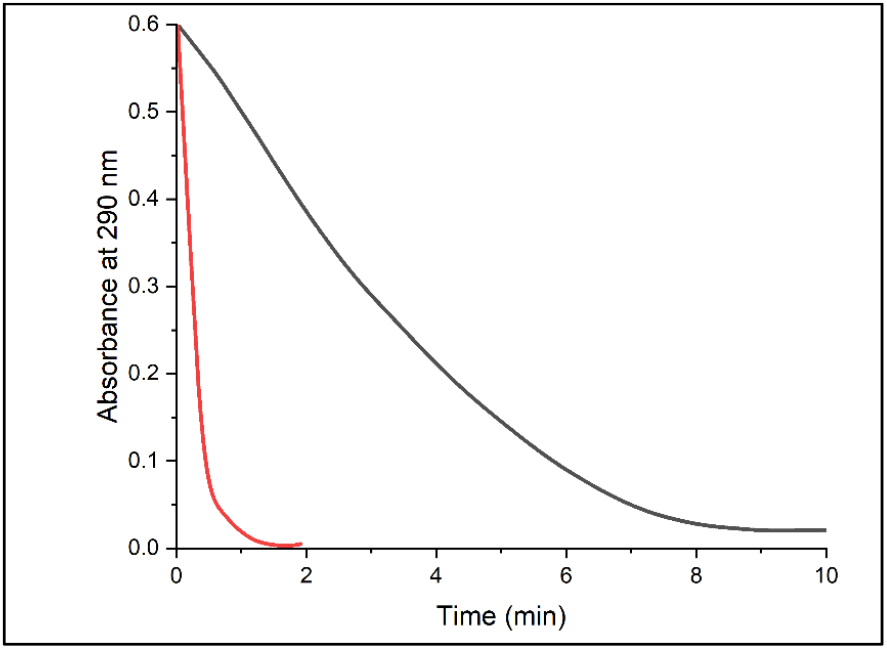
Absorbance trajectory at 290 nm in the presence of KBr and KCl. Absorbance of KCl (black) and KBr (red) at 290 nm.

Time course of MCD (µM) halogenation catalysed by *Rhoba*VHPO (0.5 uM) in the presence of KBr (200 mM, red) and KCl (200 mM, black), Na_3_VO_4_ (10 uM) in 50 mM MES buffer pH=6. The reactions were initiated by addition of H_2_O_2_ (1 mM). Reactions were monitored by decrease in absorbance at 290 nm. The rate of reaction was faster in the presence of KBr than KCl (Figure 5).

The MCD assay confirmed the chlorination activity and showed around five-fold stronger preference for bromide.

## CONCLUSION

This study provides a user-friendly and transparent way of detecting and categorizing halogenases and mining for novel enzymes across the SAM-dependent, NHFe-dependent, dimetal-carboxylate, copper-dependent, flavin-dependent and vanadium-dependent families. Currently known enzymes are collected in a database with information on halide-acceptance, mutagenesis studies and structure analysis as well as the positions of catalytic residues in the pHMMs.

Although catalytic residues strengthen the annotation, search for small motifs e.g. HXA/G in NHFe-dependent homologs can result in false positives. This problem mitigated by restricting the search area in the pHMM alignment. However, it raises another issue. Sequences with deletions or insertions around the searched sequence regions may be discarded by the workflow. This tradeoff should be weighed by the user. Therefore, using recommended cutoffs is optional and the user can view hits with and without the catalytic residues.

Moreover, running the general workflow on UniRef50 helped us detect several promising candidates for halogenation, supported by the presence of conserved motifs and catalytic residues. The SSNs show underexplored clusters across all discussed enzyme families containing putative halogenases. Furthermore, we confirmed the chlorinating activity of the *Rhoba*VHPO which guides us in the exploration of the cluster holding other putative chloroperoxidases and potential iodinases. Additional experimental and structural studies are needed to determine the biological function *Rhoba*VHPO and what drives the low sequence identity with other chloroperoxidases.

## MATERIALS AND METHODS

### Halogenase Data Base (DB)

Literature search was carried out to extract information on halogenases across the SAM-dependent, NHFe-dependent, dimetal-carboxylate, FDH, and VHPO families. We developed a database in PostgreSQL to store the results of the literature search, which was converted to JSON for easier accessibility. The database stores data about enzyme name, accession/availability, producing organism, type of validation, halide-acceptance, and references. If information on mutagenesis and structure analysis studies were available, it was deposited in a table specific to that enzyme. Minimal information about the protocol was also noted.

We collected the so far experimentally validated or *in silico* inferred catalytic residues in the database to aid proper annotation. The database includes the catalytic residues and their roles, as well as how the mutation of those residues affected enzyme activity in mutagenesis studies.

HMMEditor [48] was used to determine the index of the catalytic residue in the given pHMM.

### pHMM curation and testing

pHMMs were built from Multiple Sequence Alignments (MSA). For FDHs, the sequences were often trimmed at the N-and C-terminal regions if they showed high variability. As the low-homology regions at the terminals were excluded, MUSCLE was suitable for MSA generation. In case of the SAM-dependent and metal-dependent families, the pHMMs were often built from whole sequence alignments (Supplementary Table 1.). Therefore, we used MAFFT for SAM-dependent, NHFe-dependent, dimetal-carboxylate halogenases and VHPOs, an alignment algorithm which is more suitable to handle low homology N-terminal or C-terminal extensions [49, 50]

Sequences were retrieved from the study which described them, UniProt or NCBI [51].

### SSN curation

UniRef50 was searched with hmmer [52] against the curated pHMMs without thresholds. We added the known enzymes used for building the pHMMs to each set of sequences. The resulting FASTA files were the input for the Enzyme Function Initiative-Enzyme Similarity Tool (EFI-EST) [53, 54] to generate the SSNs. We visualized the generated SSN with Cytoscape 3.10.1 [55] with the default E-value for edge calculation (negative log of 5). The Sequence Alignment threshold parameter in EFI-EST [53,54] was modified with consideration to the relationship between the value of the Sequence Alignment score and the percent identity. The cutoff was chosen at a value where enzymes with similar halide-or substrate-scope clustered together and were separated from their closest homologs with different activity.

Geneious Prime 2024.03.25 was used to retrieve the domain composition data of the sequences.

### MCD Assay of VHPO *Rhodopirellula baltica*

All chemicals and biological materials were obtained from commercial suppliers (SI). The assay followed the protocol described by McKinnie, Miles and Moore [55].

### Construction of pET28a(+)_*Rhoba*VHPO

The *E. coli*-optimised gene (synthesised by Twist) for VHPOs from *Rhodopirellula baltica* was cloned into pET28a(+) using *NdeI* and *XhoI* restriction sites to yield pET28a(+)_VHPO_Rhoba.

### Protein production and purification

For expression of *Rhoba*VHPO, chemically competent *E. coli* BL21 (DE3) were transformed with the relevant pET28a(+)_VHPO construct. Single colonies of freshly transformed cells were cultured for 18 h in 5 mL LB medium containing 50 µg mL^−1^ kanamycin. Starter cultures (5 mL) were used to inoculate 500 mL TB medium supplemented with 50 µg mL^−1^ kanamycin. Cultures grew at 37 °C, 200 rpm to an optical density at 600 nm (OD600) of around 0.5. Protein expression was induced with the addition of IPTG to a final concentration of 0.1 mM. Induced cultures were incubated for 20 h at 18 °C and the cells were subsequently collected by centrifugation (4,000 rpm for 30 min). Pelleted cells were resuspended in a lysis buffer (50 mM HEPES, 300 mM NaCl, pH 7.5 containing 20 mM imidazole) and lysed by sonication. Cell lysates were cleared by centrifugation (10,000 rpm for 30 min) and supernatants were subjected to affinity chromatography using Ni-NTA Agarose (Qiagen). Purified protein was eluted using 50 mM HEPES, 300 mM NaCl, pH 7.5 containing 250 mM imidazole. The protein was desalted using 10DG desalting columns (Bio-Rad) with 50 mM MES pH 6 and analyzed by SDS PAGE. The protein was aliquoted, flash-frozen in liquid nitrogen and stored at −80 °C. Protein concentration was determined by measuring the absorbance at 280 nm and assuming an extinction coefficient of 76,320 M^−1^ cm^−1^.

### Monochlorodimedone (MCD) assay

Assays were performed in a quartz cuvette using a Cary 50 Bio UV-vis spectrophotometer measuring absorbance at 290 nm. MCD (50 μM), KCl or KBr (200 mM) Na_3_VO_4_ (10 uM) and purified *Rhoba*VHPO (0.5 μM) in 50 mM MES pH 6 were mixed in a cuvette and measured for 1 minute. After 1 minute the reaction was initiated by addition of H_2_O_2_ (1 mM). The total reaction volume was 1 mL. Data was processed and plotted in OriginPro 2025.

## SUPPLEMENTARY INFORMATION

ppt file with the SSN parameters, metric analysis and figures xxgml files for the SSNs

## DATA AVAILABILITY

Availability of the database:

Notebook: https://colab.research.google.com/drive/1imau-B7BDzHzTjVrzo6vXCOxtDUcKnq7?usp=sharing

Hal-miner github: https://github.com/JudSze/hal_miner

Halogenase DB: https://halogenases.secondarymetabolites.org/

Github repository of the pHMM curation, cross-validation and testing: https://github.com/JudSze/hal_miner_curation

## ACKNOWLEDGEMENTS

We thank Sanjoy Adak for guidance with the chloroperoxidase group and Vinayak Agarwal for providing the SrpI tryptophan-6 brominase sequence. Special thanks to Thom Booth for suggesting tryptophan 7-halogenases as starting point of this work.

This project has received funding from the European Union’s Horizon Europe program under the Marie Skłodowska-Curie grant agreement No 101072485. A. L. received funding from FCT PhD scholarship 2020.08183.BD. A. B. acknowledges funding through Scripps Institutional Postdoctoral Program. P.N.L. acknowledges funding through an ERC Consolidator Grant No 101088806. T.W. and K.B. also received funding from the Novo Nordisk Foundation No NNF20CC0035580. We also acknowledge support from the National Institute of General Medical Sciences (1R35GM159745 to A.L.L. and 2R01GM085770 to B.S.M.). P.I.N. acknowledges funding from the European Union’s Horizon 2020 Research and Innovation Programme under grant agreement No. 101082049 (TOLERATE), and by the The Novo Nordisk Foundation through grants NNF20CC0035580, LiFe (NNF18OC0034818), TARGET (NNF21OC0067996), FM·Pseudomonas (NNF24OC0091501), and NovoF (NNF23OC0083631).

## REFERENCES

1. Atashgahi, S., Häggblom, M. M., & Smidt, H. (2018). Organohalide respiration in pristine environments: implications for the natural halogen cycle. Environmental microbiology, 20(3), 934–948.

2. Zlamal, Jaime E., Theodore K. Raab, Mark Little, Robert A. Edwards, and David A. Lipson. 2017. “Biological Chlorine Cycling in the Arctic Coastal Plain.” Biogeochemistry 134 (3): 243–60.

3. Crowe, Charlotte, Samuel Molyneux, Sunil V. Sharma, Ying Zhang, Danai S. Gkotsi, Helen Connaris, and Rebecca J. M. Goss. 2021. “Halogenases: A Palette of Emerging Opportunities for Synthetic Biology-Synthetic Chemistry and C-H Functionalisation.” Chemical Society Reviews 50 (17): 9443–81.

4. Agarwal, V., Miles, Z. D., Winter, J. M., Eustáquio, A. S., El Gamal, A. A., & Moore, B. S. (2017). Enzymatic halogenation and dehalogenation reactions: pervasive and mechanistically diverse. Chemical reviews, 117(8), 5619–5674.

5. Cochereau, B., Meslet-Cladière, L., Pouchus, Y. F., Grovel, O., & Roullier, C. (2022). Halogenation in fungi: what do we know and what remains to be discovered?. Molecules, 27(10), 3157.

6. H.M. Berman, J. Westbrook, Z. Feng, G. Gilliland, T.N. Bhat, H. Weissig, I.N. Shindyalov, P.E. Bourne, The Protein Data Bank (2000) Nucleic Acids Research 28: 235–242 10.1093/nar/28.1.235. (RCSB.org)

7. Prakinee, K., Jitkaroon, W., & Chaiyen, P. (2025). Flavin-Dependent Halogenases: Emerging Enzymes and New Functions. ChemCatChem, e00558.

8. Adak, S., Lukowski, A. L., Schäfer, R. J., & Moore, B. S. (2022). From tryptophan to toxin: nature’s convergent biosynthetic strategy to aetokthonotoxin. Journal of the American Chemical Society, 144(7), 2861–2866.

9. Lukowski, A. L., Hubert, F. M., Ngo, T. E., Avalon, N. E., Gerwick, W. H., & Moore, B. S. (2023). Enzymatic halogenation of terminal alkynes. Journal of the American Chemical Society, 145(34), 18716–18721.

10. Phintha, A., Prakinee, K., & Chaiyen, P. (2020). Structures, mechanisms and applications of flavin-dependent halogenases. The Enzymes, 47, 327–364.

11. Minges, H., & Sewald, N. (2020). Recent advances in synthetic application and engineering of halogenases. ChemCatChem, 12(18), 4450–4470.

12. Podzelinska, K., Latimer, R., Bhattacharya, A., Vining, L. C., Zechel, D. L., & Jia, Z. (2010). Chloramphenicol biosynthesis: the structure of CmlS, a flavin-dependent halogenase showing a covalent flavin–aspartate bond. Journal of molecular biology, 397(1), 316–331.

13. Liu, M., Ohashi, M., Hung, Y. S., Scherlach, K., Watanabe, K., Hertweck, C., & Tang, Y. (2021). AoiQ catalyzes geminal dichlorination of 1, 3-diketone natural products. Journal of the American Chemical Society, 143(19), 7267–7271.

14. Chankhamjon, P., Tsunematsu, Y., Ishida-Ito, M., Sasa, Y., Meyer, F., Boettger-Schmidt, D., … & Hertweck, C. (2016). Regioselective Dichlorination of a non-activated Aliphatic Carbon Atom and Phenolic Bismethylation by a multifunctional fungal flavoenzyme. Angewandte Chemie International Edition, 55(39), 11955–11959.

15. Martinez, J. S., Carroll, G. L., Tschirret-Guth, R. A., Altenhoff, G., Little, R. D., & Butler, A. (2001). On the regiospecificity of vanadium bromoperoxidase. Journal of the American Chemical Society, 123(14), 3289–3294.

16. Baumgartner, J. T., & McKinnie, S. M. (2024). Regioselective Halogenation of Lavanducyanin by a Site-Selective Vanadium-Dependent Chloroperoxidase. Organic Letters, 26(27), 5725–5730.

17. Deng, H., O’Hagan, D., & Schaffrath, C. (2004). Fluorometabolite biosynthesis and the fluorinase from Streptomyces cattleya. Natural product reports, 21(6), 773–784.

18. Schaffrath, C., Deng, H., & O’Hagan, D. (2003). Isolation and characterisation of 5′-fluorodeoxyadenosine synthase, a fluorination enzyme from Streptomyces cattleya. FEBS letters, 547(1-3), 111–114.

19. Pardo, I., Bednar, D., Calero, P., Volke, D. C., Damborsky, J., & Nikel, P. I. (2022). A nonconventional Archaeal fluorinase identified by in silico mining for enhanced fluorine biocatalysis. ACS catalysis, 12(11), 6570–6577.

20. Hillwig, M. L., & Liu, X. (2014). A new family of iron-dependent halogenases acts on freestanding substrates. Nature chemical biology, 10(11), 921–923.

21. Ni, J., Zhuang, J., Shi, Y., Chiang, Y. C., & Cheng, G. J. (2024). Discovery and substrate specificity engineering of nucleotide halogenases. Nature Communications, 15(1), 5254.

22. Vaillancourt, F. H., Yin, J., & Walsh, C. T. (2005). SyrB2 in syringomycin E biosynthesis is a nonheme FeII α-ketoglutarate-and O2-dependent halogenase. Proceedings of the National Academy of Sciences, 102(29), 10111–10116.

23. Eusebio, N., Rego, A., Glasser, N. R., Castelo-Branco, R., Balskus, E. P., & Leão, P. N. (2021). Distribution and diversity of dimetal-carboxylate halogenases in cyanobacteria. BMC genomics, 22(1), 633.

24. Pawlik-Skowrońska, B., Bownik, A., Pogorzelec, M., Kulczycka, J., & Sumińska, A. (2023). First report on adverse effects of cyanobacterial anabaenopeptins, aeruginosins, microginin and their mixtures with microcystin and cylindrospermopsin on aquatic plant physiology: An experimental approach. Toxicon, 236, 107333.

25. Leão, P. N., Martins, T. P., Abt, K., Reis, J. P., Figueiredo, S., Castelo-Branco, R., & Freitas, S. (2023). Incorporation and modification of fatty acids in cyanobacterial natural products biosynthesis. Chemical Communications, 59(30), 4436–4446.

26. Ramaswamy, A. V., Sorrels, C. M., & Gerwick, W. H. (2007). Cloning and biochemical characterization of the hectochlorin biosynthetic gene cluster from the marine cyanobacterium Lyngbya majuscula. Journal of Natural Products, 70(12), 1977–1986.

27. Pratter, S., & Straganz, G. D. (2010). Non-heme Fe (II) Enzymes: First Insights into Fatty Acid Halogenation by the Cyanobacterial Enzyme HctB. In Pacifichem 2010.

28. Chiang, C. Y., Ohashi, M., Le, J., Chen, P. P., Zhou, Q., Qu, S., … & Tang, Y. (2025). Copper-dependent halogenase catalyses unactivated C− H bond functionalization. Nature, 638(8049), 126–132.

29. Hon, J., Borko, S., Stourac, J., Prokop, Z., Zendulka, J., Bednar, D., … & Damborsky, J. (2020). EnzymeMiner: automated mining of soluble enzymes with diverse structures, catalytic properties and stabilities. Nucleic acids research, 48(W1), W104–W109.

30. Altschul, S. F., Gish, W., Miller, W., Myers, E. W., & Lipman, D. J. (1990). Basic local alignment search tool. Journal of molecular biology, 215(3), 403–410.

31. Eddy, S. R. (1998). Profile hidden Markov models. Bioinformatics (Oxford, England), 14(9), 755–763.

32. Söding, J., Biegert, A., & Lupas, A. N. (2005). The HHpred interactive server for protein homology detection and structure prediction. Nucleic acids research, 33(suppl_2), W244–W248.

33. Blum, M., Chang, H. Y., Chuguransky, S., Grego, T., Kandasaamy, S., Mitchell, A., … & Finn, R. D. (2021). The InterPro protein families and domains database: 20 years on. Nucleic acids research, 49(D1), D344–D354.

34. Jeon, J., Lee, J., Jung, S. M., Shin, J. H., Song, W. J., & Rho, M. (2021). Genomic determinants encode the reactivity and regioselectivity of flavin-dependent halogenases in bacterial genomes and metagenomes. Msystems, 6(3), 10–1128.

35. Ahmad, S., Jose da Costa Gonzales, L., Bowler-Barnett, E. H., Rice, D. L., Kim, M., Wijerathne, S., … & Martin, M. J. (2025). The UniProt website API: facilitating programmatic access to protein knowledge. Nucleic Acids Research, gkaf394.

36. Frank, A., Seel, C. J., Groll, M., & Gulder, T. (2016). Characterization of a cyanobacterial haloperoxidase and evaluation of its biocatalytic halogenation potential. ChemBioChem, 17(21), 2028–2032.

37. Soedjak, H. S., & Butler, A. (1990). Characterization of vanadium bromoperoxidase from Macrocystis and Fucus: reactivity of vanadium bromoperoxidase toward acyl and alkyl peroxides and bromination of amines. Biochemistry, 29(34), 7974–7981.

38. Frank, A., Seel, C. J., Groll, M., & Gulder, T. (2016). Characterization of a cyanobacterial haloperoxidase and evaluation of its biocatalytic halogenation potential. ChemBioChem, 17(21), 2028–2032.

39. Soedjak, H. S., & Butler, A. (1990). Characterization of vanadium bromoperoxidase from Macrocystis and Fucus: reactivity of vanadium bromoperoxidase toward acyl and alkyl peroxides and bromination of amines. Biochemistry, 29(34), 7974–7981.

40. Chen, P. Y. T., Adak, S., Chekan, J. R., Liscombe, D. K., Miyanaga, A., Bernhardt, P., … & Moore, B. S. (2022). Structural basis of stereospecific vanadium-dependent haloperoxidase family enzymes in napyradiomycin biosynthesis. Biochemistry, 61(17), 1844–1852.

41. Neugebauer, M. E., Sumida, K. H., Pelton, J. G., McMurry, J. L., Marchand, J. A., & Chang, M. C. (2019). A family of radical halogenases for the engineering of amino-acid-based products. Nature chemical biology, 15(10), 1009–1016.

42. Wang, M. L., Glasser, N. R., Nair, M. A., Krebs, C., Martin Bollinger Jr, J., & Balskus, E. P. (2025). Biochemical Studies of a Cyanobacterial Halogenase Support the Involvement of a Dimetal Cofactor. Biochemistry, 64(10), 2173–2180.

43. Eustáquio, A. S., Pojer, F., Noel, J. P., & Moore, B. S. (2008). Discovery and characterization of a marine bacterial SAM-dependent chlorinase. Nature Chemical Biology, 4(1), 69–74.

44. Agarwal, V., El Gamal, A. A., Yamanaka, K., Poth, D., Kersten, R. D., Schorn, M., … & Moore, B. S. (2014). Biosynthesis of polybrominated aromatic organic compounds by marine bacteria. Nature chemical biology, 10(8), 640–647.

45. Song, R., Shi, H., Zhu, J., Wang, H., & Shen, Y. (2019). A single-component flavoenzyme catalyzed regioselective halogenation of pyrone in the biosynthesis of venemycins. ACS Chemical Biology, 14(12), 2533–2537.

46. Chen, Z., Duan, H., Zhao, M., Xiao, F., & Li, W. (2025). Discovery and Characterization of a Single-Component Halogenase for Phenazine Halogenation. Organic Letters, 27(37), 10553–10558.

47. Foulston, L. C., & Bibb, M. J. (2010). Microbisporicin gene cluster reveals unusual features of lantibiotic biosynthesis in actinomycetes. Proceedings of the National Academy of Sciences, 107(30), 13461–13466.

48. Winter, J. M., & Moore, B. S. (2009). Exploring the chemistry and biology of vanadium-dependent haloperoxidases. Journal of Biological Chemistry, 284(28), 18577–18581.

49. Katoh, Kazutaka, Kazuharu Misawa, Kei-Ichi Kuma, and Takashi Miyata. 2002. “MAFFT: A Novel Method for Rapid Multiple Sequence Alignment Based on Fast Fourier Transform.” Nucleic Acids Research 30 (14): 3059–66.

50. Edgar, Robert C. 2004. “MUSCLE: Multiple Sequence Alignment with High Accuracy and High Throughput.” Nucleic Acids Research 32 (5): 1792–97.

51. Sayers, E. W., Beck, J., Bolton, E. E., Brister, J. R., Chan, J., Connor, R., … & Pruitt, K. D. (2025). Database resources of the National Center for Biotechnology Information in 2025. Nucleic acids research, 53(D1), D20–D29.

52. Eddy, S. (1992). HMMER user’s guide. Department of Genetics, Washington University School of Medicine, 2(1), 13.

53. Gerlt, J. A., Bouvier, J. T., Davidson, D. B., Imker, H. J., Sadkhin, B., Slater, D. R., & Whalen, K. L. (2015). Enzyme function initiative-enzyme similarity tool (EFI-EST): a web tool for generating protein sequence similarity networks. Biochimica Et Biophysica Acta (BBA)-Proteins and Proteomics, 1854(8), 1019–1037.

54. Oberg, N., Zallot, R., & Gerlt, J. A. (2023). EFI-EST, EFI-GNT, and EFI-CGFP: enzyme function initiative (EFI) web resource for genomic enzymology tools. Journal of molecular biology, 435(14), 168018.

55. Shannon, P., Markiel, A., Ozier, O., Baliga, N. S., Wang, J. T., Ramage, D., … & Ideker, T. (2003). Cytoscape: a software environment for integrated models of biomolecular interaction networks. Genome research, 13(11), 2498–2504.

56. McKinnie, Shaun M. K., Zachary D. Miles, and Bradley S. Moore. 2018. “Characterization and Biochemical Assays of Streptomyces Vanadium-Dependent Chloroperoxidases.” Methods in Enzymology 604 (April): 405–24.

