## Supplemental Figures for "Computational pipeline reveals nature’s untapped reservoir of halogenating enzymes"

### Monochlorodimedone (MCD) assay

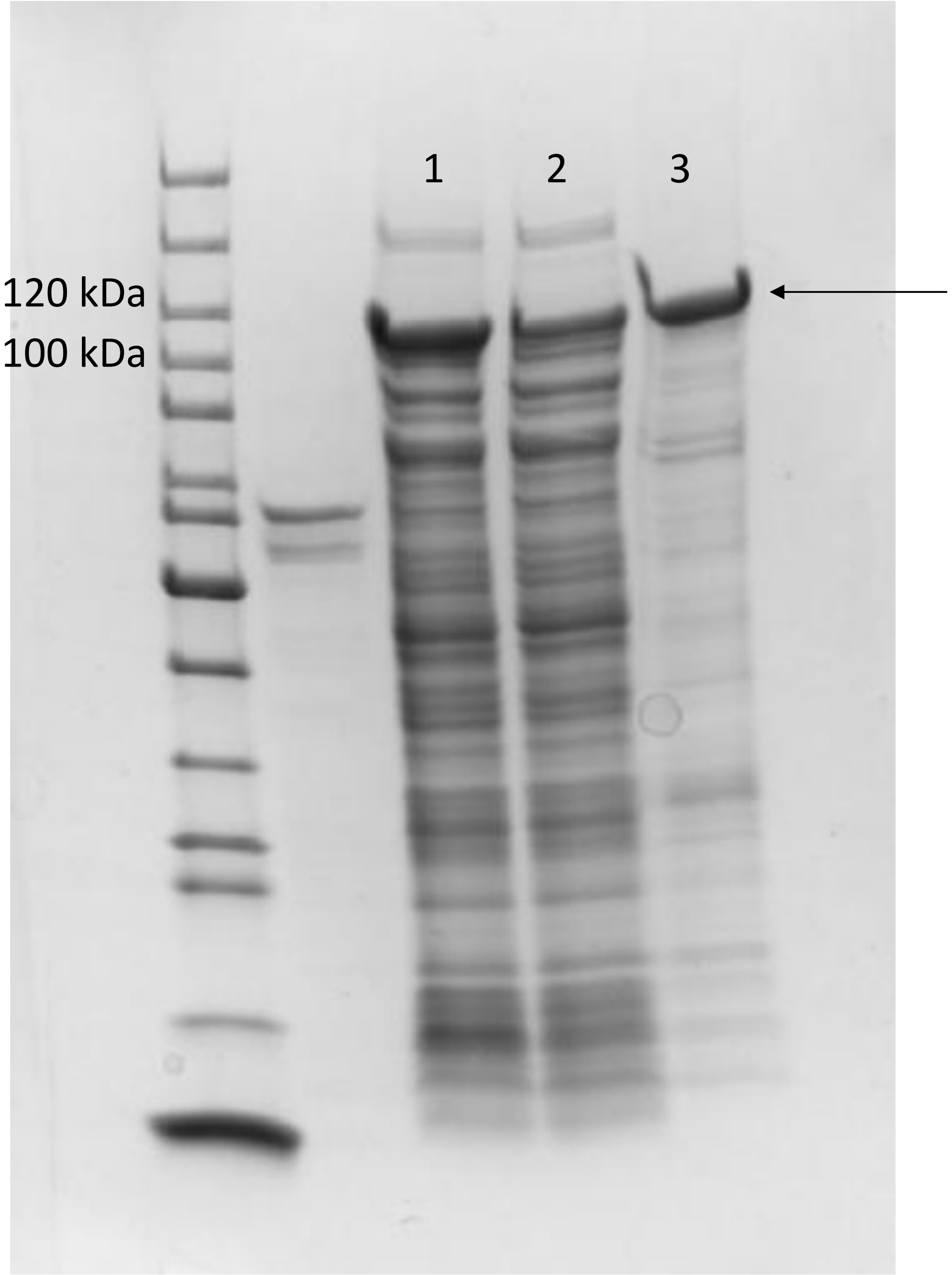


1- Total proteins
2- Soluble proteins
3- Purified fraction

Expected size of his-tagged protein 78.7 kDa

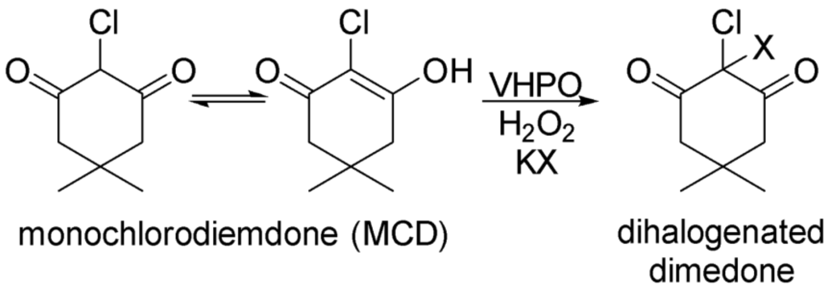


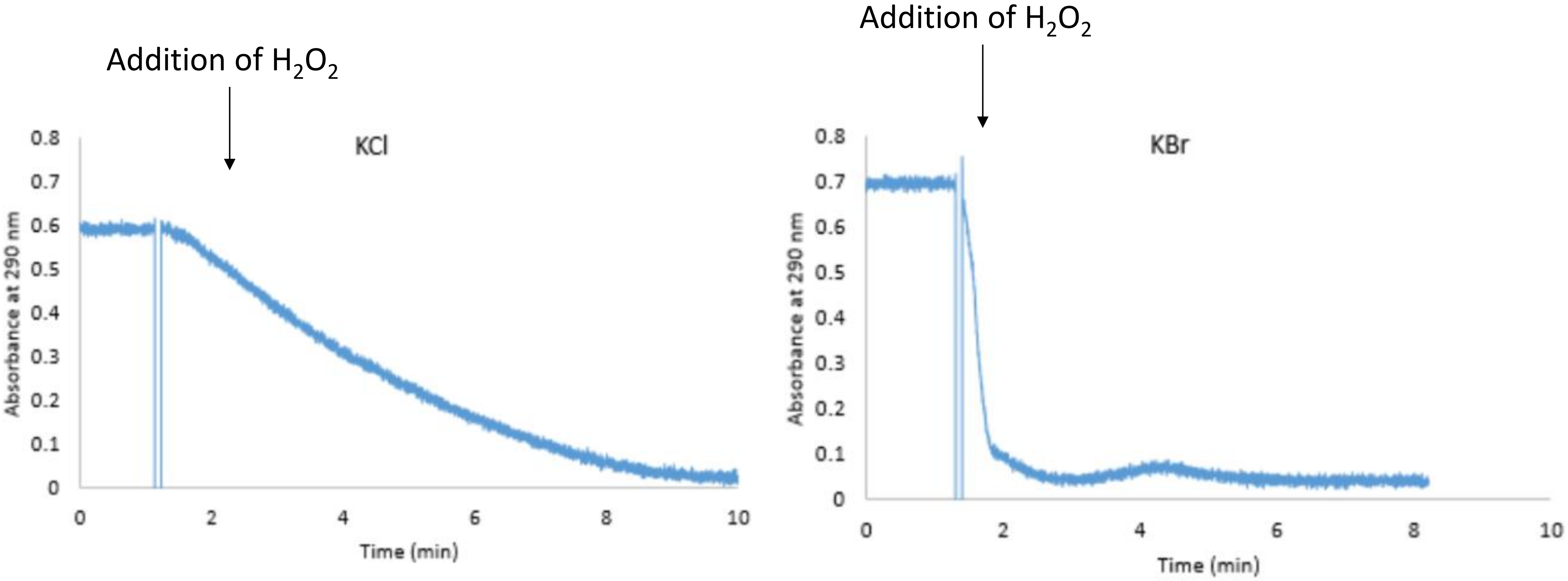


Assay conditions: 50 uM MCD, 200 mM KBr or KCl, 10 uM $Na_3VO_4$, 0.5 uM Enzyme, Buffer: 50 mM MES pH=6. Initiate reaction with 1 mM $H_2O_2$.

### Sequence Similarity Networks

### Vanadium-dependent

| | |
|---|---|
| Job Number | 156836 |
| Time Started -- Finished | 7/2 06:54 AM -- 7/2 07:00 AM |
| Database Version | UniProt: 2025-02 / InterPro: 105 |
| Input Option | FASTA (Option C), no FASTA header reading |
| Job Name | VHPOs |
| Input Sequence Source | UniRef50 |
| E-Value for SSN Edge Calculation | 5 |
| Uploaded FASTA File | ssn.fasta |
| Number of Sequences in Uploaded File | 3,654 |
| Exclude Fragments | No |
| Total Number of Sequences in Dataset | 3,654 |
| Total Number of Edges | 1,640,171 |
| Number of Unique Sequences | 3,650 |
| Convergence Ratio? | 0.246 |

Alignment Score threshold: 80

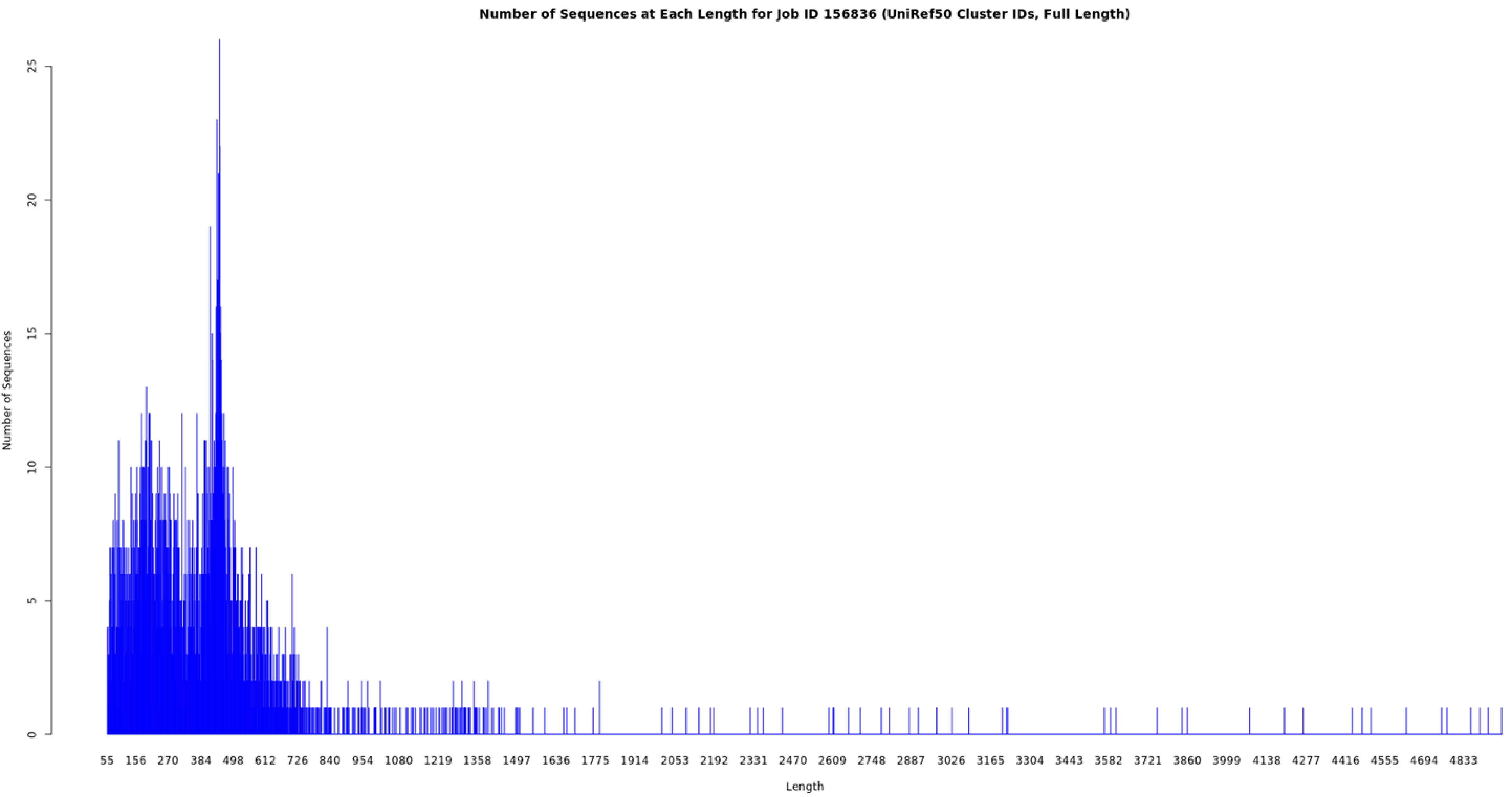
Number of Sequences at Each Length for Job ID 156836 (UniRef50 Cluster IDs, Full Length)
Number of Sequences
0
5
10
15
20
25
55 156 270 384 498 612 726 840 954 1080 1219 1358 1497 1636 1775 1914 2053 2192 2331 2470 2609 2748 2887 3026 3165 3304 3443 3582 3721 3860 3999 4138 4277 4416 4555 4694 4833
Length

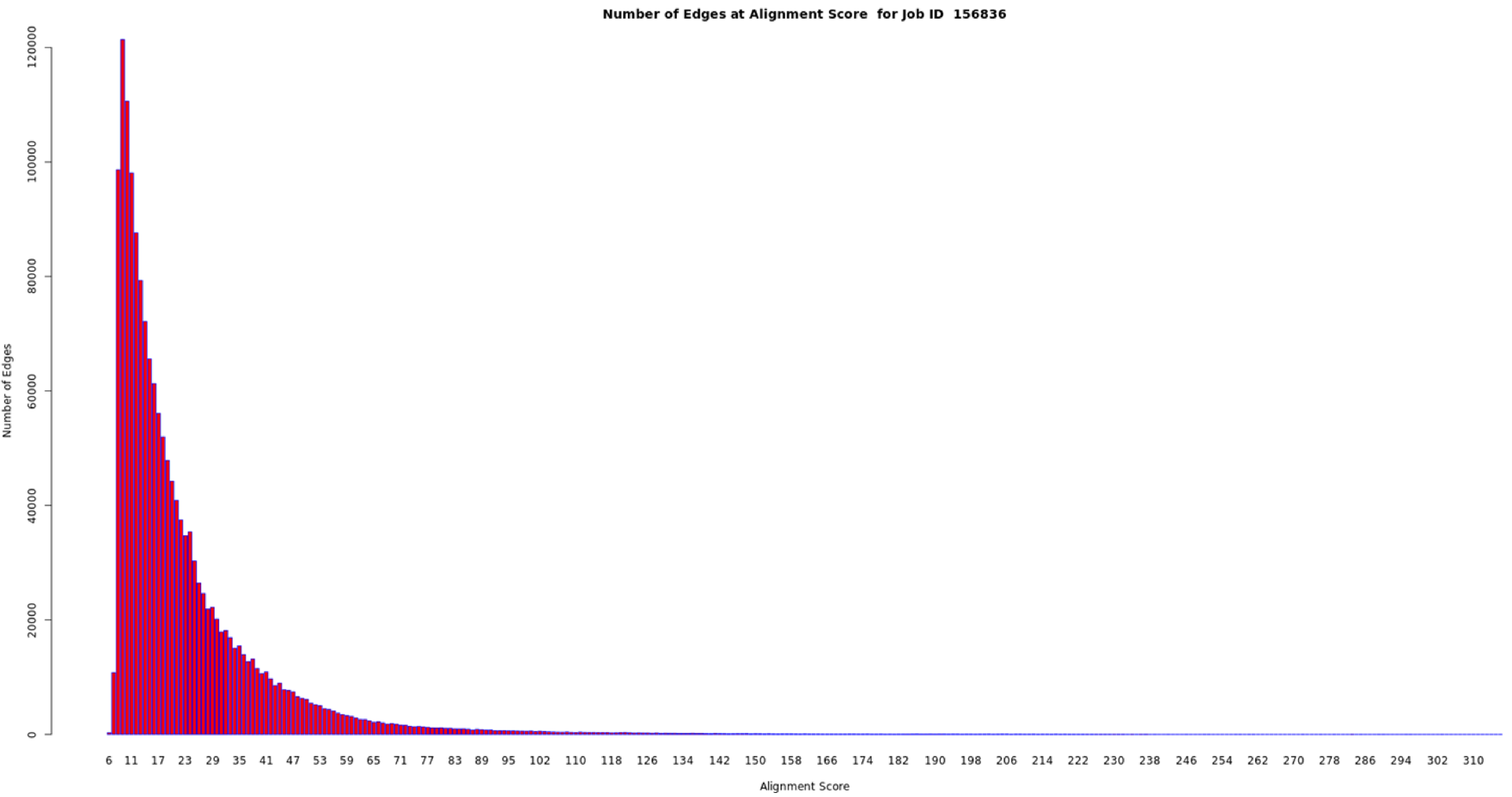

Number of Edges at Alignment Score for Job ID 156836
Number of Edges
120000
100000
80000
60000
40000
20000
0
6 11 17 23 29 35 41 47 53 59 65 71 77 83 89 95 102 110 118 126 134 142 150 158 166 174 182 190 198 206 214 222 230 238 246 254 262 270 278 286 294 302 310
Alignment Score

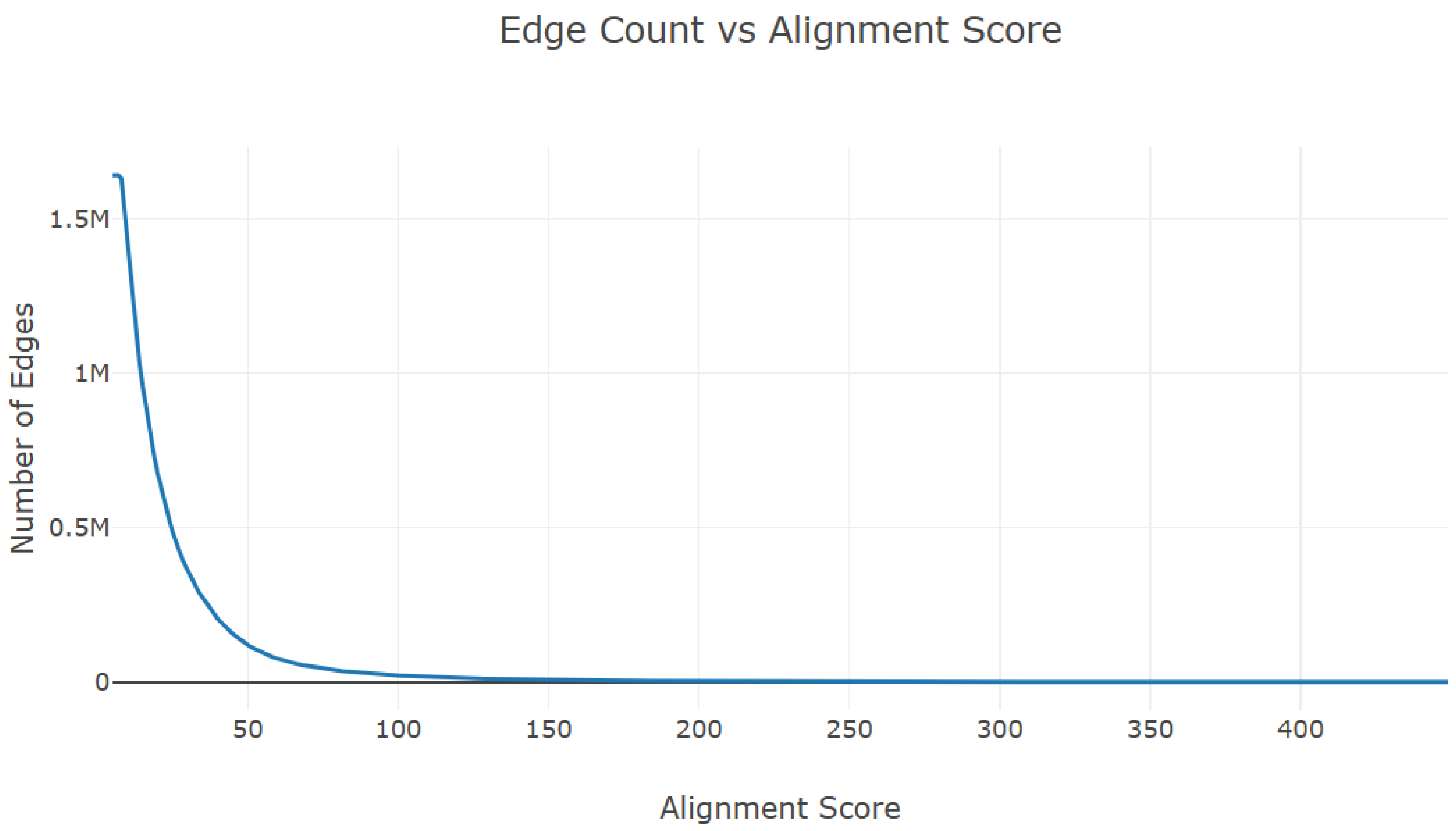
Edge Count vs Alignment Score
1.5M
1M
0.5M
0
50
100
150
200
250
300
350
400
Number of Edges
Alignment Score

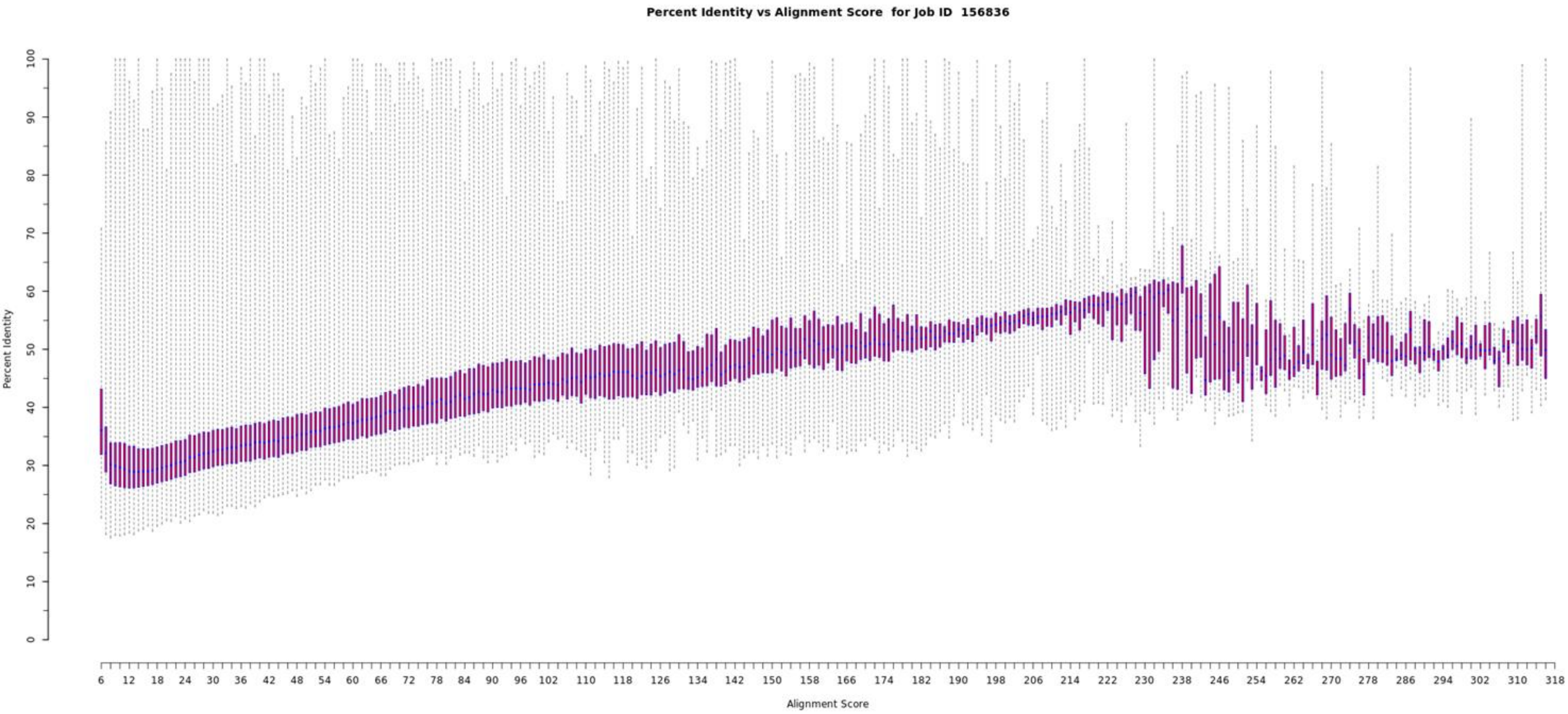

Percent Identity vs Alignment Score for Job ID 156836
Percent Identity
0
10
20
30
40
50
60
70
80
90
100
Alignment Score
6
12
18
24
30
36
42
48
54
60
66
72
78
84
90
96
102
110
118
126
134
142
150
158
166
174
182
190
198
206
214
222
230
238
246
254
262
270
278
286
294
302
310
318

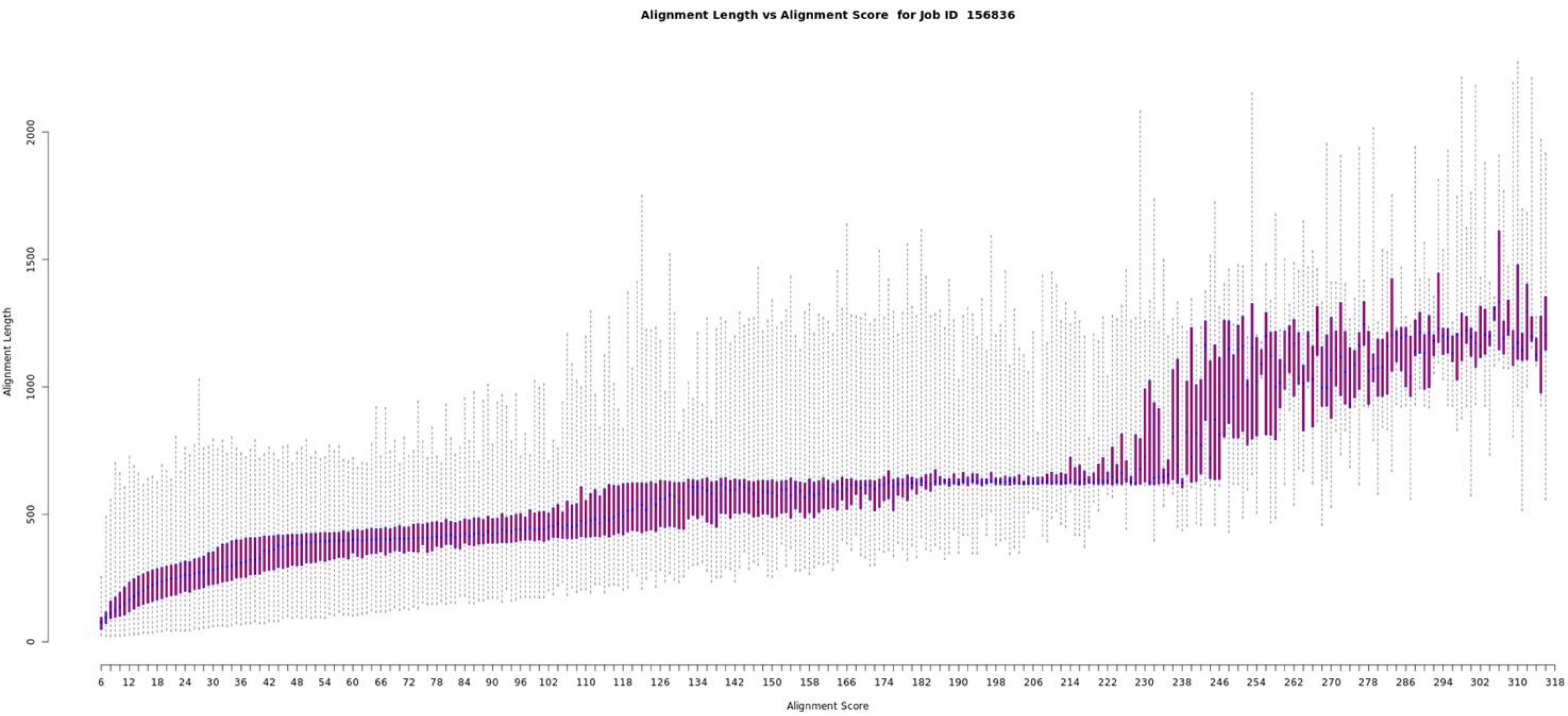
Alignment Length vs Alignment Score for Job ID 156836
Alignment Length
0
500
1000
1500
2000
Alignment Score
6 12 18 24 30 36 42 48 54 60 66 72 78 84 90 96 102 110 118 126 134 142 150 158 166 174 182 190 198 206 214 222 230 238 246 254 262 270 278 286 294 302 310 318

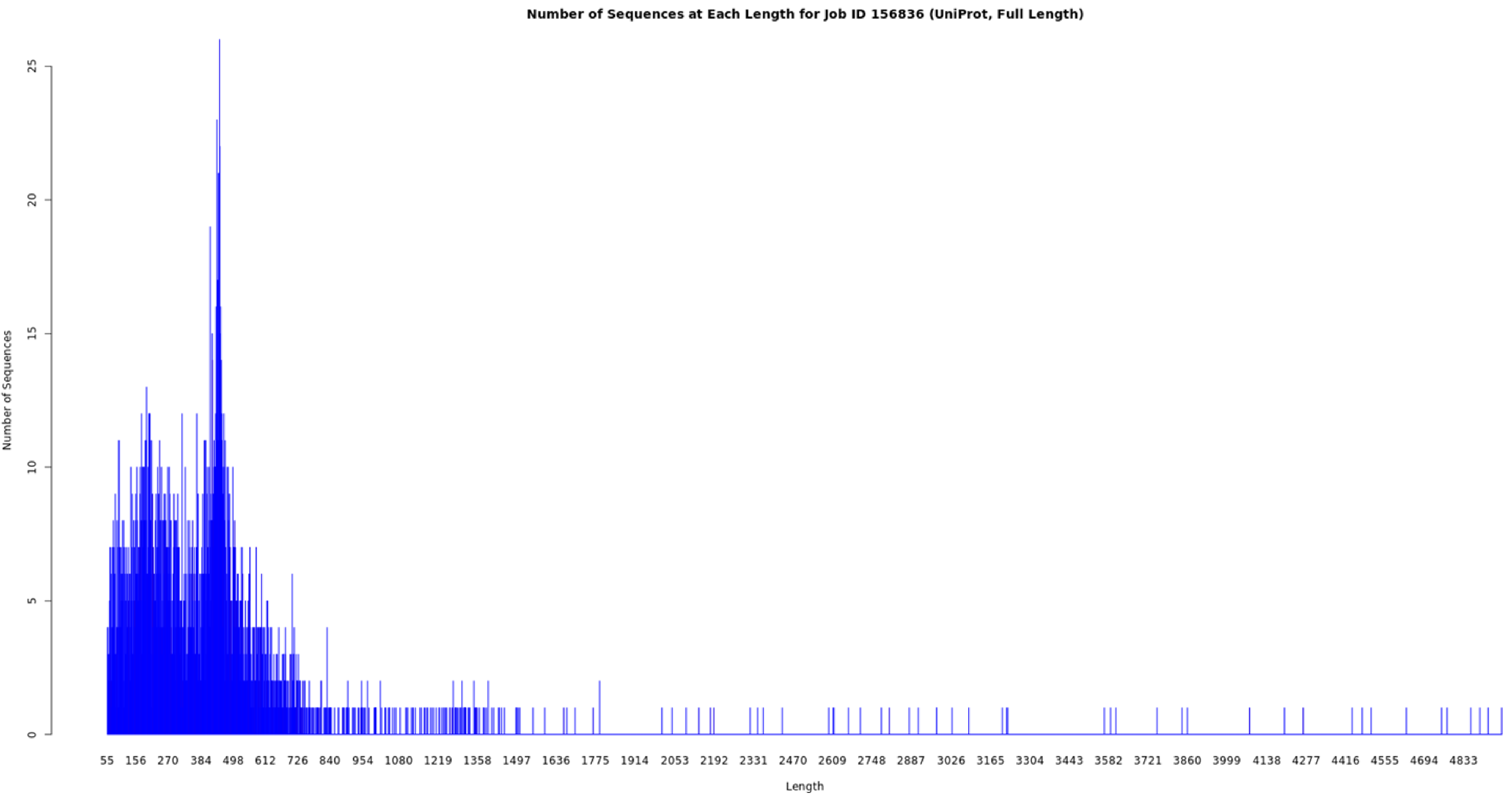
Number of Sequences at Each Length for Job ID 156836 (UniProt, Full Length)
Number of Sequences
0
5
10
15
20
25
55 156 270 384 498 612 726 840 954 1080 1219 1358 1497 1636 1775 1914 2053 2192 2331 2470 2609 2748 2887 3026 3165 3304 3443 3582 3721 3860 3999 4138 4277 4416 4555 4694 4833
Length

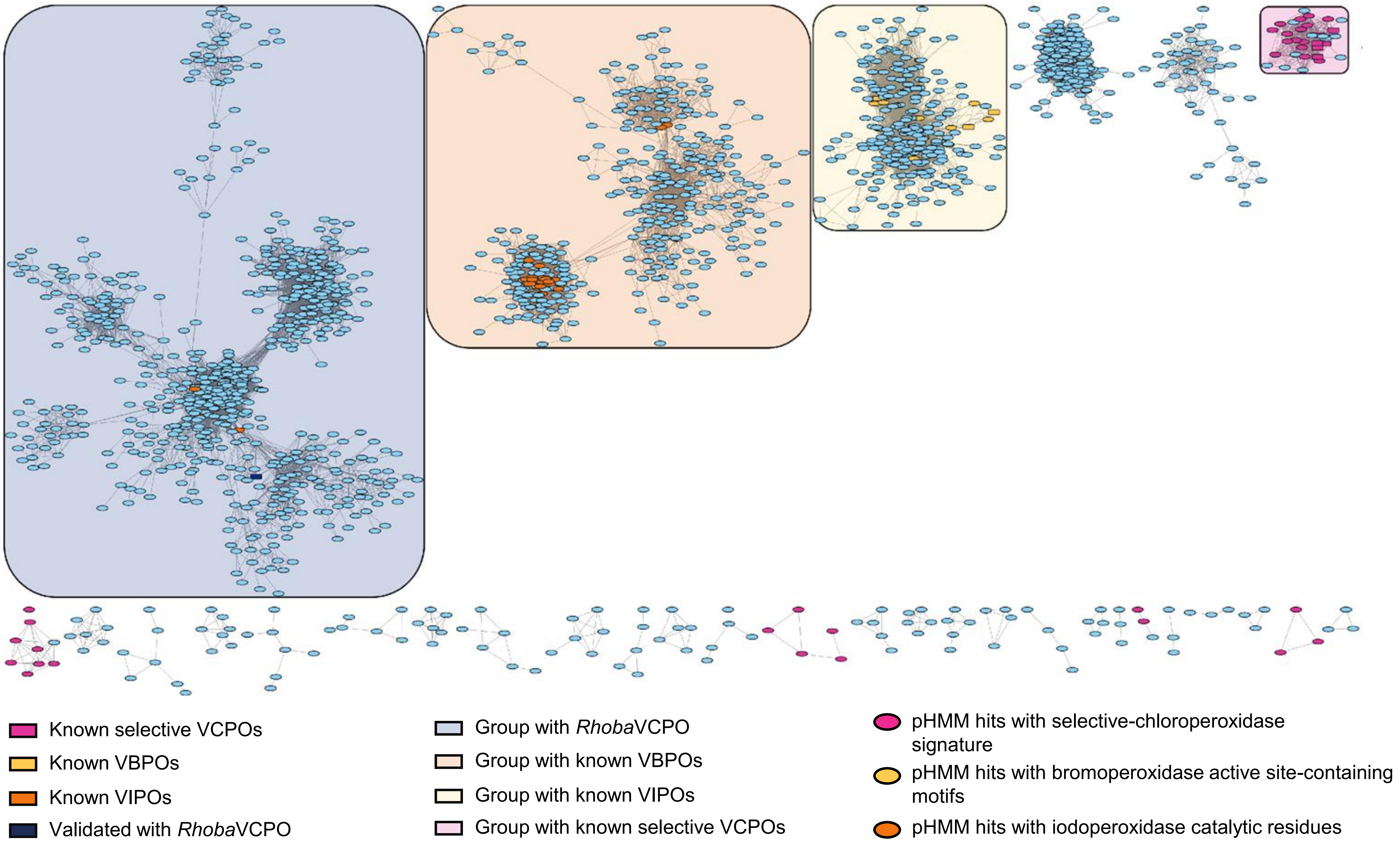

Known selective VCPOs
Known VBPOs
Known VIPOs
Validated with *Rhoba*VCPO
Group with *Rhoba*VCPO
Group with known VBPOs
Group with known VIPOs
Group with known selective VCPOs
pHMM hits with selective-chloroperoxidase signature
pHMM hits with bromoperoxidase active site-containing motifs
pHMM hits with iodoperoxidase catalytic residues

### Dimetal-carboxylate

| | |
|---|---|
| Job Number | 156847 |
| Time Started -- Finished | 7/2 10:00 AM -- 7/2 10:03 AM |
| Database Version | UniProt: 2025-02 / InterPro: 105 |
| Input Option | FASTA (Option C), no FASTA header reading |
| Job Name | 5-dimetal-carboxylate |
| E-Value for SSN Edge Calculation | 5 |
| Uploaded FASTA File | dimetal.fasta |
| Number of Sequences in Uploaded File | 88 |
| Exclude Fragments | No |
| Total Number of Sequences in Dataset | 88 |
| Total Number of Edges | 969 |
| Number of Unique Sequences | 88 |
| Convergence Ratio? | 0.253 |

Alignment Score threshold: 51

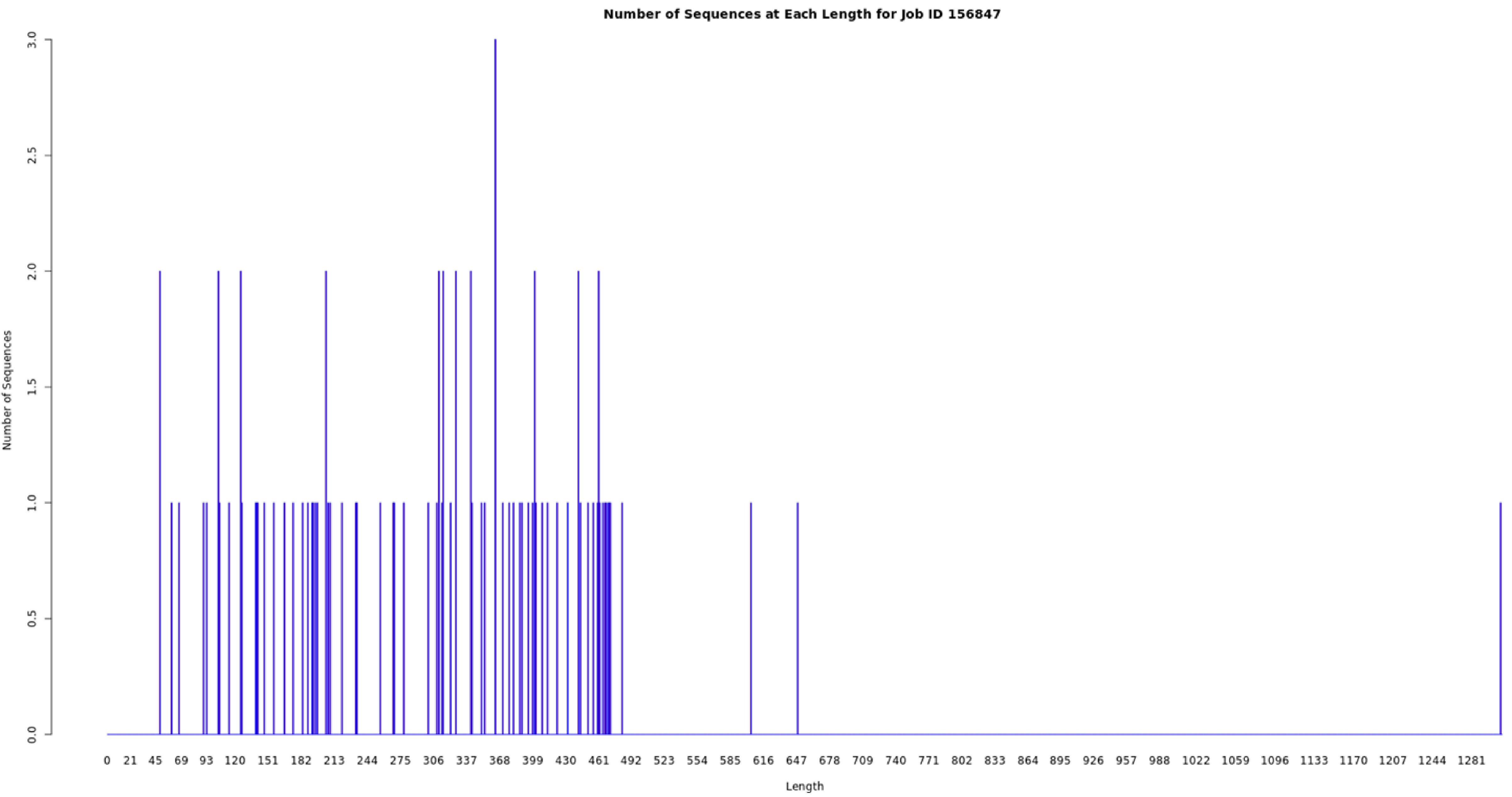

Number of Sequences at Each Length for Job ID 156847
Number of Sequences
0.0
0.5
1.0
1.5
2.0
2.5
3.0
0 21 45 69 93 120 151 182 213 244 275 306 337 368 399 430 461 492 523 554 585 616 647 678 709 740 771 802 833 864 895 926 957 988 1022 1059 1096 1133 1170 1207 1244 1281
Length

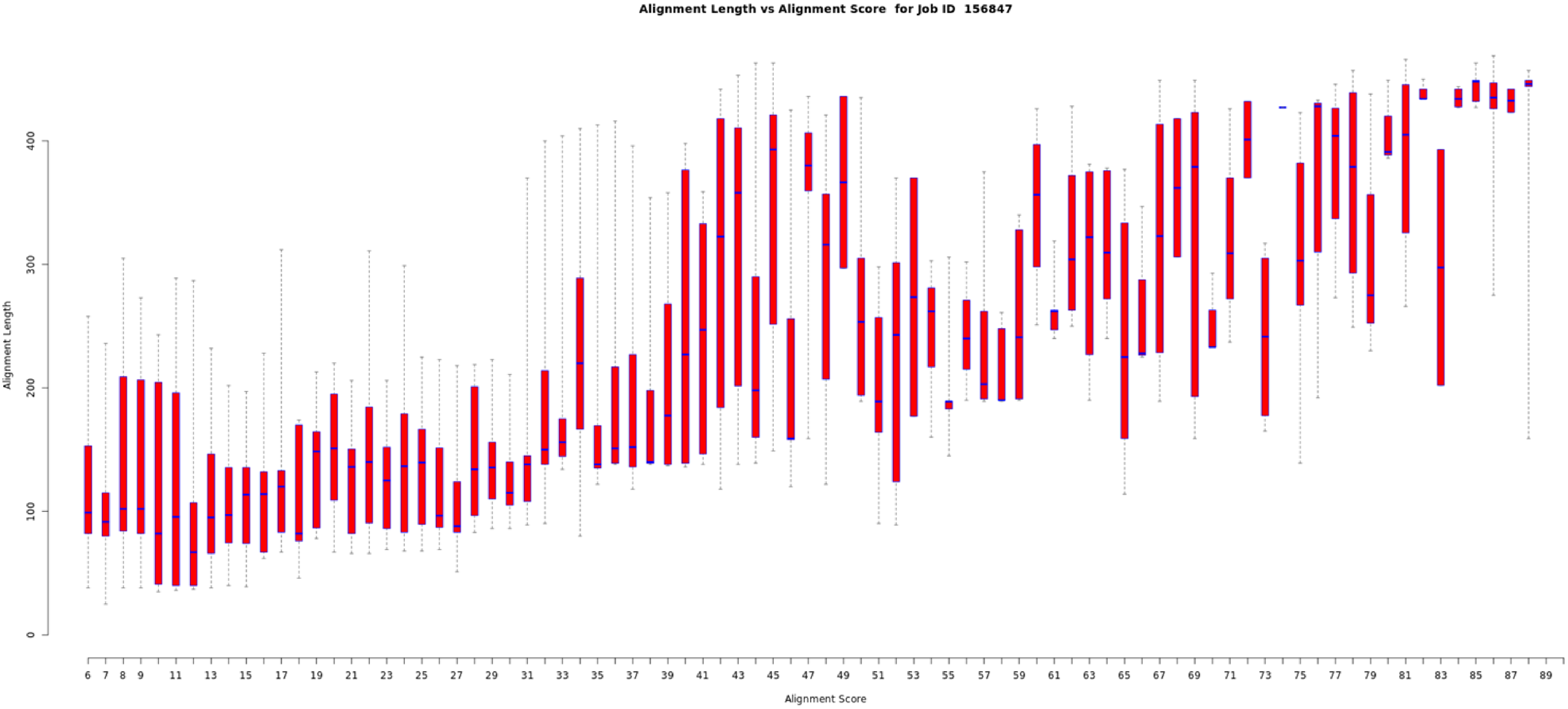
Alignment Length vs Alignment Score for Job ID 156847
Alignment Length
0
100
200
300
400
Alignment Score
6 7 8 9 11 13 15 17 19 21 23 25 27 29 31 33 35 37 39 41 43 45 47 49 51 53 55 57 59 61 63 65 67 69 71 73 75 77 79 81 83 85 87 89

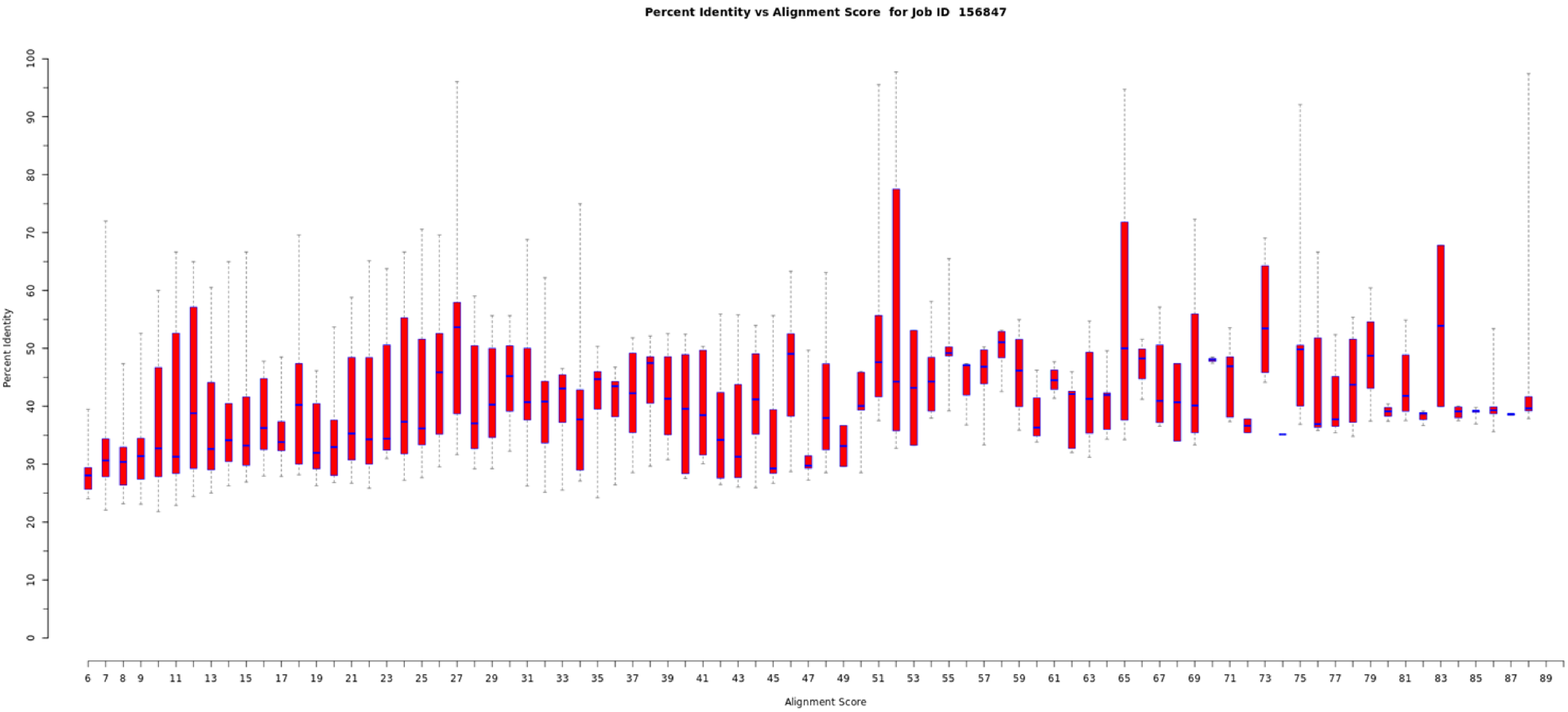
Percent Identity vs Alignment Score for Job ID 156847
Percent Identity
0
10
20
30
40
50
60
70
80
90
100
Alignment Score
6 7 8 9 11 13 15 17 19 21 23 25 27 29 31 33 35 37 39 41 43 45 47 49 51 53 55 57 59 61 63 65 67 69 71 73 75 77 79 81 83 85 87 89

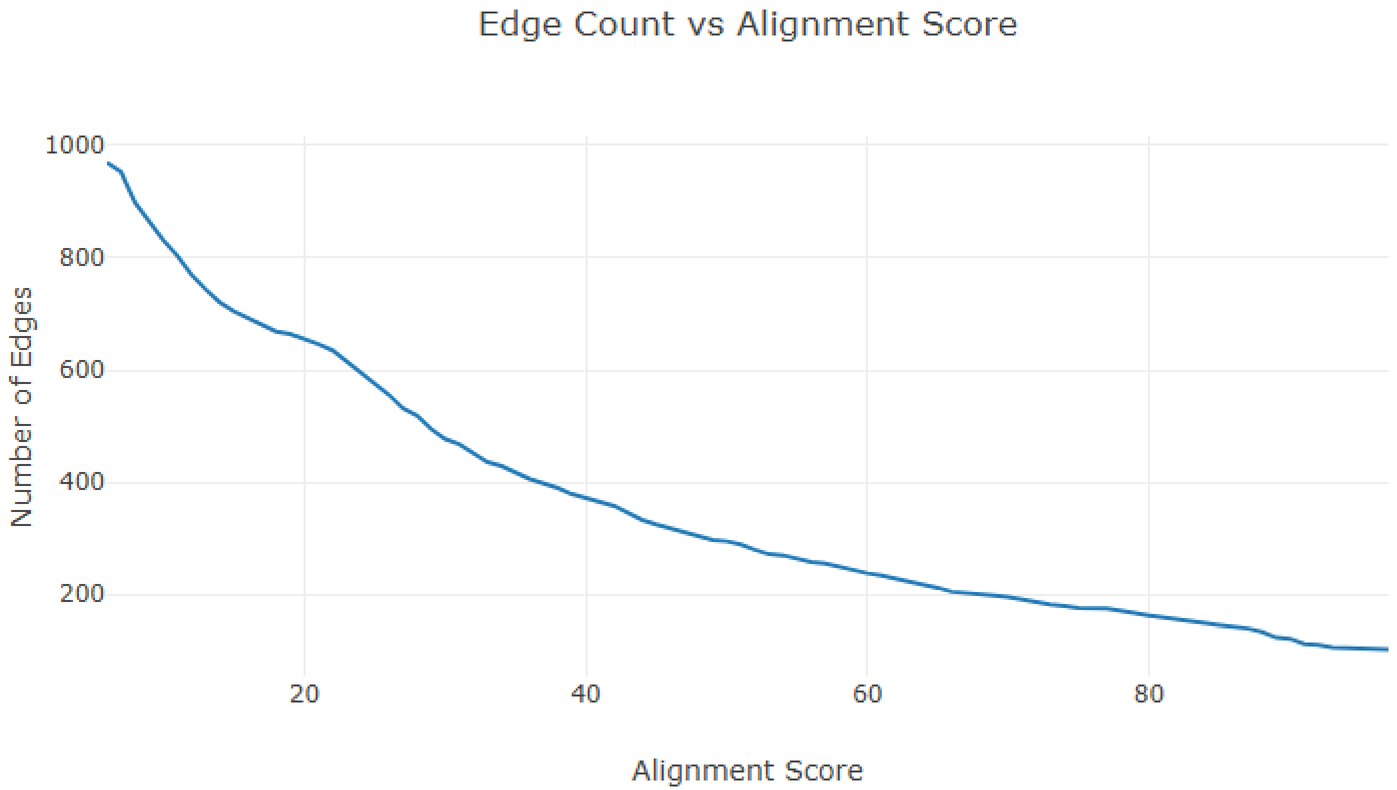
Edge Count vs Alignment Score
Number of Edges
1000
800
600
400
200
20
40
60
80
Alignment Score

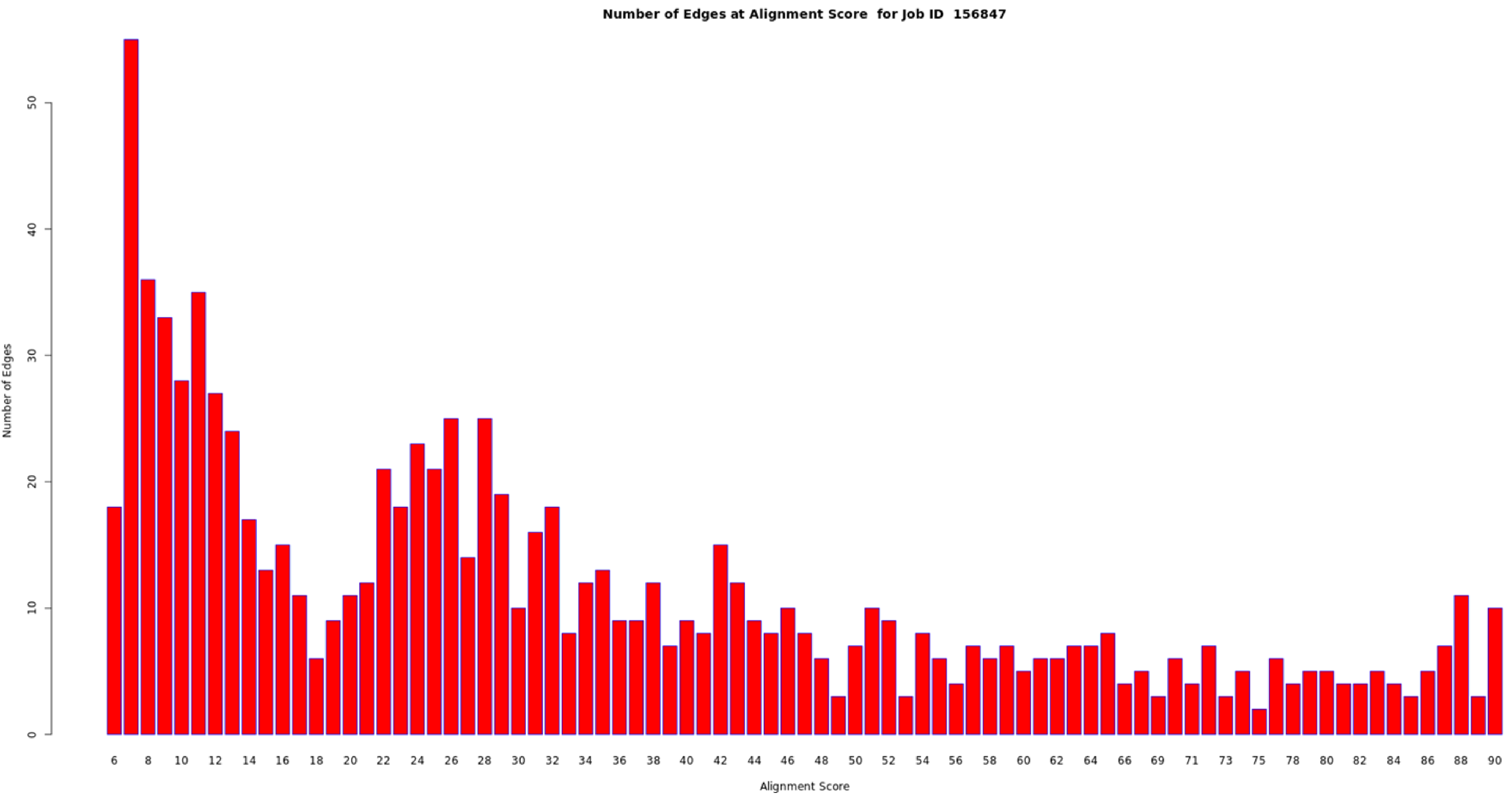
Number of Edges at Alignment Score for Job ID 156847
Number of Edges
0
10
20
30
40
50
6
8
10
12
14
16
18
20
22
24
26
28
30
32
34
36
38
40
42
44
46
48
50
52
54
56
58
60
62
64
66
69
71
73
75
78
80
82
84
86
88
90
Alignment Score

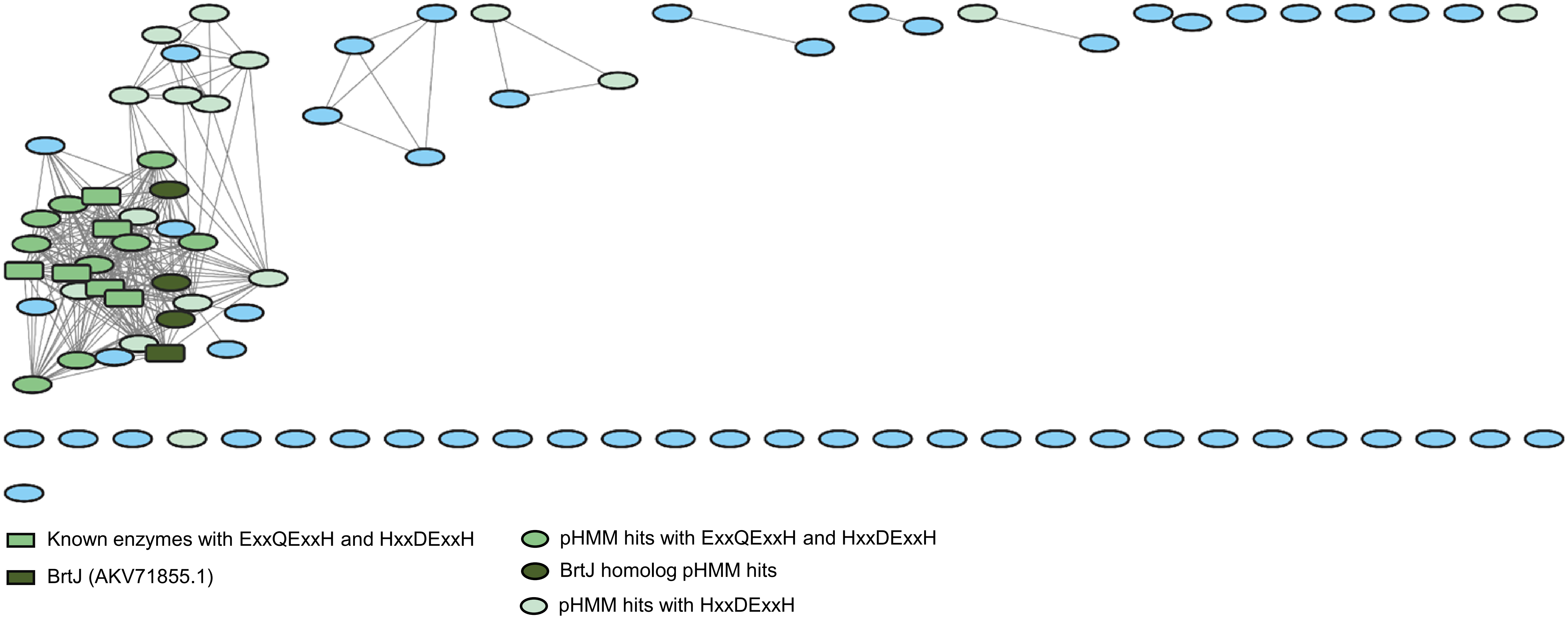
Known enzymes with ExxQExxH and HxxDExxH
BrtJ (AKV71855.1)
pHMM hits with ExxQExxH and HxxDExxH
BrtJ homolog pHMM hits
pHMM hits with HxxDExxH

### SAM-dependent

| | |
|---|---|
| Job Number | 156962 |
| Time Started -- Finished | 7/3 01:45 PM -- 7/3 01:49 PM |
| Database Version | UniProt: 2025-02 / InterPro: 105 |
| Input Option | FASTA (Option C), no FASTA header reading |
| Job Name | sam-halogenases |
| E-Value for SSN Edge Calculation | 5 |
| Uploaded FASTA File | sam_ssn.fasta |
| Number of Sequences in Uploaded File | 2,605 |
| Exclude Fragments | No |
| Total Number of Sequences in Dataset | 2,604 |
| Total Number of Edges | 2,343,179 |
| Number of Unique Sequences | 2,604 |
| Convergence Ratio? | 0.691 |

Alignment Score threshold: 60

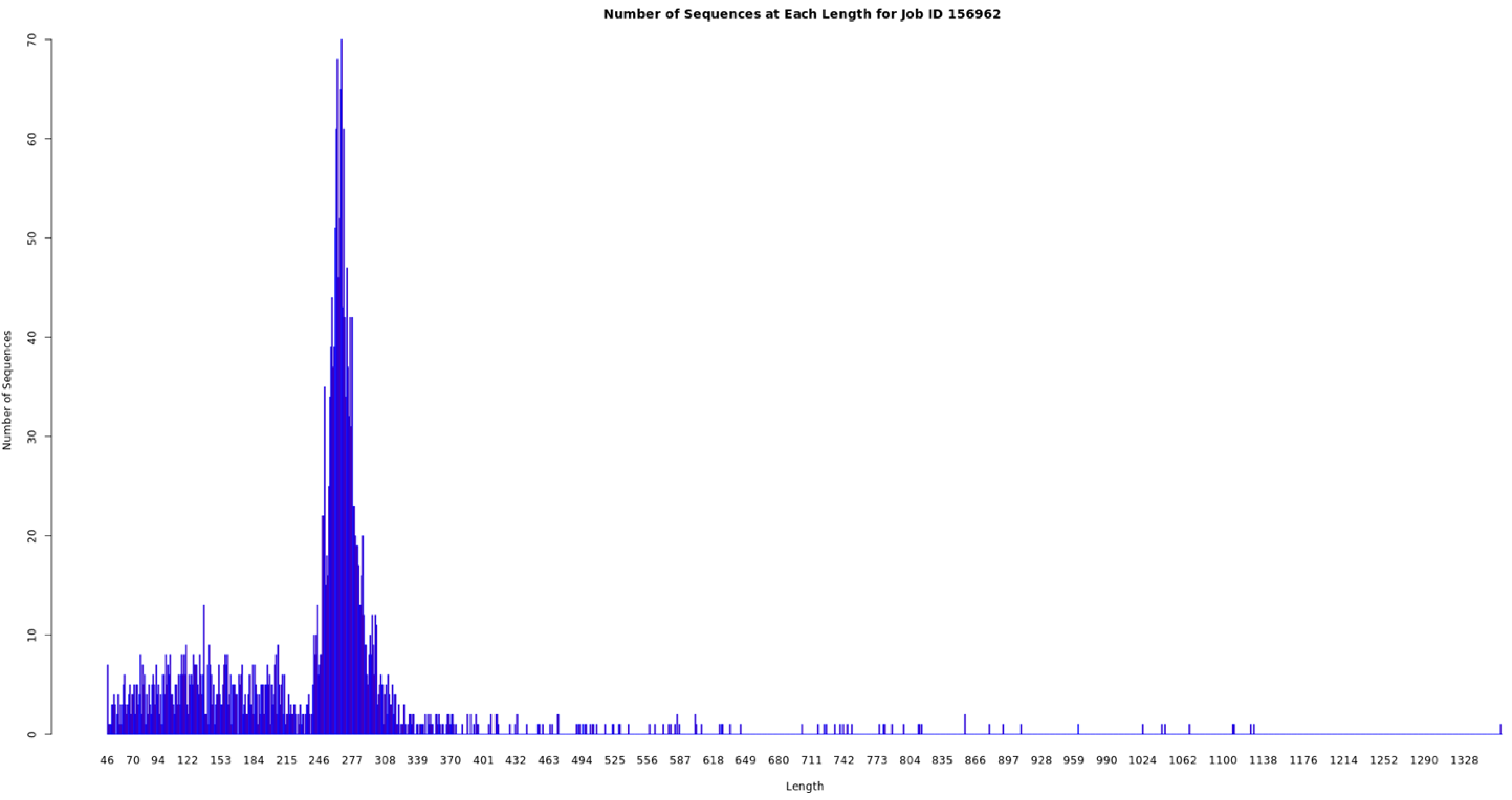
Number of Sequences at Each Length for Job ID 156962
Number of Sequences
0
10
20
30
40
50
60
70
46 70 94 122 153 184 215 246 277 308 339 370 401 432 463 494 525 556 587 618 649 680 711 742 773 804 835 866 897 928 959 990 1024 1062 1100 1138 1176 1214 1252 1290 1328
Length

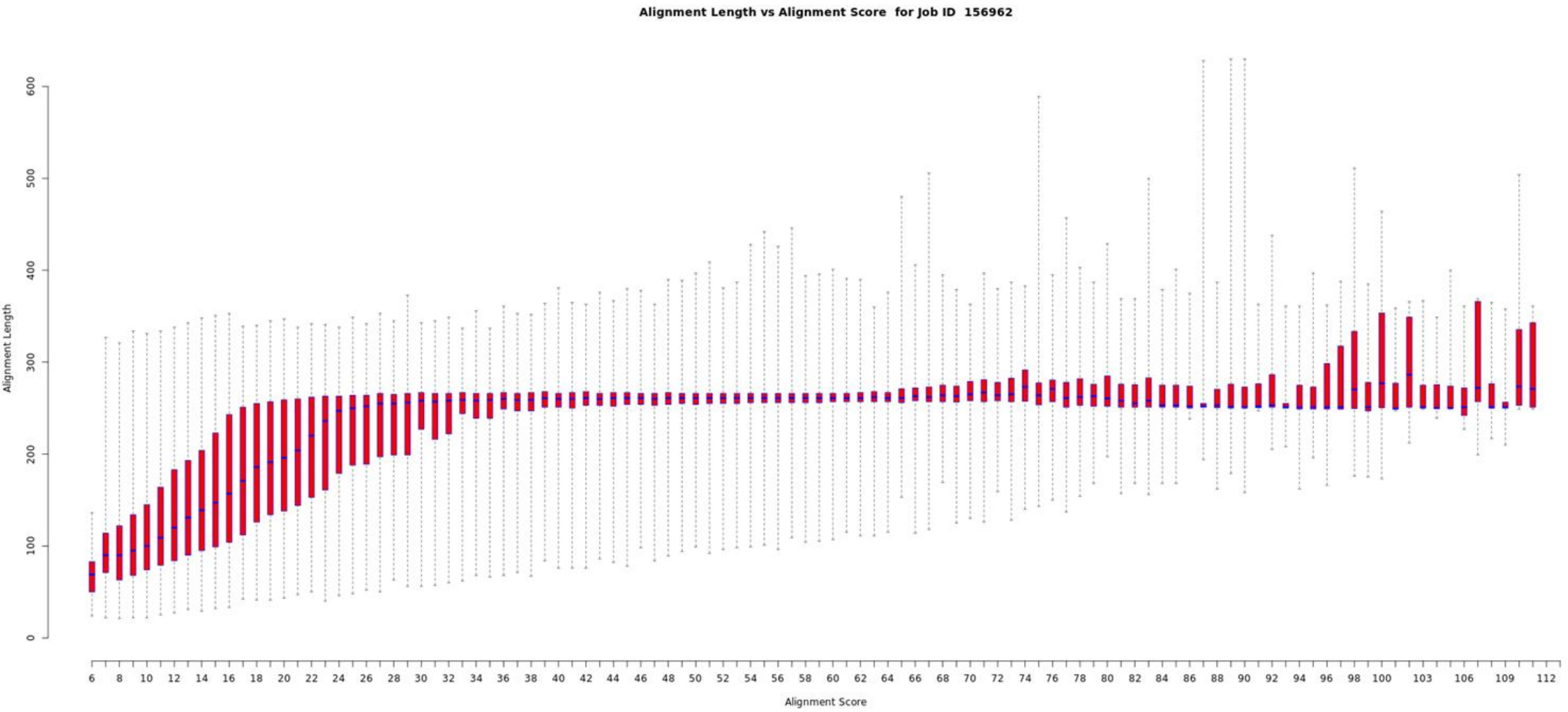
Alignment Length vs Alignment Score for Job ID 156962
Alignment Length
Alignment Score

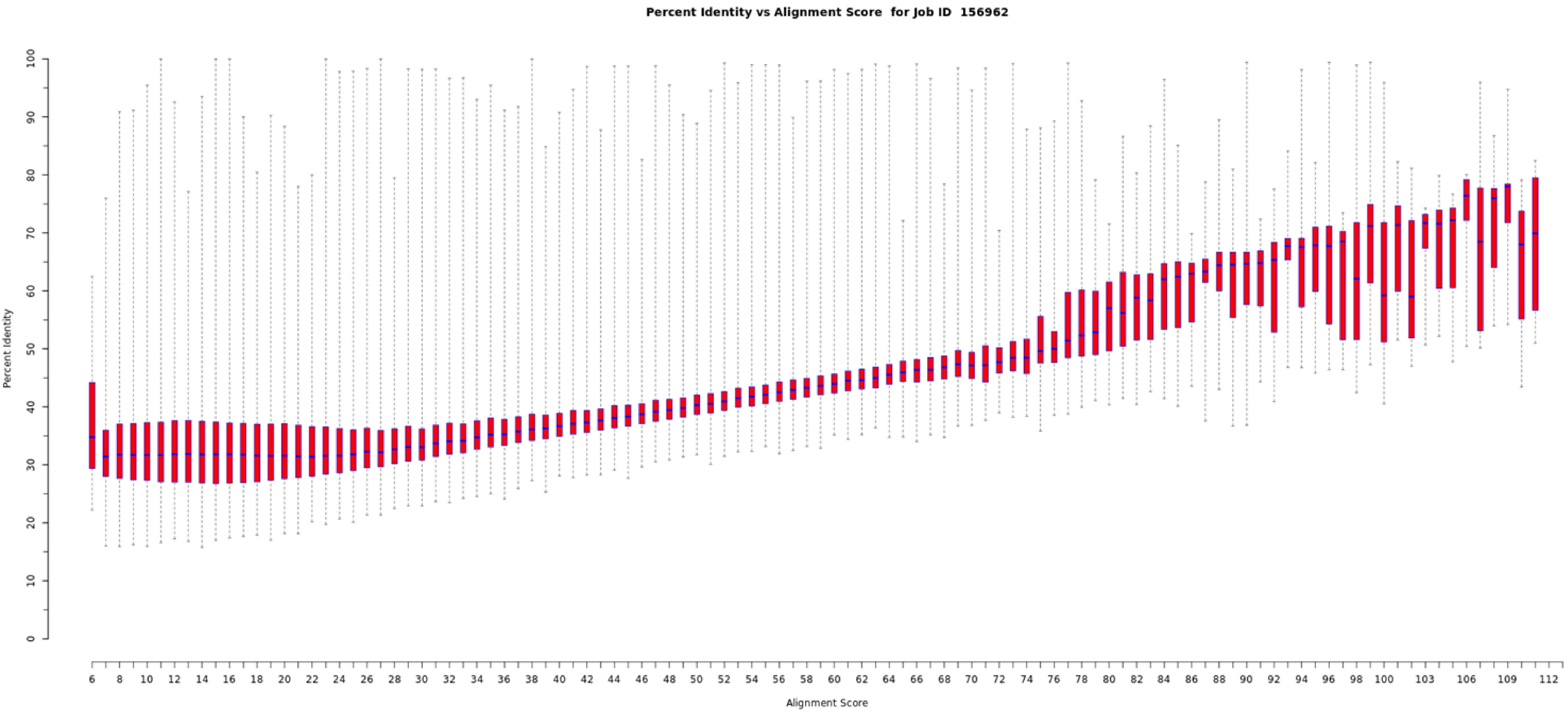

Percent Identity vs Alignment Score for Job ID 156962
Percent Identity
0
10
20
30
40
50
60
70
80
90
100
Alignment Score
6
8
10
12
14
16
18
20
22
24
26
28
30
32
34
36
38
40
42
44
46
48
50
52
54
56
58
60
62
64
66
68
70
72
74
76
78
80
82
84
86
88
90
92
94
96
98
100
103
106
109
112

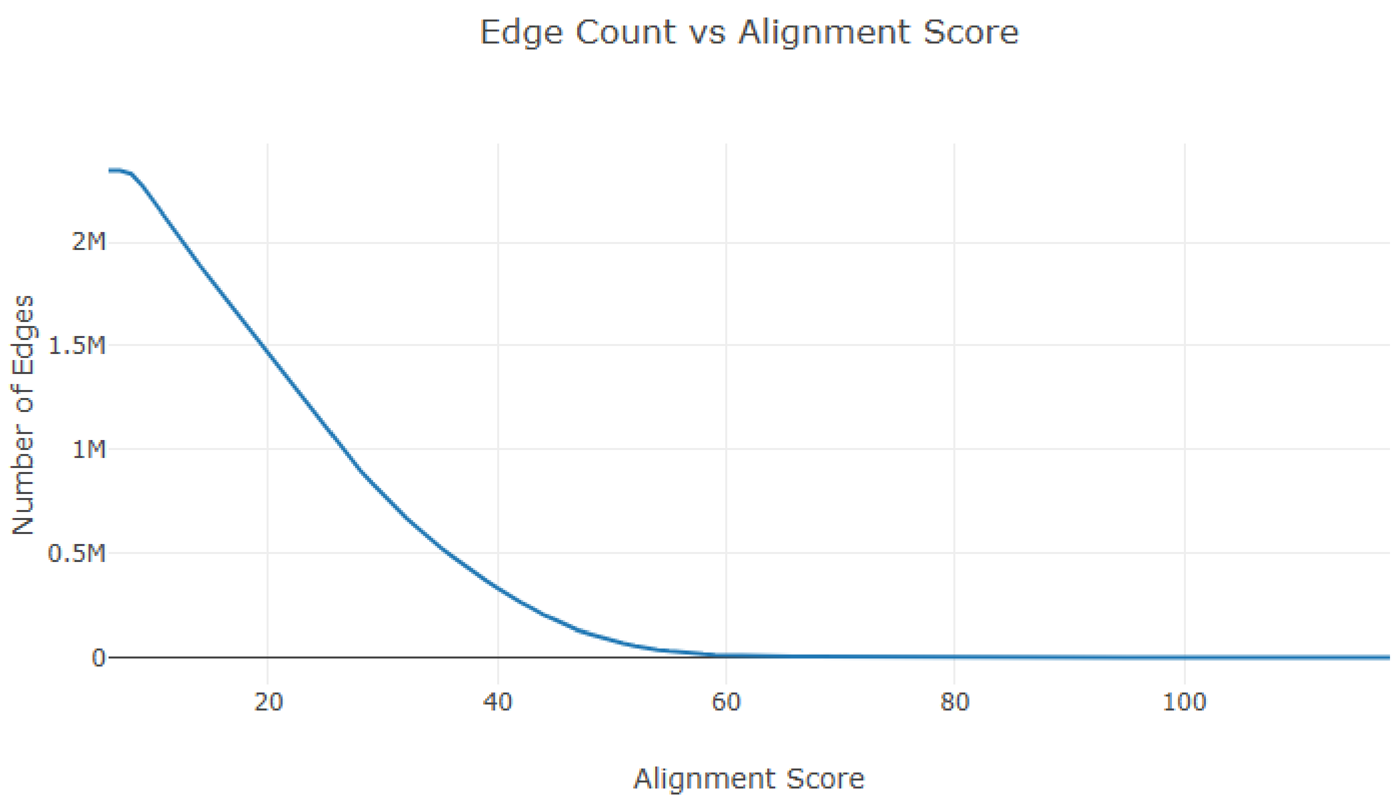
Edge Count vs Alignment Score
Number of Edges
2M
1.5M
1M
0.5M
0
20
40
60
80
100
Alignment Score

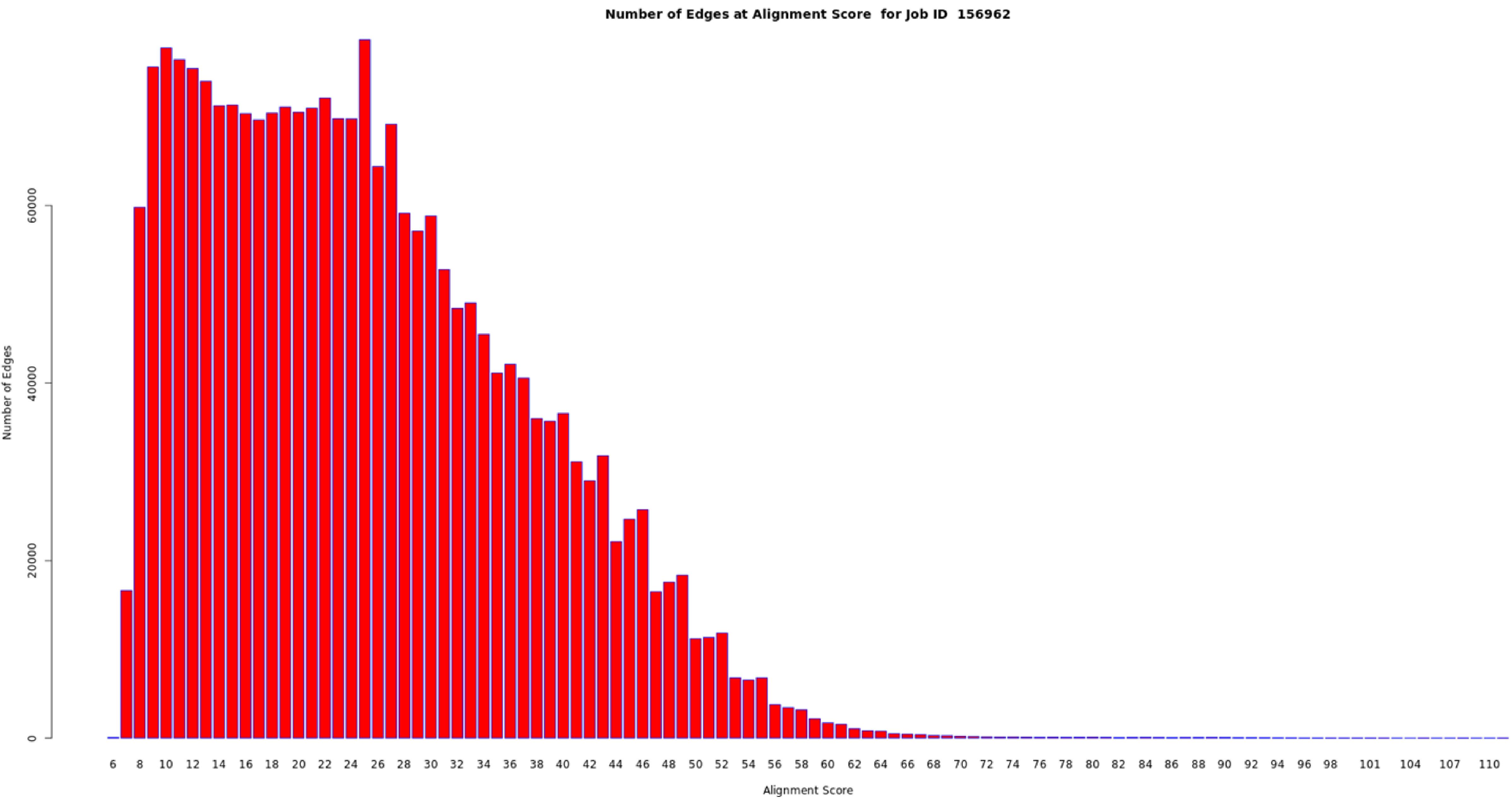

Number of Edges at Alignment Score for Job ID 156962
Number of Edges
0
20000
40000
60000
6 8 10 12 14 16 18 20 22 24 26 28 30 32 34 36 38 40 42 44 46 48 50 52 54 56 58 60 62 64 66 68 70 72 74 76 78 80 82 84 86 88 90 92 94 96 98 101 104 107 110
Alignment Score

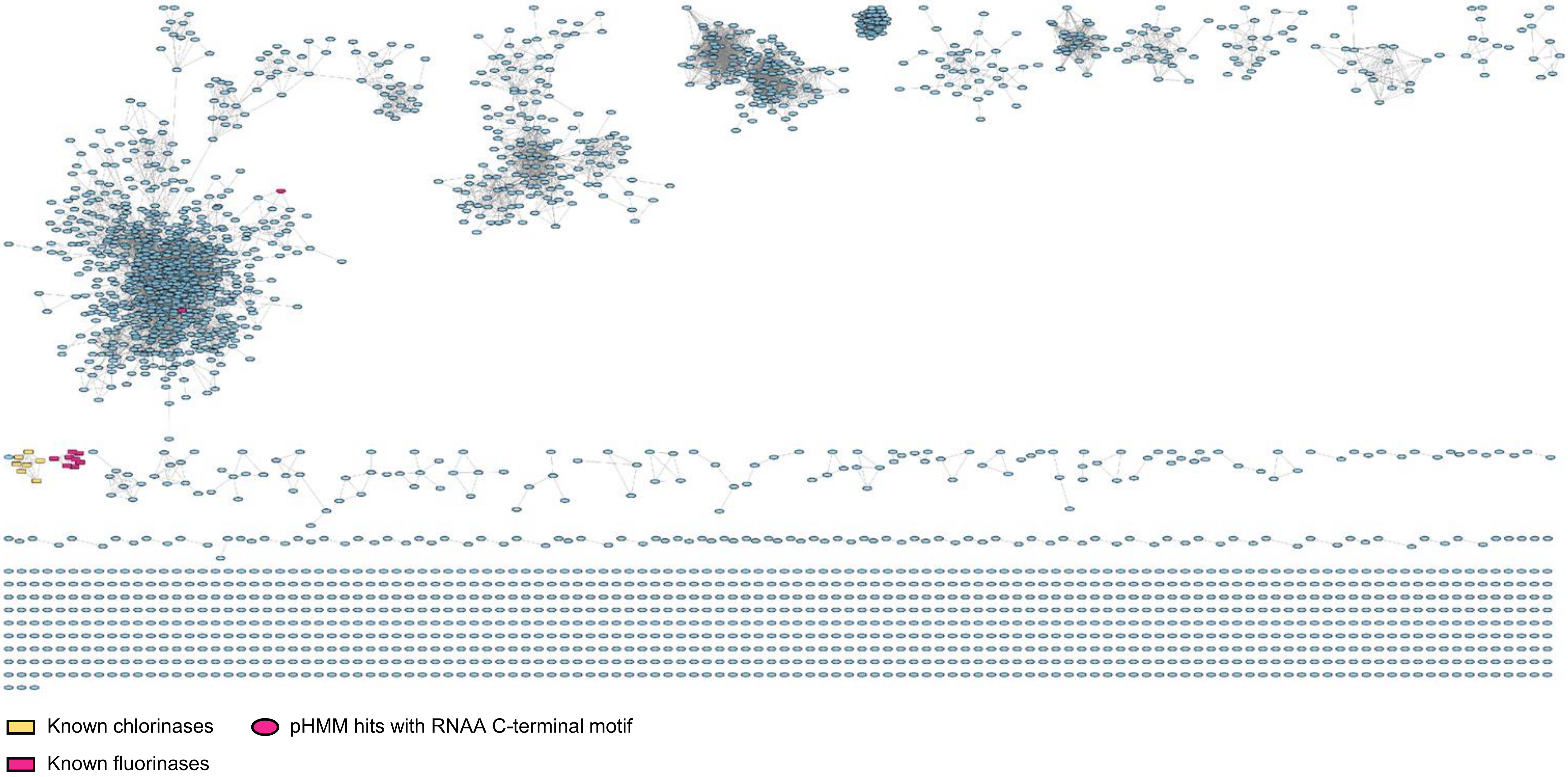
Known chlorinases
pHMM hits with RNAA C-terminal motif
Known fluorinases

### Flavin-dependent (unconventional)

| | |
|---|---|
| Job Number | 157052 |
| Time Started -- Finished | 7/6 01:20 AM -- 7/6 01:23 AM |
| Database Version | UniProt: 2025-02 / InterPro: 105 |
| Input Option | FASTA (Option C), no FASTA header reading |
| Job Name | FDH-unconventional |
| E-Value for SSN Edge Calculation | 5 |
| Uploaded FASTA File | fdh_unconventional.fasta |
| Number of Sequences in Uploaded File | 193 |
| Exclude Fragments | No |
| Total Number of Sequences in Dataset | 193 |
| Total Number of Edges | 3,530 |
| Number of Unique Sequences | 193 |
| Convergence Ratio? | 0.191 |

Alignment Score threshold: 45

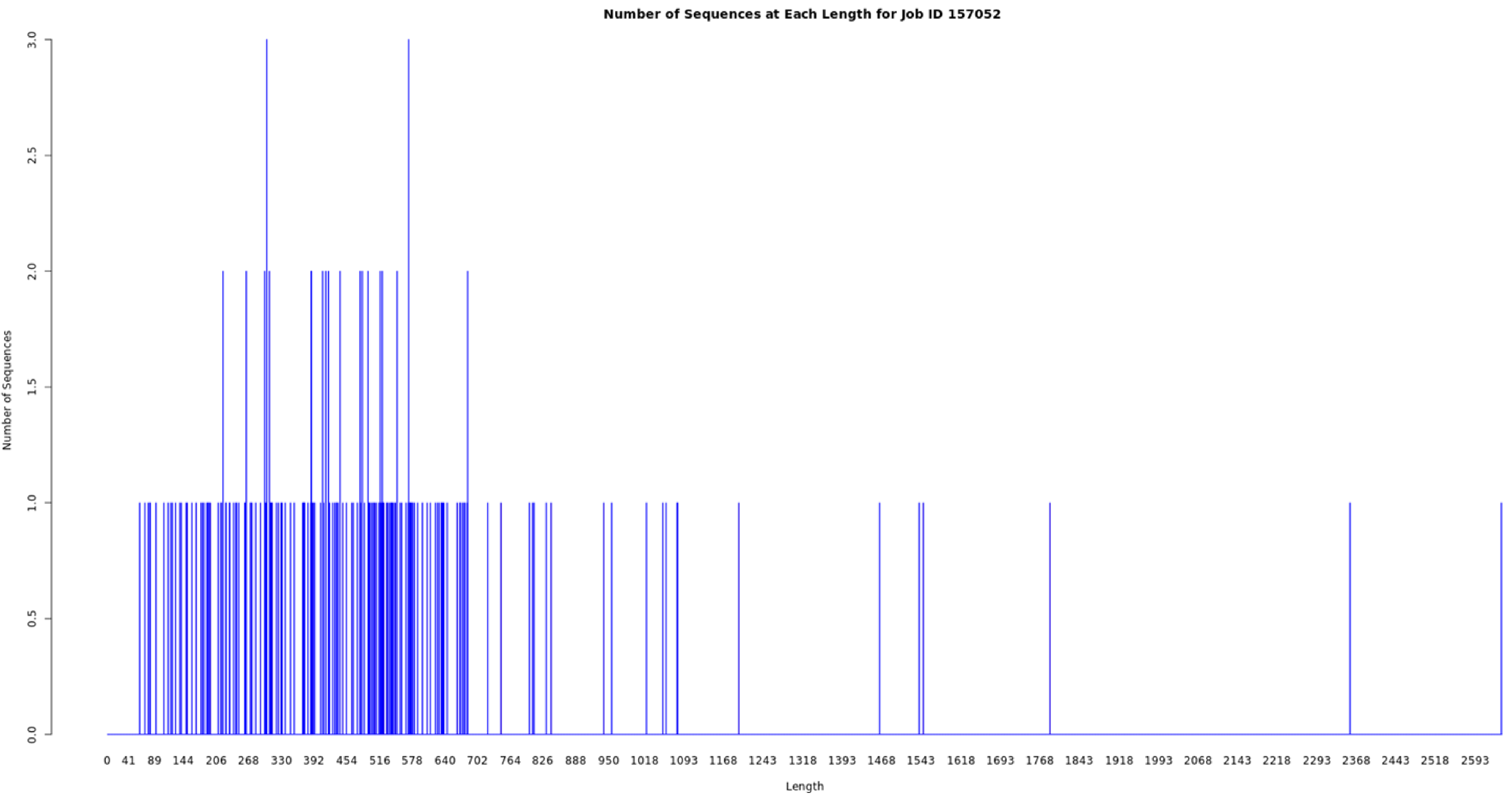
Number of Sequences at Each Length for Job ID 157052
Number of Sequences
0.0
0.5
1.0
1.5
2.0
2.5
3.0
Length
0 41 89 144 206 268 330 392 454 516 578 640 702 764 826 888 950 1018 1093 1168 1243 1318 1393 1468 1543 1618 1693 1768 1843 1918 1993 2068 2143 2218 2293 2368 2443 2518 2593

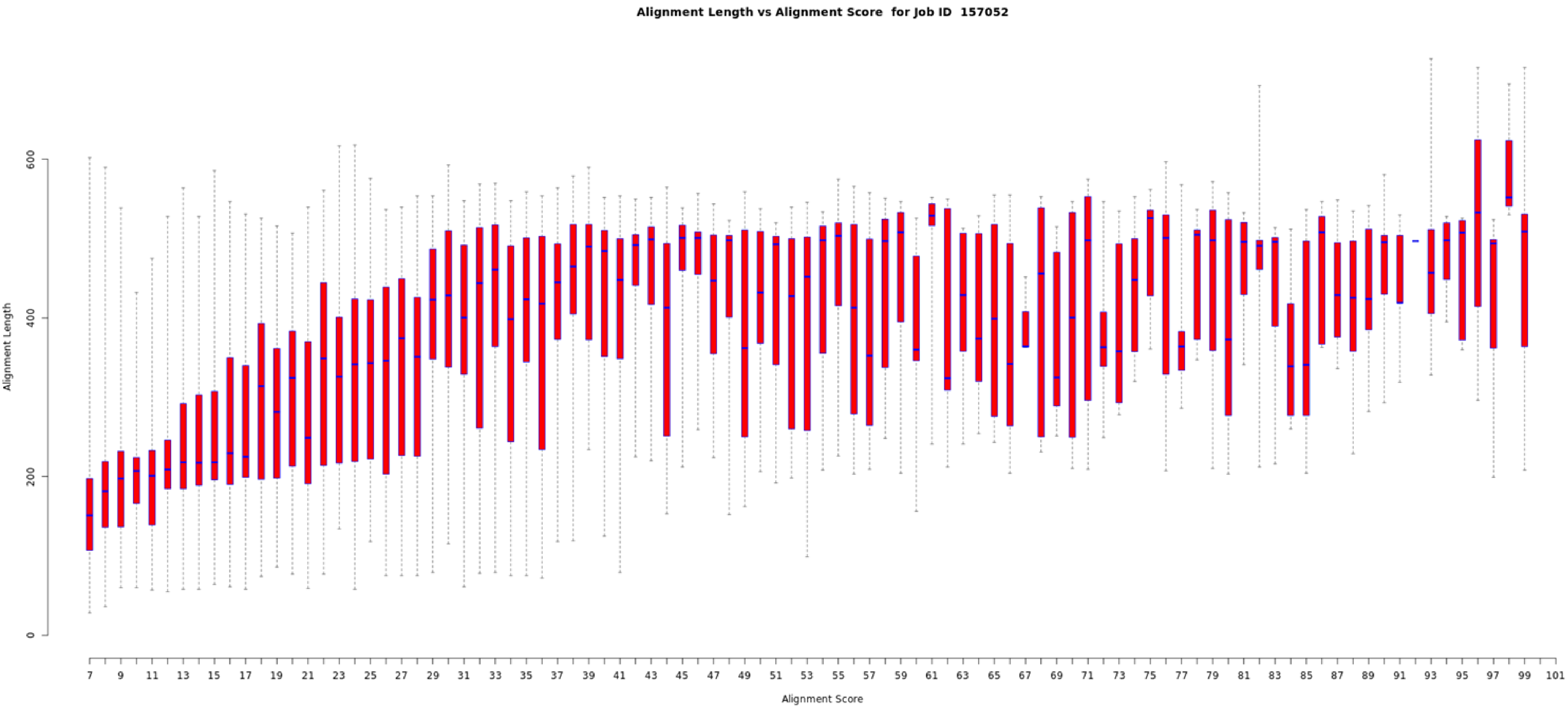
Alignment Length vs Alignment Score for Job ID 157052
Alignment Length
0
200
400
600
Alignment Score
7 9 11 13 15 17 19 21 23 25 27 29 31 33 35 37 39 41 43 45 47 49 51 53 55 57 59 61 63 65 67 69 71 73 75 77 79 81 83 85 87 89 91 93 95 97 99 101

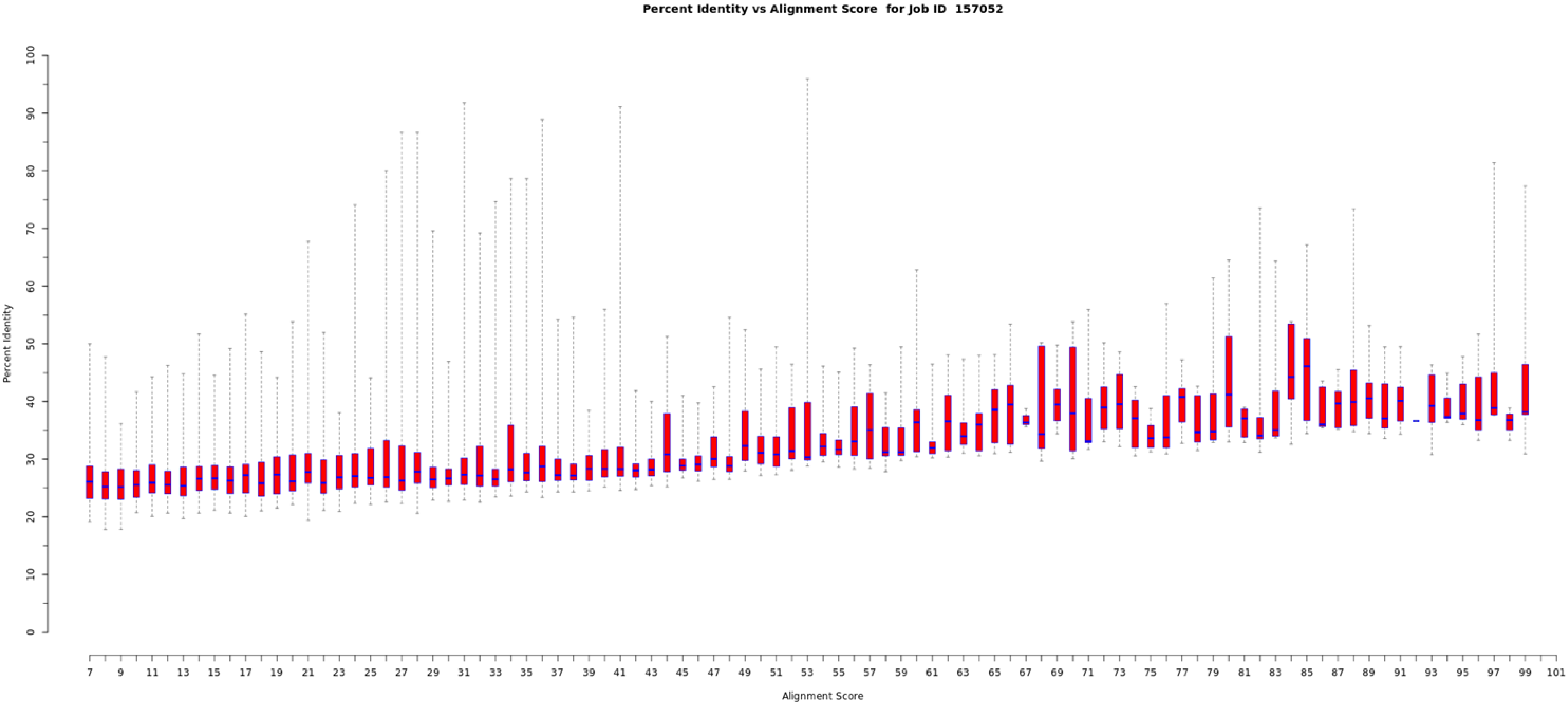
Percent Identity vs Alignment Score for Job ID 157052
Percent Identity
0
10
20
30
40
50
60
70
80
90
100
Alignment Score
7
9
11
13
15
17
19
21
23
25
27
29
31
33
35
37
39
41
43
45
47
49
51
53
55
57
59
61
63
65
67
69
71
73
75
77
79
81
83
85
87
89
91
93
95
97
99
101

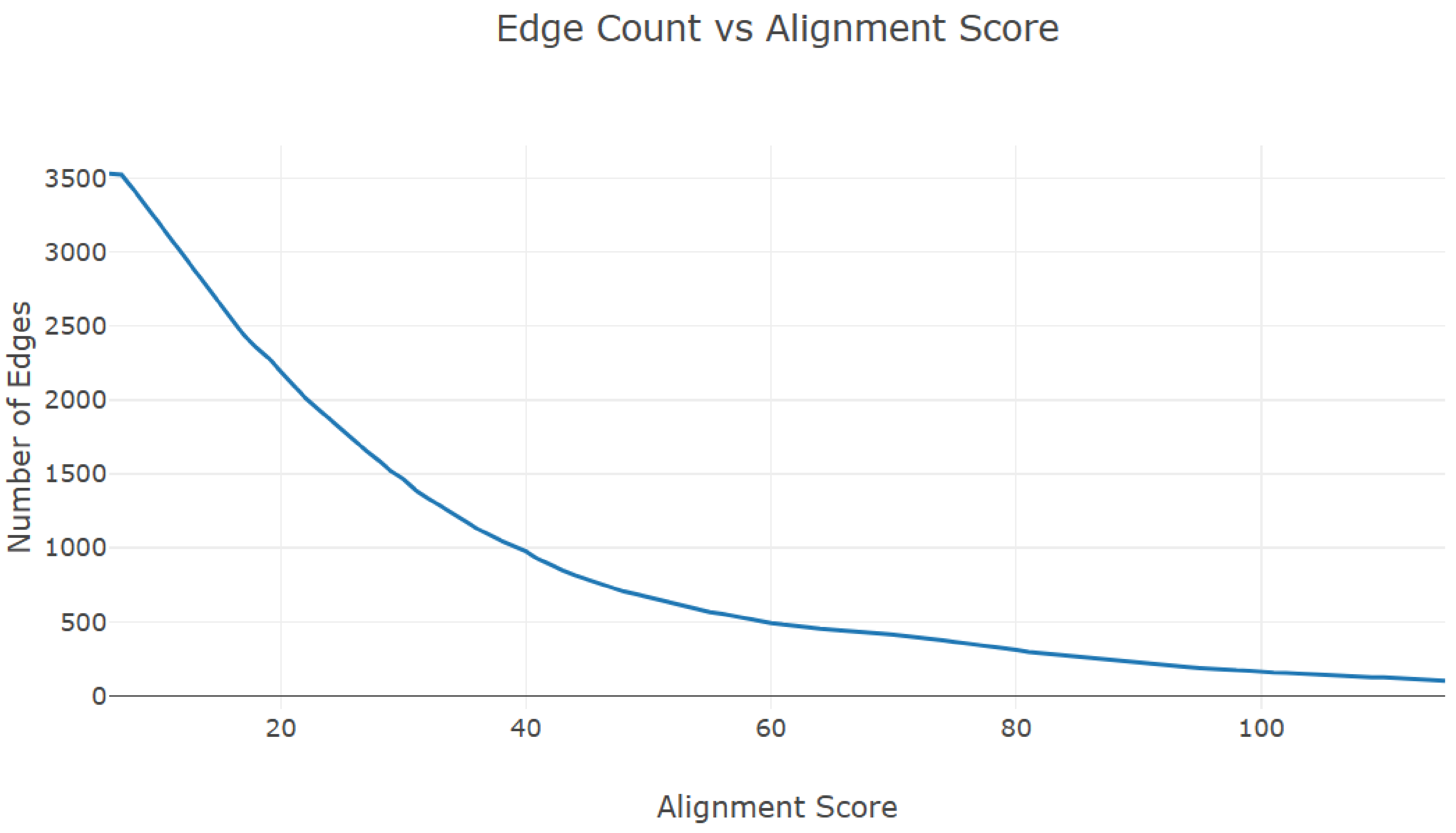
Edge Count vs Alignment Score
Number of Edges
3500
3000
2500
2000
1500
1000
500
0
20
40
60
80
100
Alignment Score

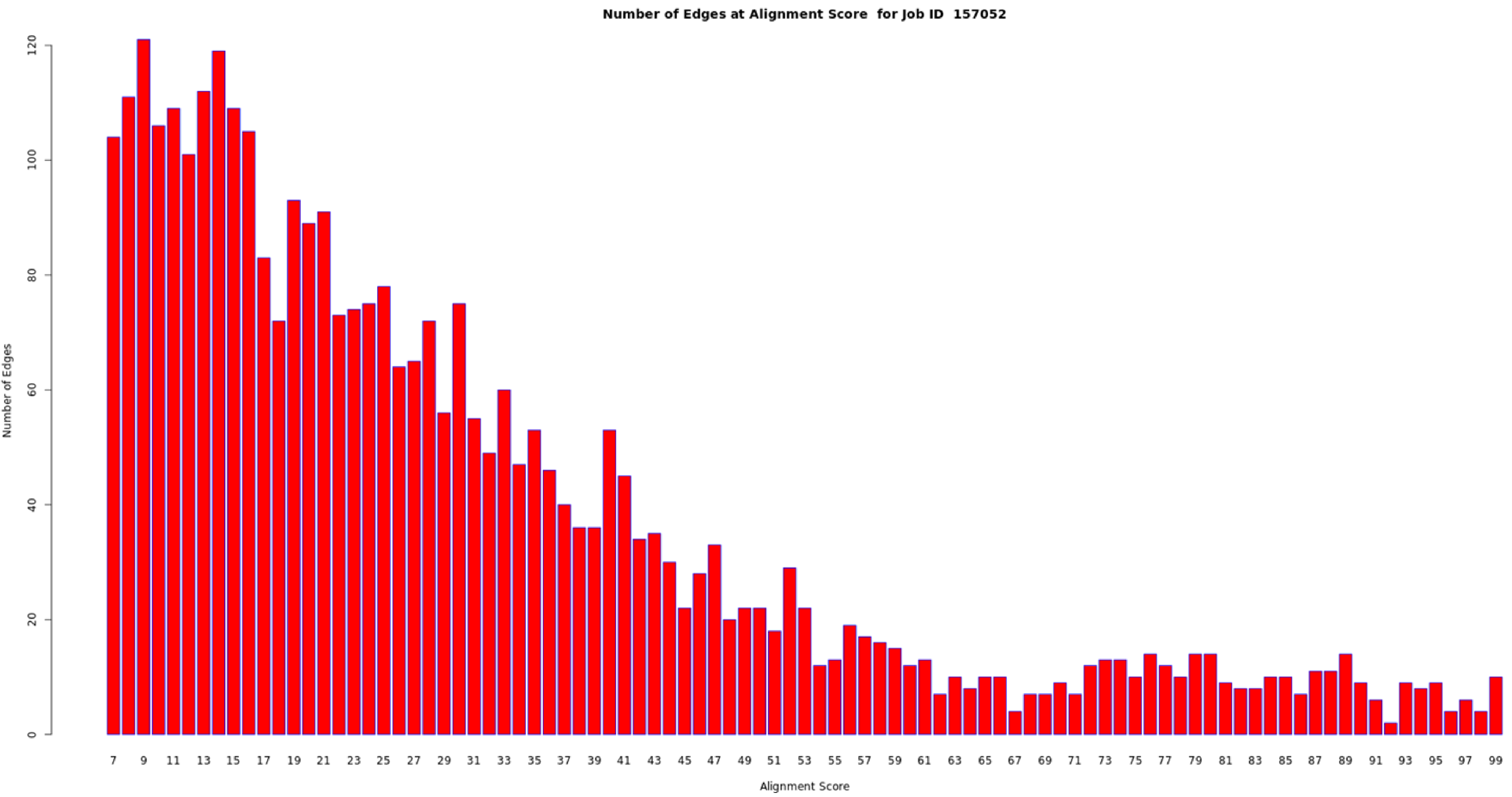
Number of Edges at Alignment Score for Job ID 157052
Number of Edges
Alignment Score
0
20
40
60
80
100
120
7
9
11
13
15
17
19
21
23
25
27
29
31
33
35
37
39
41
43
45
47
49
51
53
55
57
59
61
63
65
67
69
71
73
75
77
79
81
83
85
87
89
91
93
95
97
99

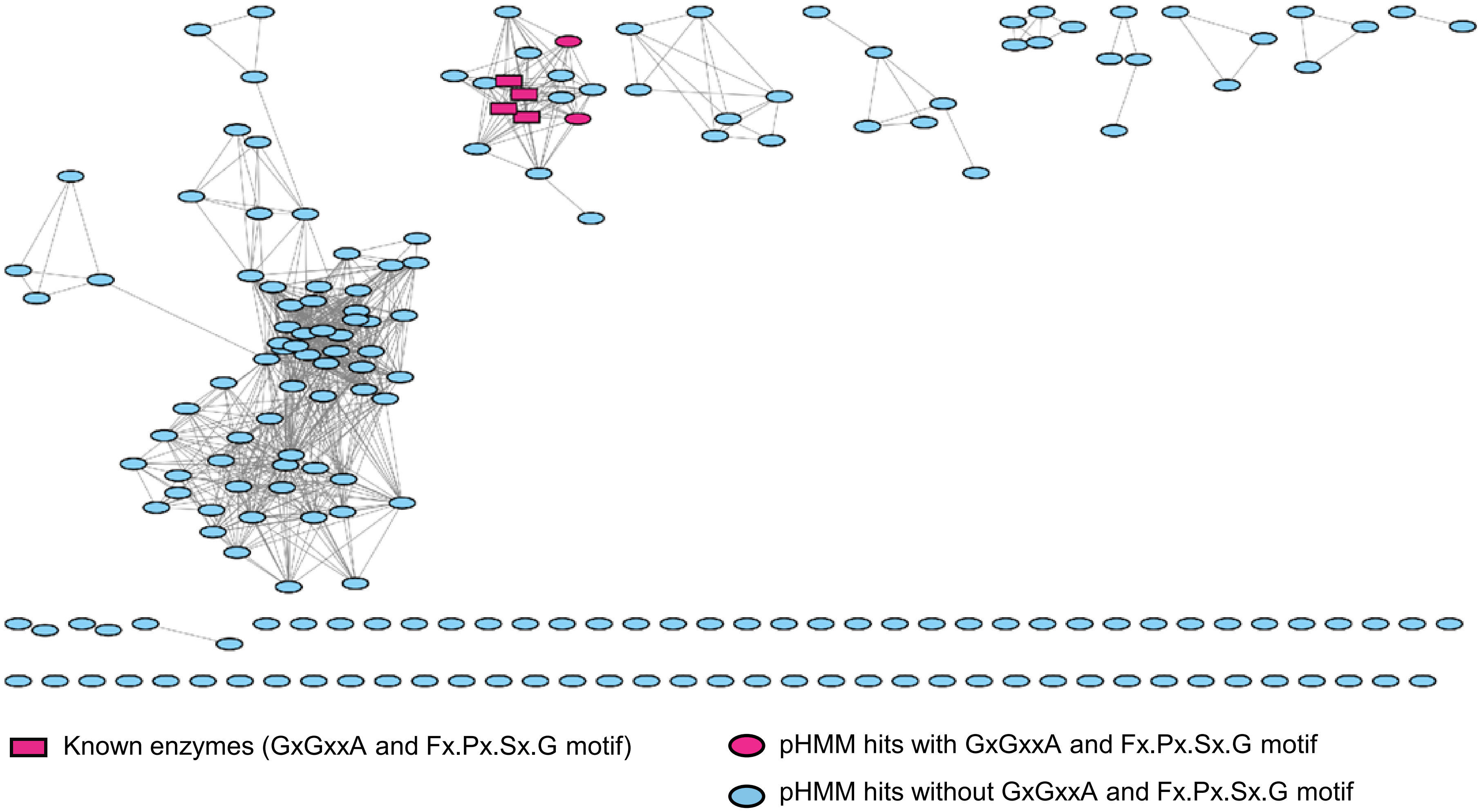
Known enzymes (GxGxxA and Fx.Px.Sx.G motif)
pHMM hits with GxGxxA and Fx.Px.Sx.G motif
pHMM hits without GxGxxA and Fx.Px.Sx.G motif

### Flavin-dependent (conventional)

| | |
|---|---|
| Job Number | 157098 |
| Time Started -- Finished | 7/7 03:29 AM -- 7/7 03:33 AM |
| Database Version | UniProt: 2025-02 / InterPro: 105 |
| Input Option | FASTA (Option C), no FASTA header reading |
| Job Name | FDH_conventional |
| E-Value for SSN Edge Calculation | 5 |
| Uploaded FASTA File | fdh_conventional_ssn.fasta |
| Number of Sequences in Uploaded File | 1,960 |
| Exclude Fragments | No |
| Total Number of Sequences in Dataset | 1,960 |
| Total Number of Edges | 751,781 |
| Number of Unique Sequences | 1,960 |
| Convergence Ratio? | 0.392 |

Alignment Score threshold: 100

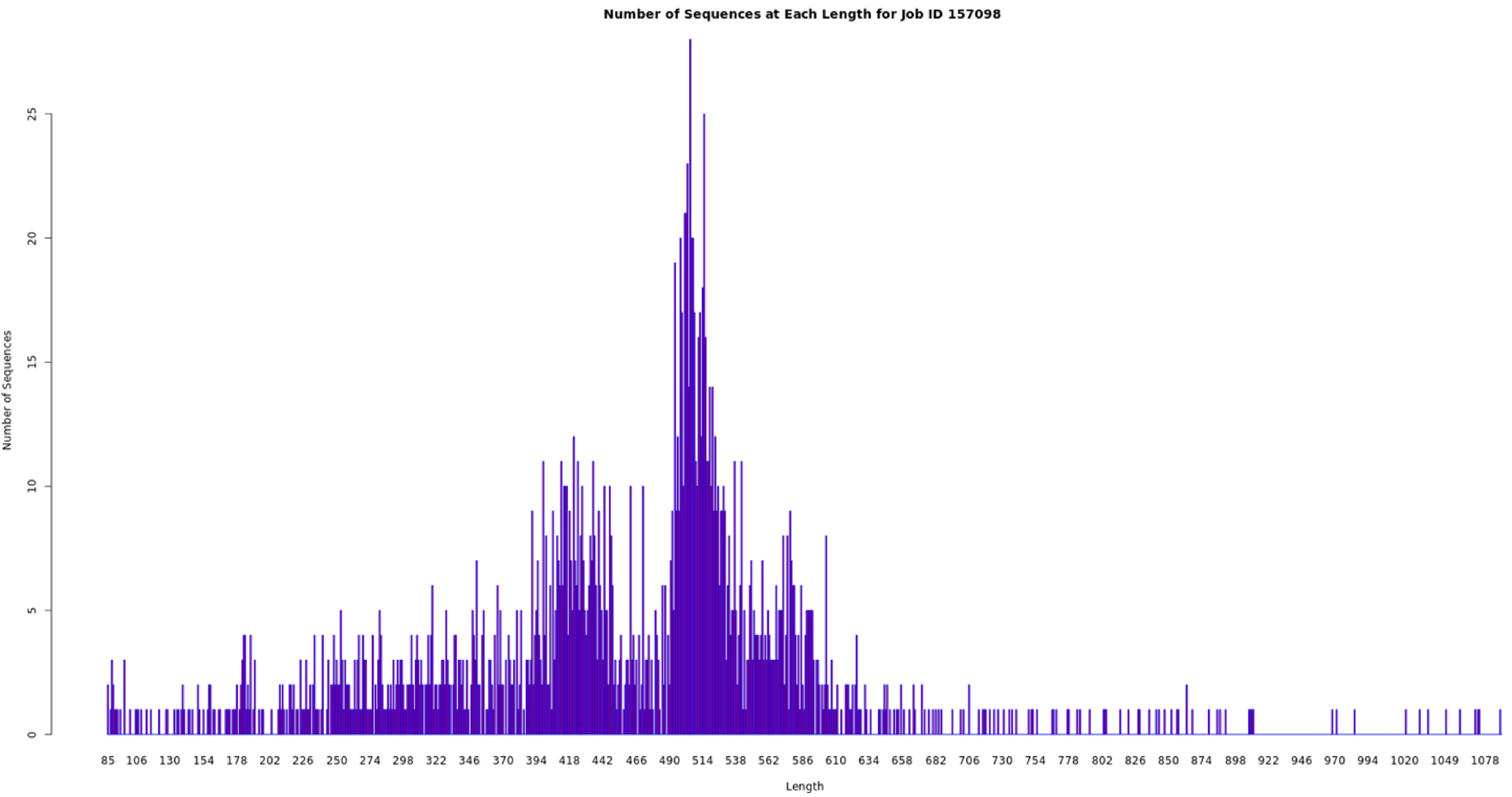
Number of Sequences at Each Length for Job ID 157098
Number of Sequences
0
5
10
15
20
25
Length
85 106 130 154 178 202 226 250 274 298 322 346 370 394 418 442 466 490 514 538 562 586 610 634 658 682 706 730 754 778 802 826 850 874 898 922 946 970 994 1020 1049 1078

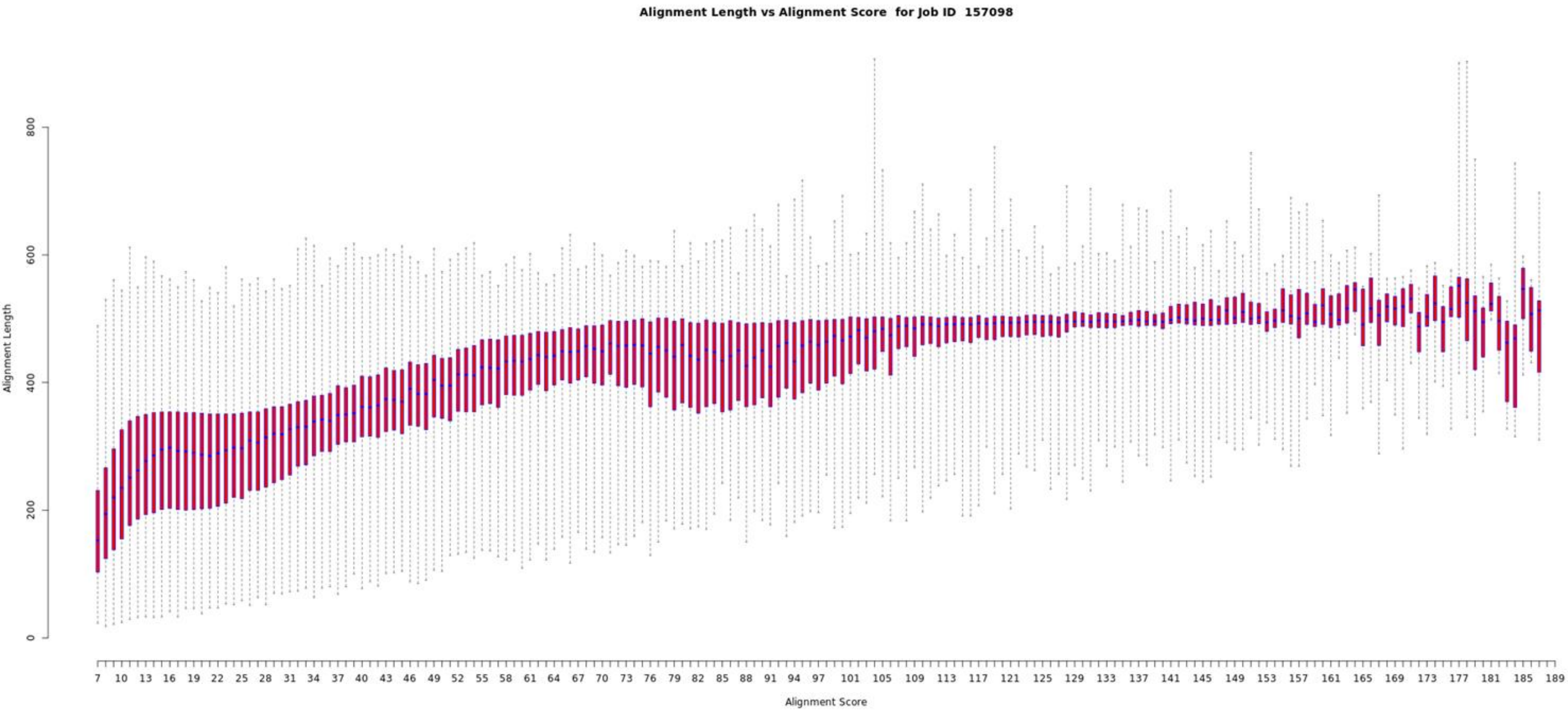
Alignment Length vs Alignment Score for Job ID 157098
Alignment Length
0
200
400
600
800
Alignment Score
7 10 13 16 19 22 25 28 31 34 37 40 43 46 49 52 55 58 61 64 67 70 73 76 79 82 85 88 91 94 97 101 105 109 113 117 121 125 129 133 137 141 145 149 153 157 161 165 169 173 177 181 185 189

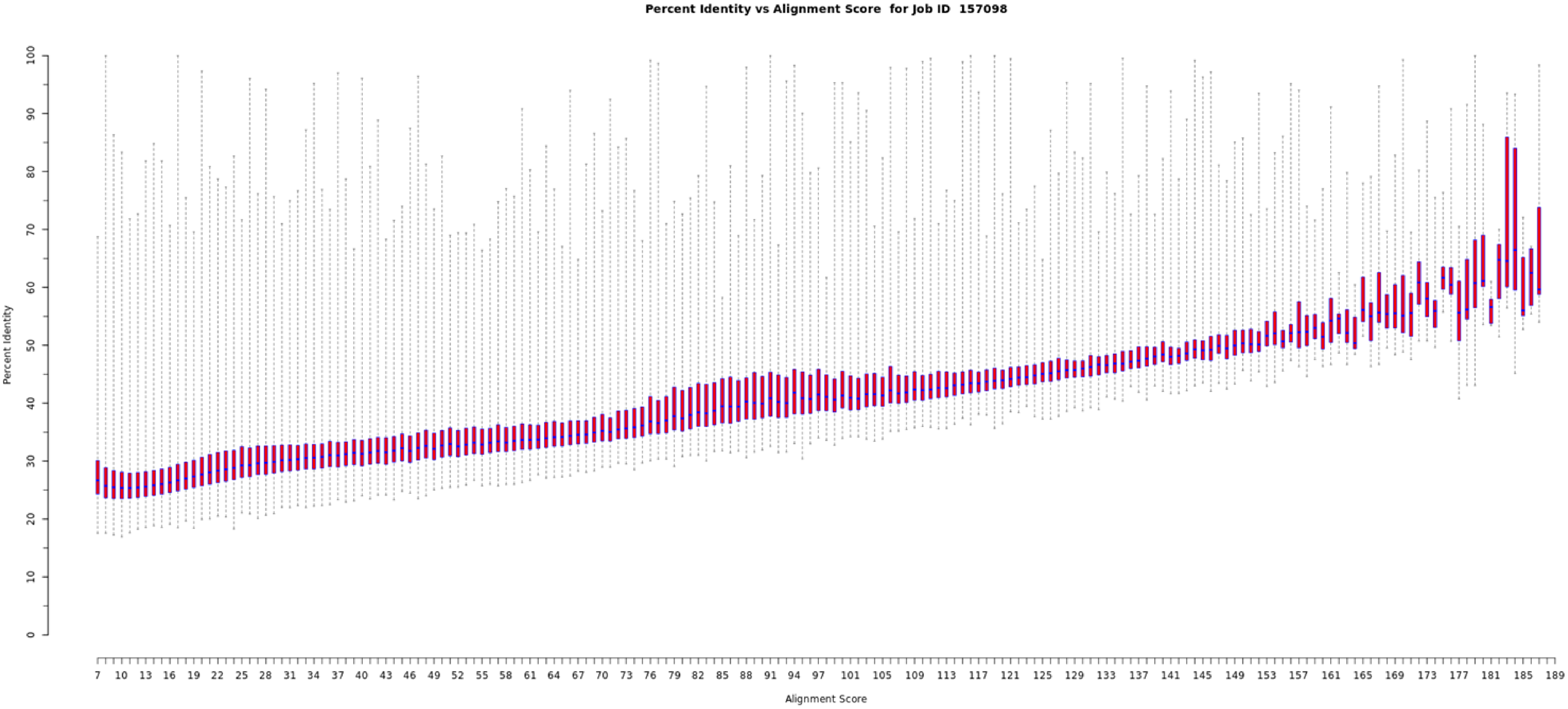
Percent Identity vs Alignment Score for Job ID 157098
Percent Identity
Alignment Score
0
10
20
30
40
50
60
70
80
90
100
7 10 13 16 19 22 25 28 31 34 37 40 43 46 49 52 55 58 61 64 67 70 73 76 79 82 85 88 91 94 97 101 105 109 113 117 121 125 129 133 137 141 145 149 153 157 161 165 169 173 177 181 185 189

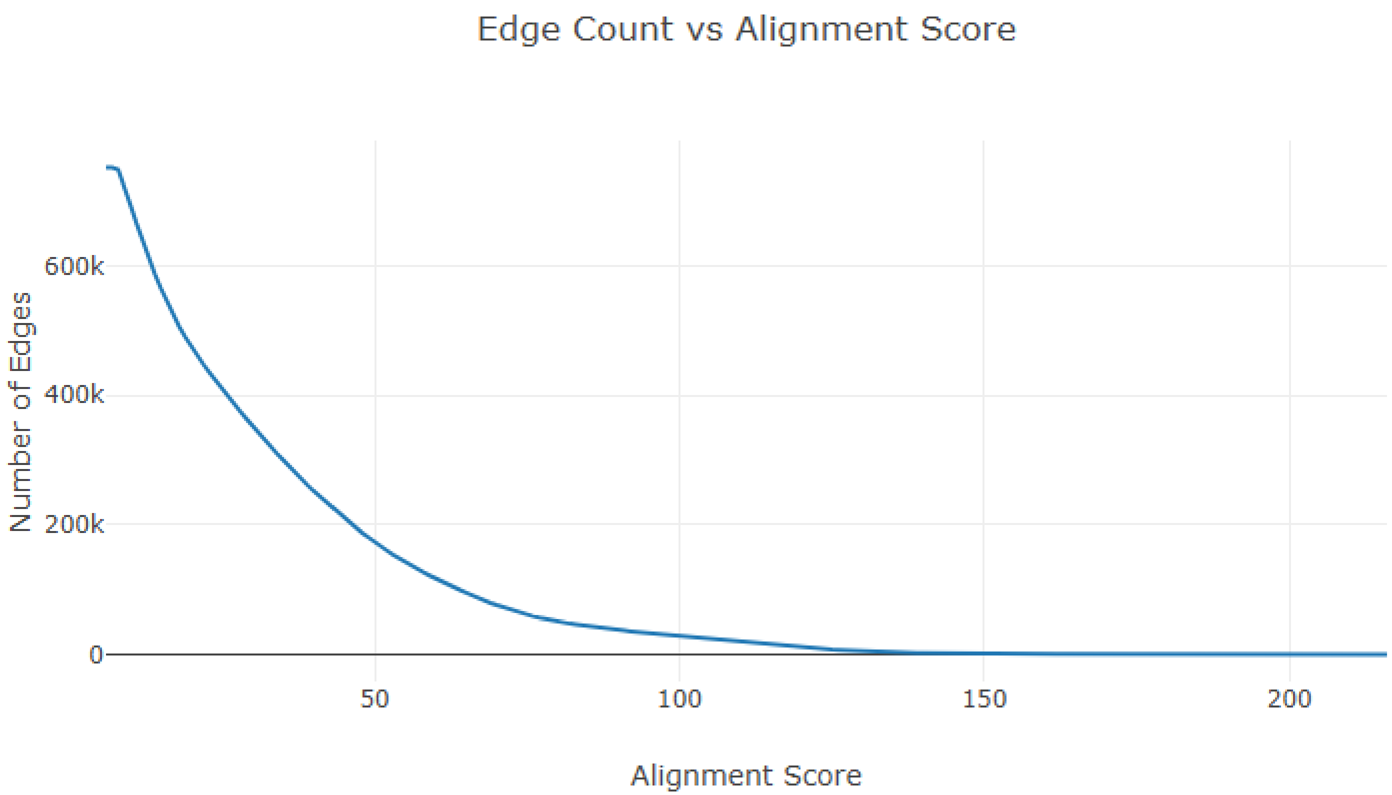
Edge Count vs Alignment Score
Number of Edges
600k
400k
200k
0
50
100
150
200
Alignment Score

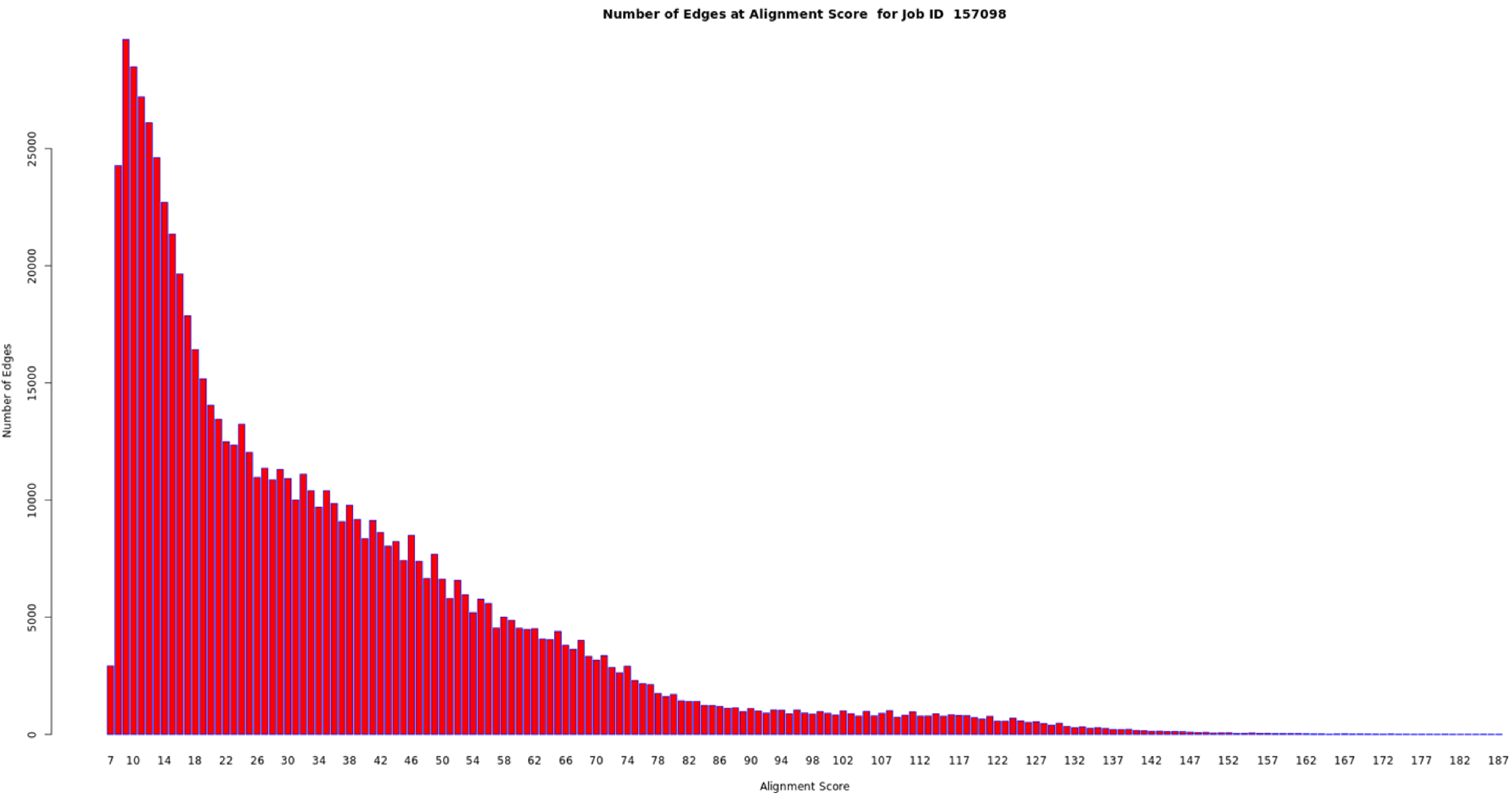
Number of Edges at Alignment Score for Job ID 157098
Number of Edges
0
5000
10000
15000
20000
25000
7 10 14 18 22 26 30 34 38 42 46 50 54 58 62 66 70 74 78 82 86 90 94 98 102 107 112 117 122 127 132 137 142 147 152 157 162 167 172 177 182 187
Alignment Score

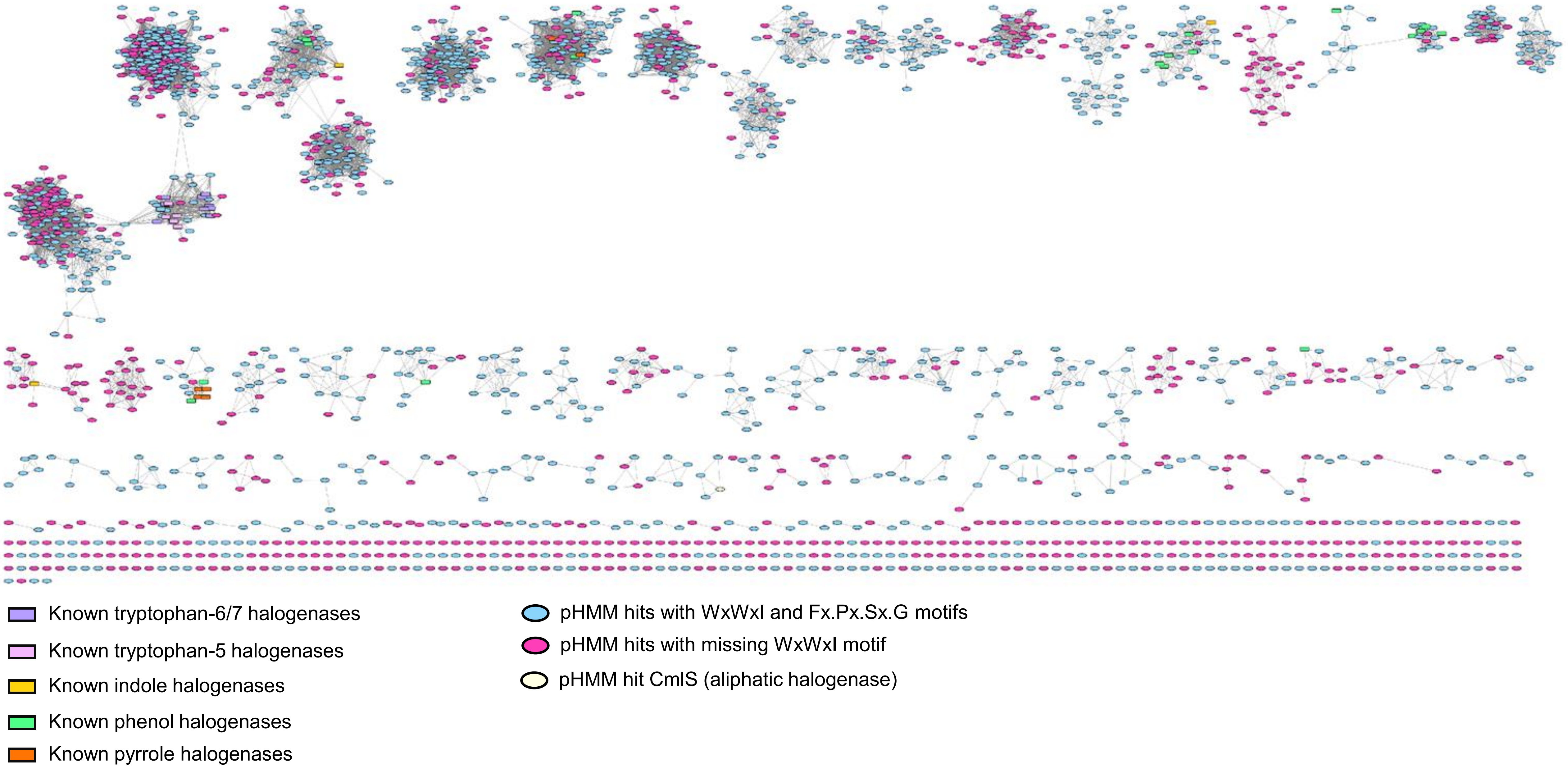
Known tryptophan-6/7 halogenases
Known tryptophan-5 halogenases
Known indole halogenases
Known phenol halogenases
Known pyrrole halogenases
pHMM hits with WxWxI and Fx.Px.Sx.G motifs
pHMM hits with missing WxWxI motif
pHMM hit CmlS (aliphatic halogenase)

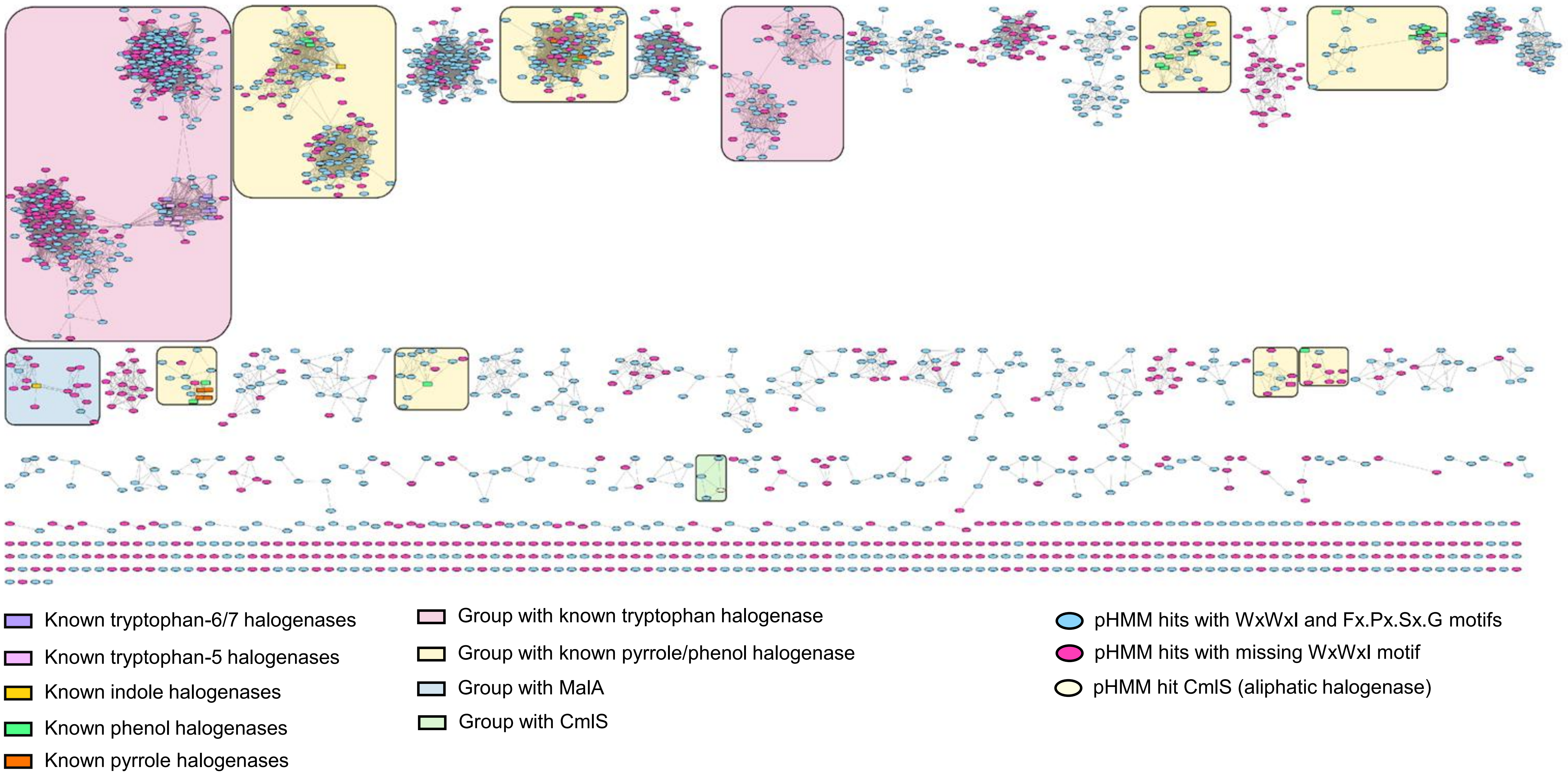
Known tryptophan-6/7 halogenases
Known tryptophan-5 halogenases
Known indole halogenases
Known phenol halogenases
Known pyrrole halogenases
Group with known tryptophan halogenase
Group with known pyrrole/phenol halogenase
Group with MalA
Group with CmlS
pHMM hits with WxWxI and Fx.Px.Sx.G motifs
pHMM hits with missing WxWxI motif
pHMM hit CmlS (aliphatic halogenase)

### Non-heme iron-dependent

| | |
|---|---|
| Job Number | 157360 |
| Time Started -- Finished | 7/11 12:05 AM -- 7/11 12:08 AM |
| Database Version | UniProt: 2025-03 / InterPro: 106 |
| Input Option | FASTA (Option C), no FASTA header reading |
| Job Name | NHFe |
| E-Value for SSN Edge Calculation | 5 |
| Uploaded FASTA File | nhfe_final_ssn.fasta |
| Number of Sequences in Uploaded File | 1,134 |
| Exclude Fragments | No |
| Total Number of Sequences in Dataset | 1,134 |
| Total Number of Edges | 177,036 |
| Number of Unique Sequences | 1,134 |
| Convergence Ratio? | 0.276 |

Alignment Score threshold: 45

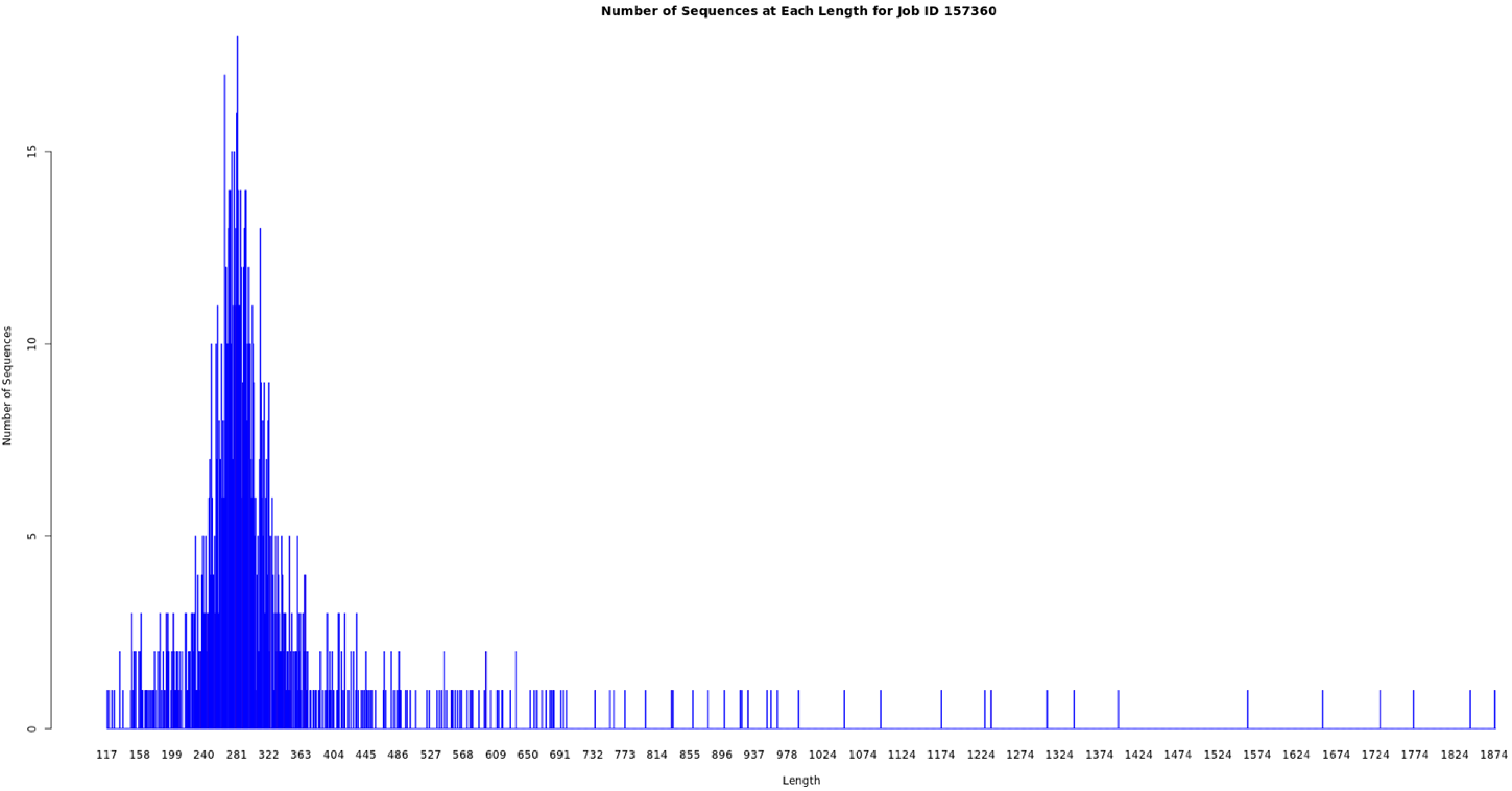

Number of Sequences at Each Length for Job ID 157360
Number of Sequences
0
5
10
15
117
158
199
240
281
322
363
404
445
486
527
568
609
650
691
732
773
814
855
896
937
978
1024
1074
1124
1174
1224
1274
1324
1374
1424
1474
1524
1574
1624
1674
1724
1774
1824
1874
Length

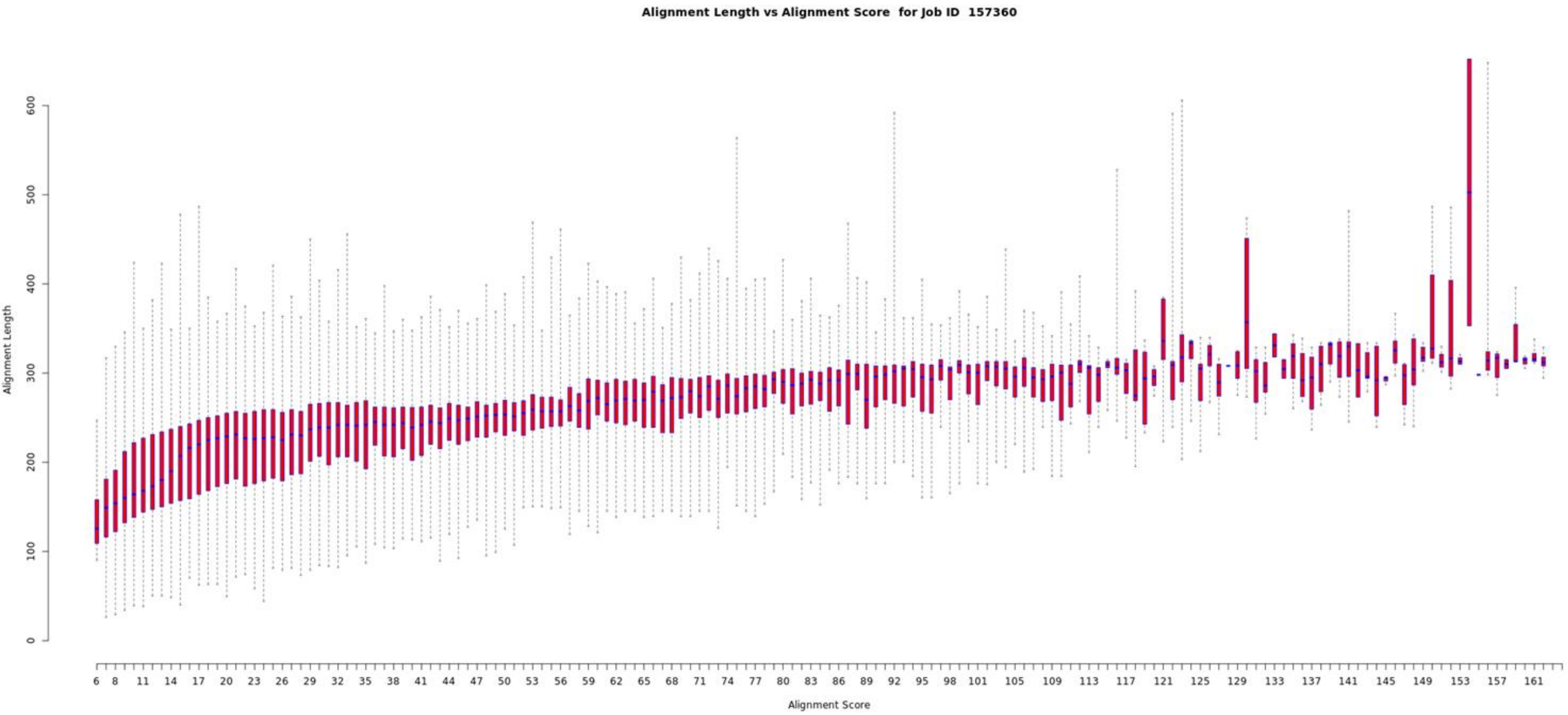

Alignment Length vs Alignment Score for Job ID 157360
Alignment Length
0
100
200
300
400
500
600
Alignment Score
6 8 11 14 17 20 23 26 29 32 35 38 41 44 47 50 53 56 59 62 65 68 71 74 77 80 83 86 89 92 95 98 101 105 109 113 117 121 125 129 133 137 141 145 149 153 157 161

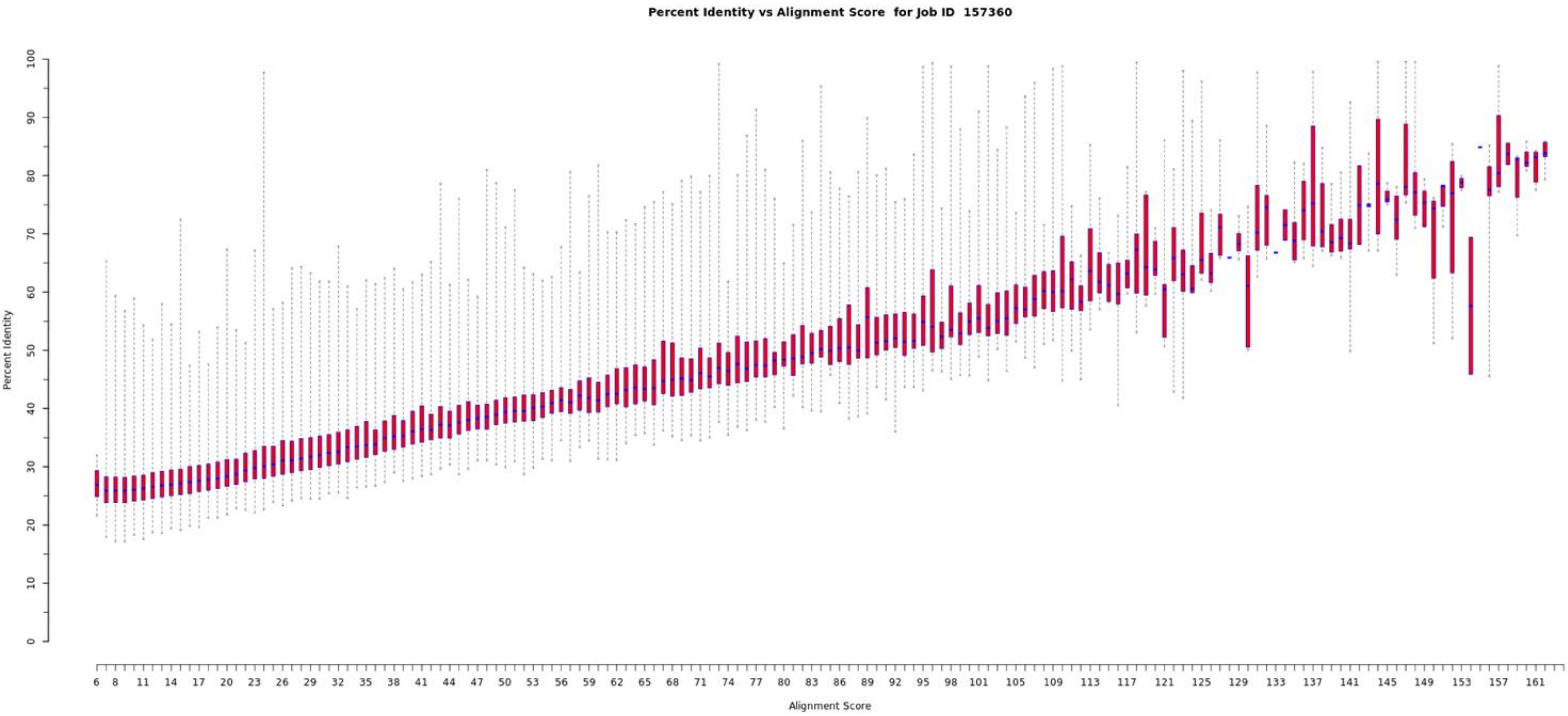

Percent Identity vs Alignment Score for Job ID 157360
Percent Identity
0
10
20
30
40
50
60
70
80
90
100
Alignment Score
6 8 11 14 17 20 23 26 29 32 35 38 41 44 47 50 53 56 59 62 65 68 71 74 77 80 83 86 89 92 95 98 101 105 109 113 117 121 125 129 133 137 141 145 149 153 157 161

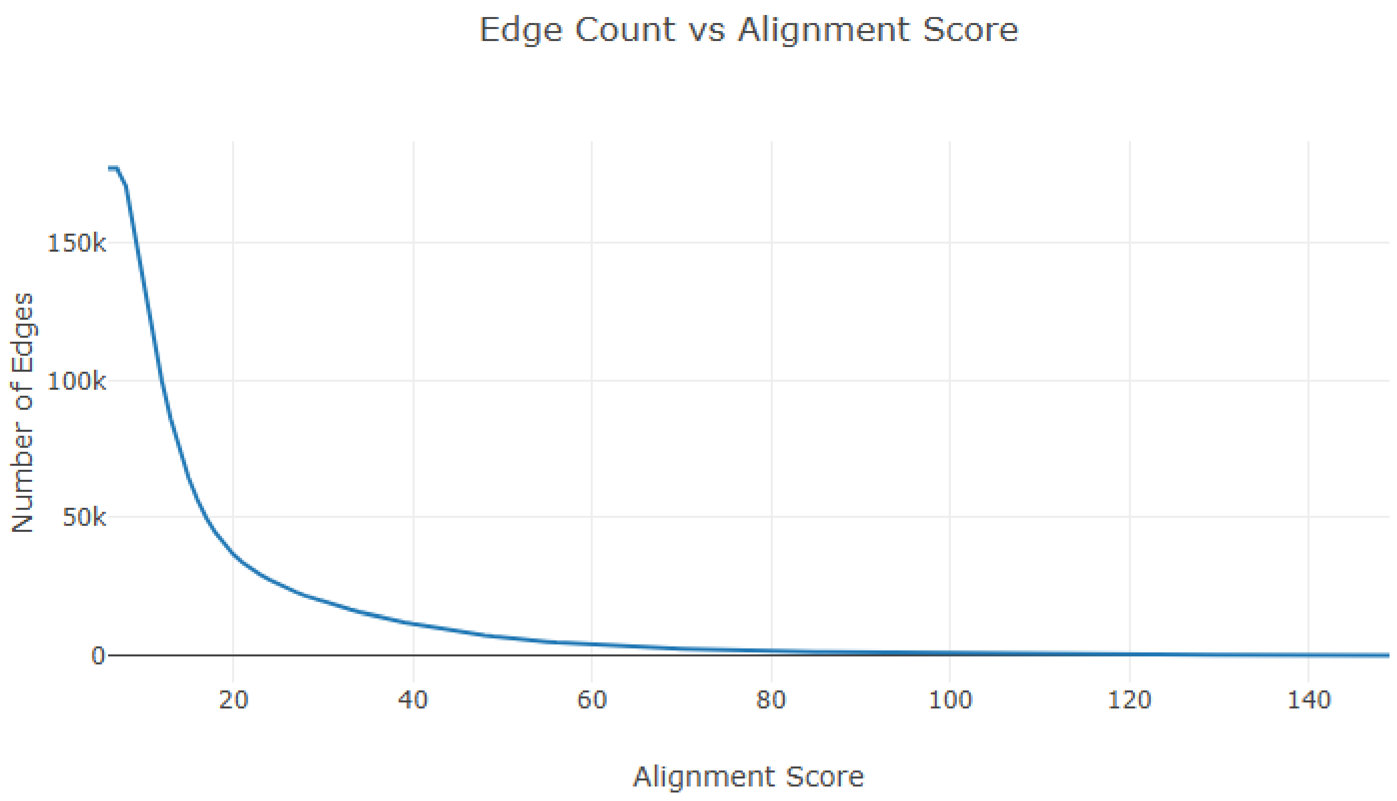
Edge Count vs Alignment Score
Number of Edges
150k
100k
50k
0
20
40
60
80
100
120
140
Alignment Score

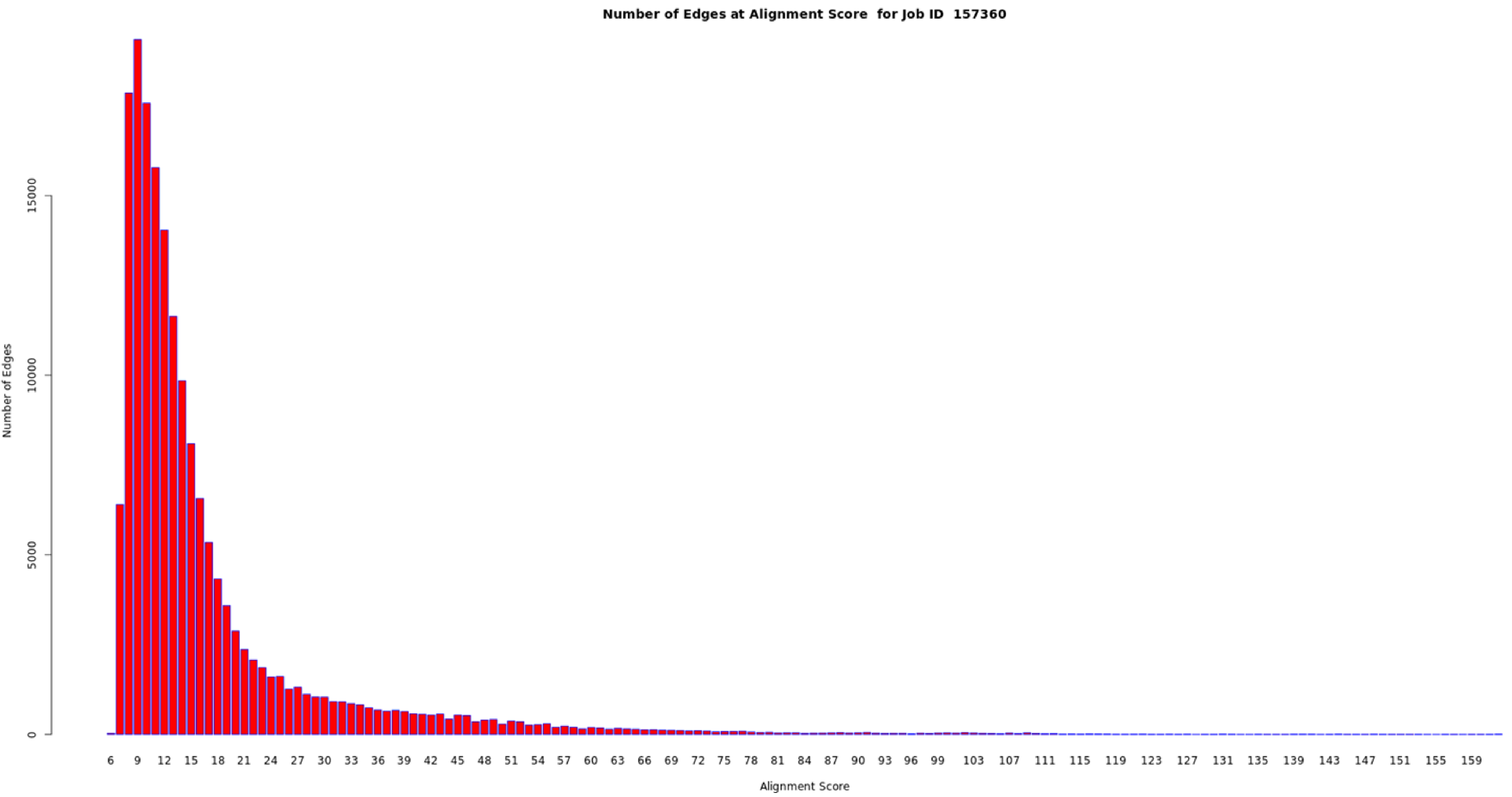

Number of Edges at Alignment Score for Job ID 157360
Number of Edges
0
5000
10000
15000
6 9 12 15 18 21 24 27 30 33 36 39 42 45 48 51 54 57 60 63 66 69 72 75 78 81 84 87 90 93 96 99 103 107 111 115 119 123 127 131 135 139 143 147 151 155 159
Alignment Score

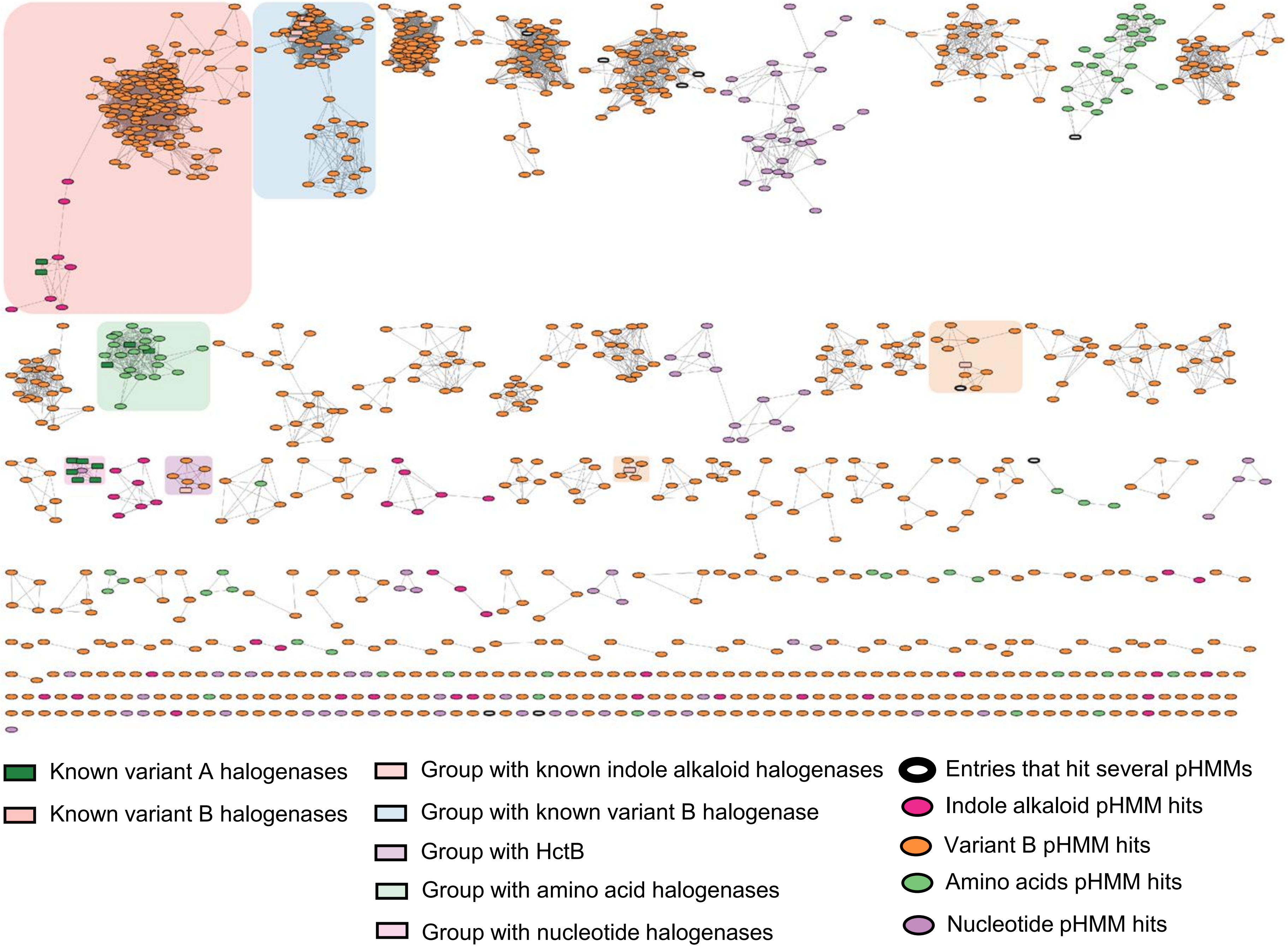

Known variant A halogenases
Known variant B halogenases
Group with known indole alkaloid halogenases
Group with known variant B halogenase
Group with HctB
Group with amino acid halogenases
Group with nucleotide halogenases
Entries that hit several pHMMs
Indole alkaloid pHMM hits
Variant B pHMM hits
Amino acids pHMM hits
Nucleotide pHMM hits

### Copper-dependent

| | |
|---|---|
| Job Number | 168369 |
| Time Started -- Finished | 12/22 09:34 AM -- 12/22 09:42 AM |
| Database Version | UniProt: 2025-03 / InterPro: 106 |
| Input Option | FASTA (Option C), no FASTA header reading |
| Job Name | copper-dependent |
| E-Value for SSN Edge Calculation | 5 |
| Uploaded FASTA File | copper-dependent_hits.fasta |
| Number of Sequences in Uploaded File | 5,299 |
| Exclude Fragments | No |
| Total Number of Sequences in Dataset | 5,299 |
| Total Number of Edges | 6,040,909 |
| Number of Unique Sequences | 5,299 |
| Convergence Ratio? | 0.430 |

Alignment Score threshold: 50

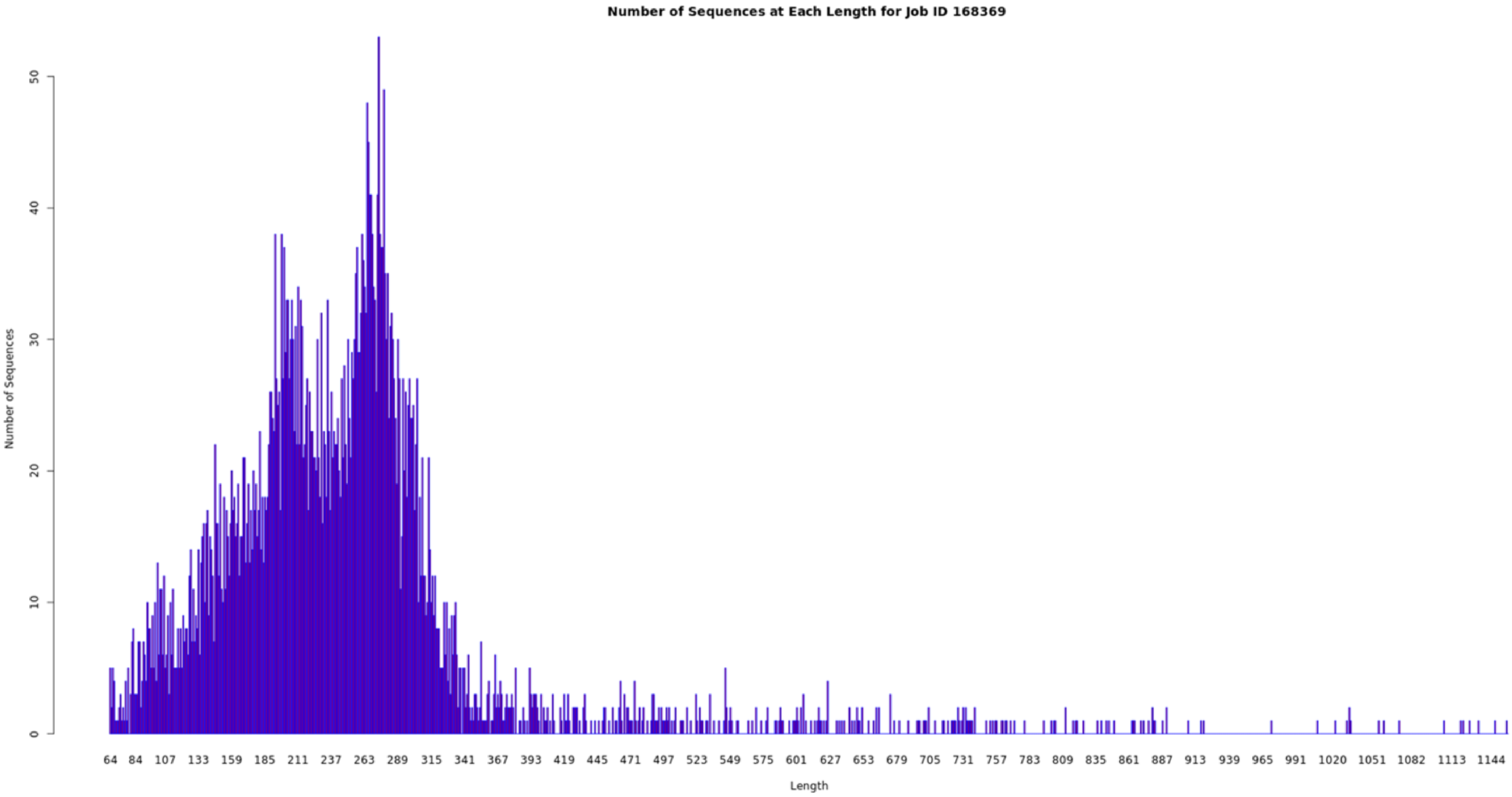

Number of Sequences at Each Length for Job ID 168369
Number of Sequences
0
10
20
30
40
50
64 84 107 133 159 185 211 237 263 289 315 341 367 393 419 445 471 497 523 549 575 601 627 653 679 705 731 757 783 809 835 861 887 913 939 965 991 1020 1051 1082 1113 1144
Length

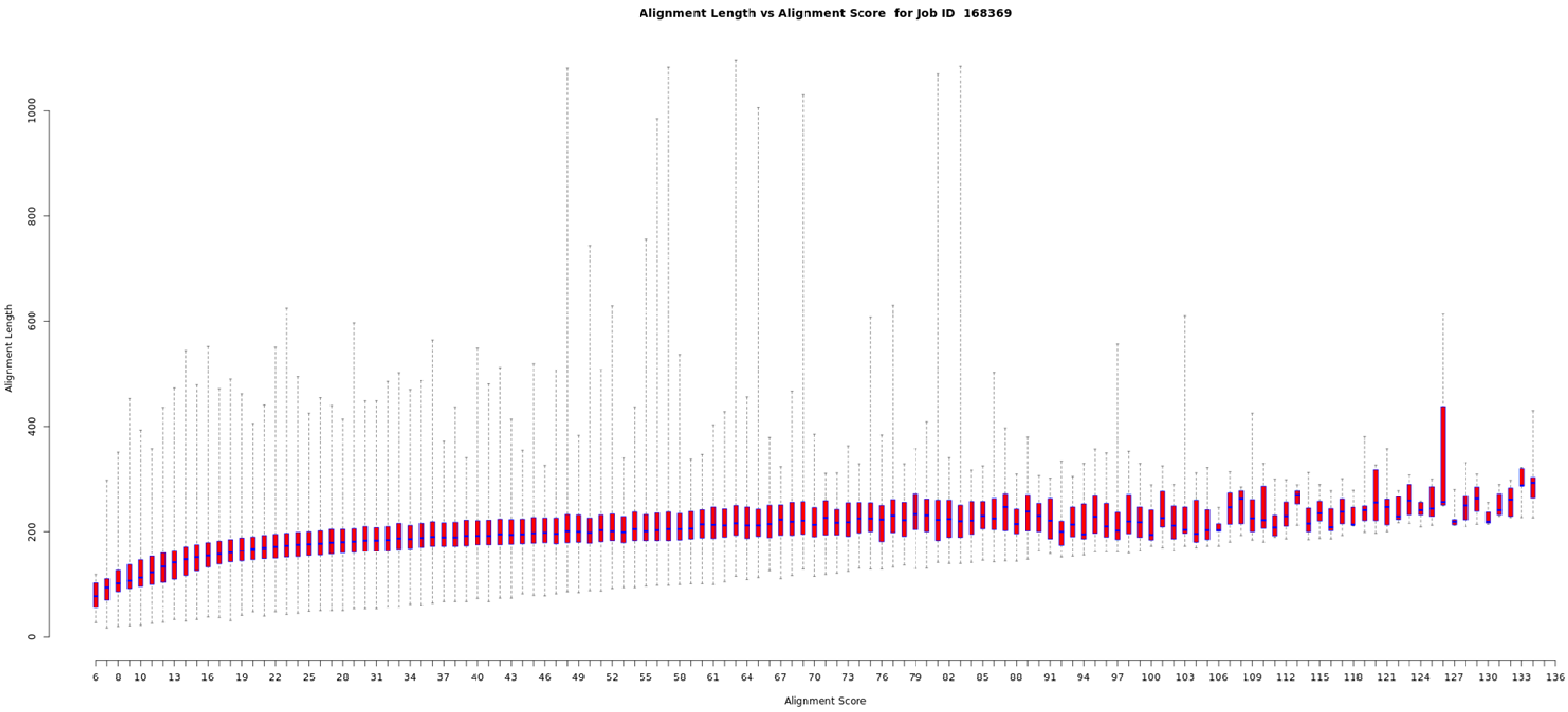
Alignment Length vs Alignment Score for Job ID 168369
Alignment Length
0
200
400
600
800
1000
6
8
10
13
16
19
22
25
28
31
34
37
40
43
46
49
52
55
58
61
64
67
70
73
76
79
82
85
88
91
94
97
100
103
106
109
112
115
118
121
124
127
130
133
136
Alignment Score

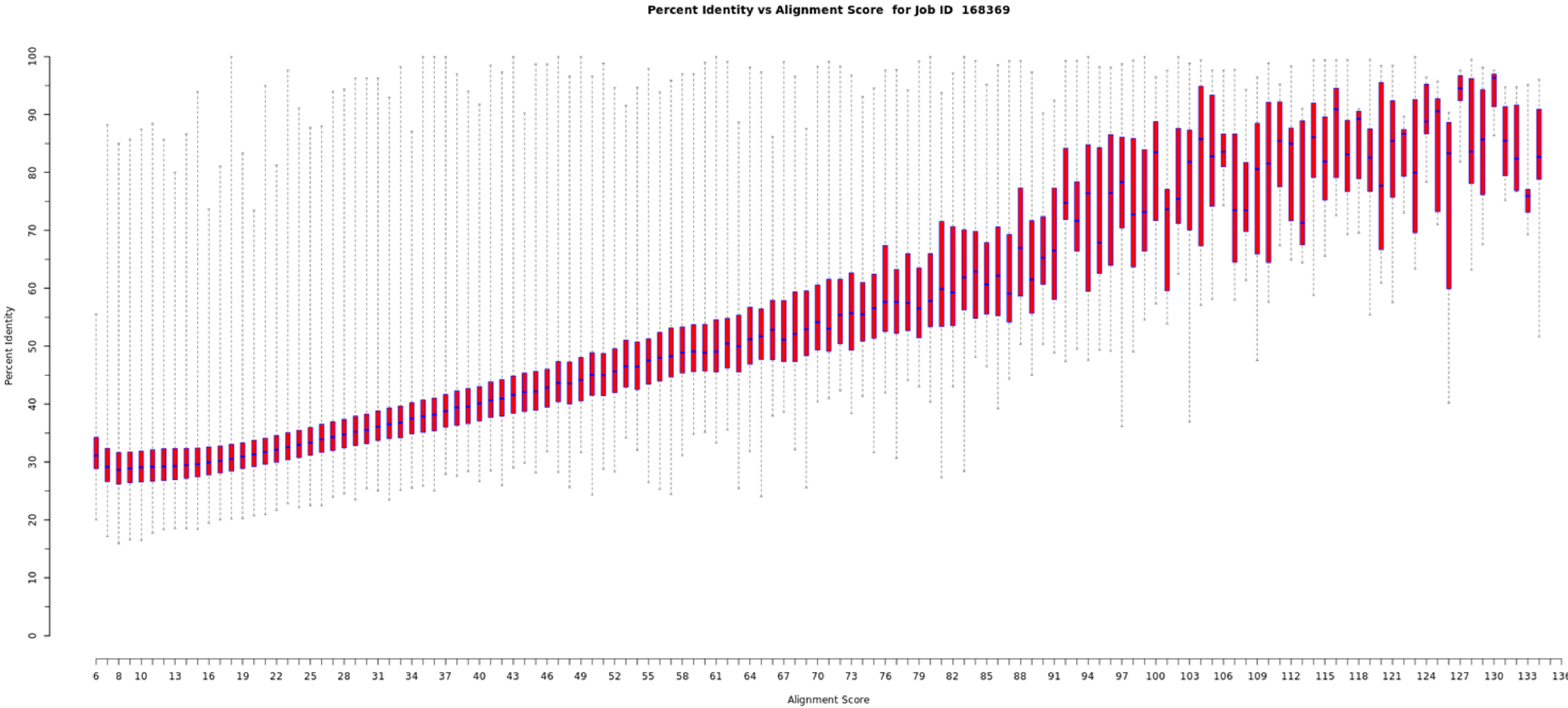

Percent Identity vs Alignment Score for Job ID 168369
Percent Identity
Alignment Score
0
10
20
30
40
50
60
70
80
90
100
6
8
10
13
16
19
22
25
28
31
34
37
40
43
46
49
52
55
58
61
64
67
70
73
76
79
82
85
88
91
94
97
100
103
106
109
112
115
118
121
124
127
130
133
136

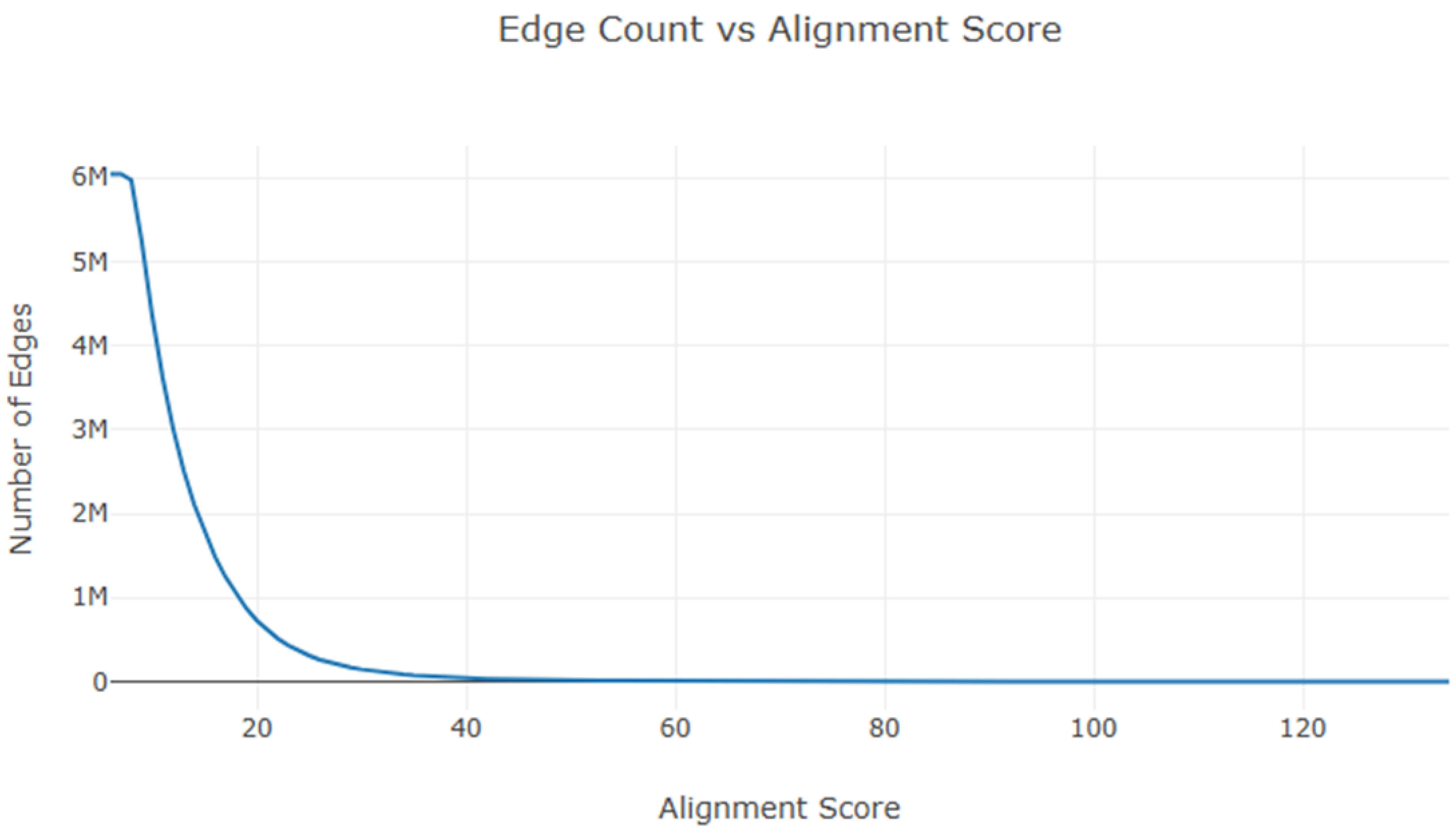

Edge Count vs Alignment Score
Number of Edges
6M
5M
4M
3M
2M
1M
0
20
40
60
80
100
120
Alignment Score

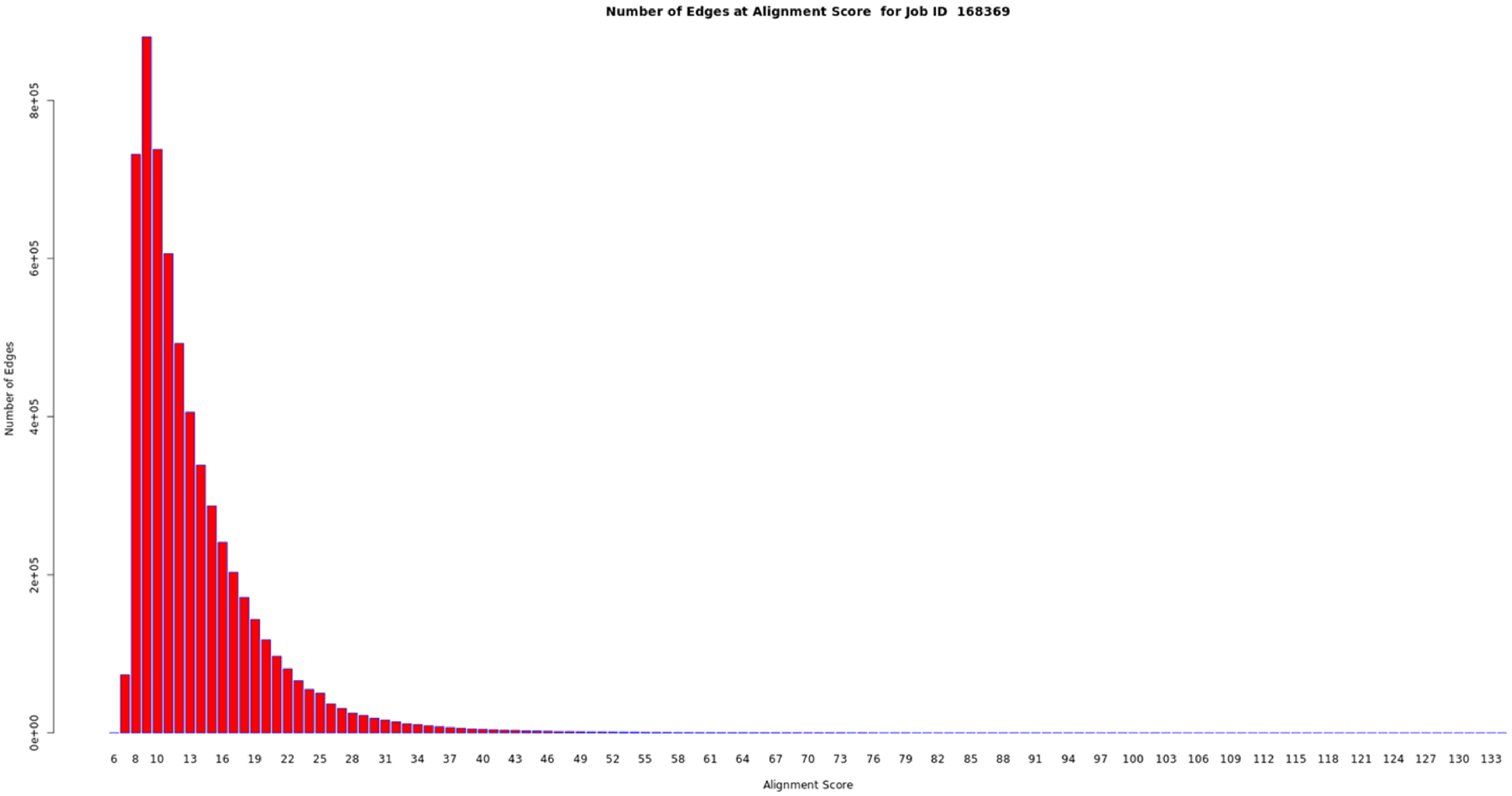

Number of Edges at Alignment Score for Job ID 168369
Number of Edges
0e+00
2e+05
4e+05
6e+05
8e+05
6 8 10 13 16 19 22 25 28 31 34 37 40 43 46 49 52 55 58 61 64 67 70 73 76 79 82 85 88 91 94 97 100 103 106 109 112 115 118 121 124 127 130 133
Alignment Score

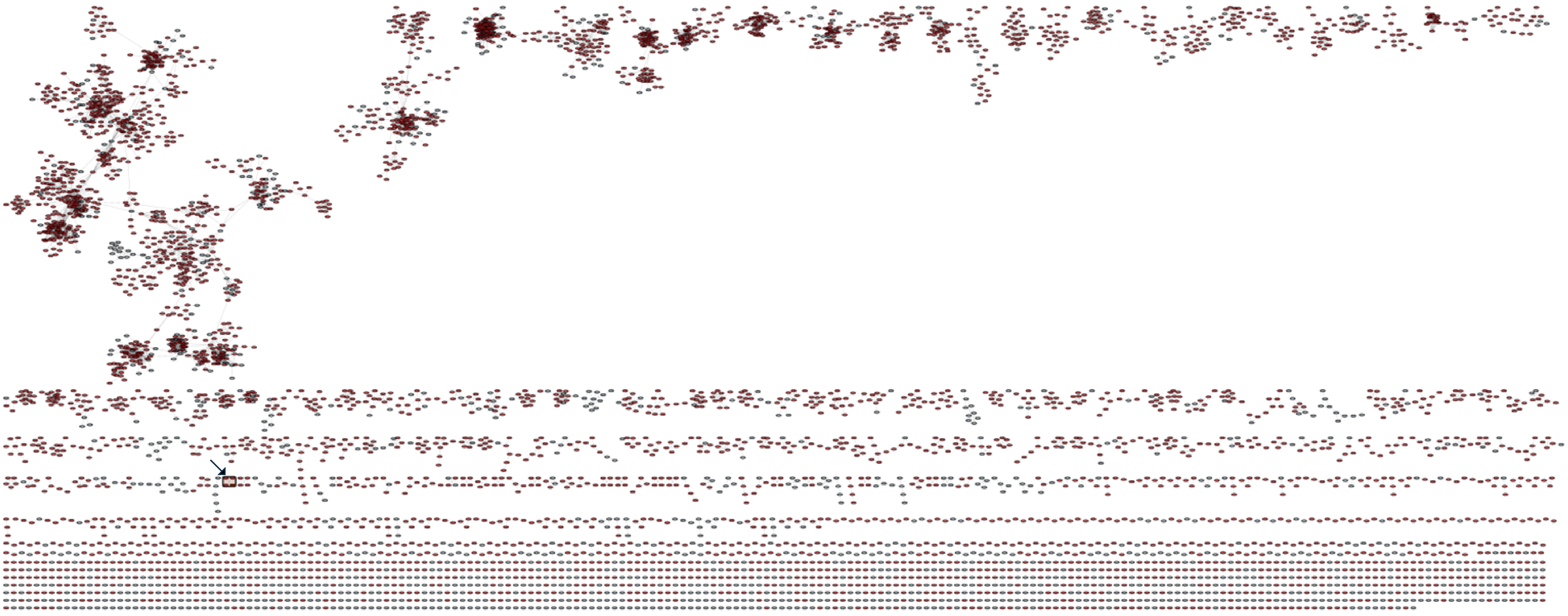

Enzymes with two HxxHC motifs

Enzymes not containing two HxxHC motifs

Cluster with ApnU

### Workflow

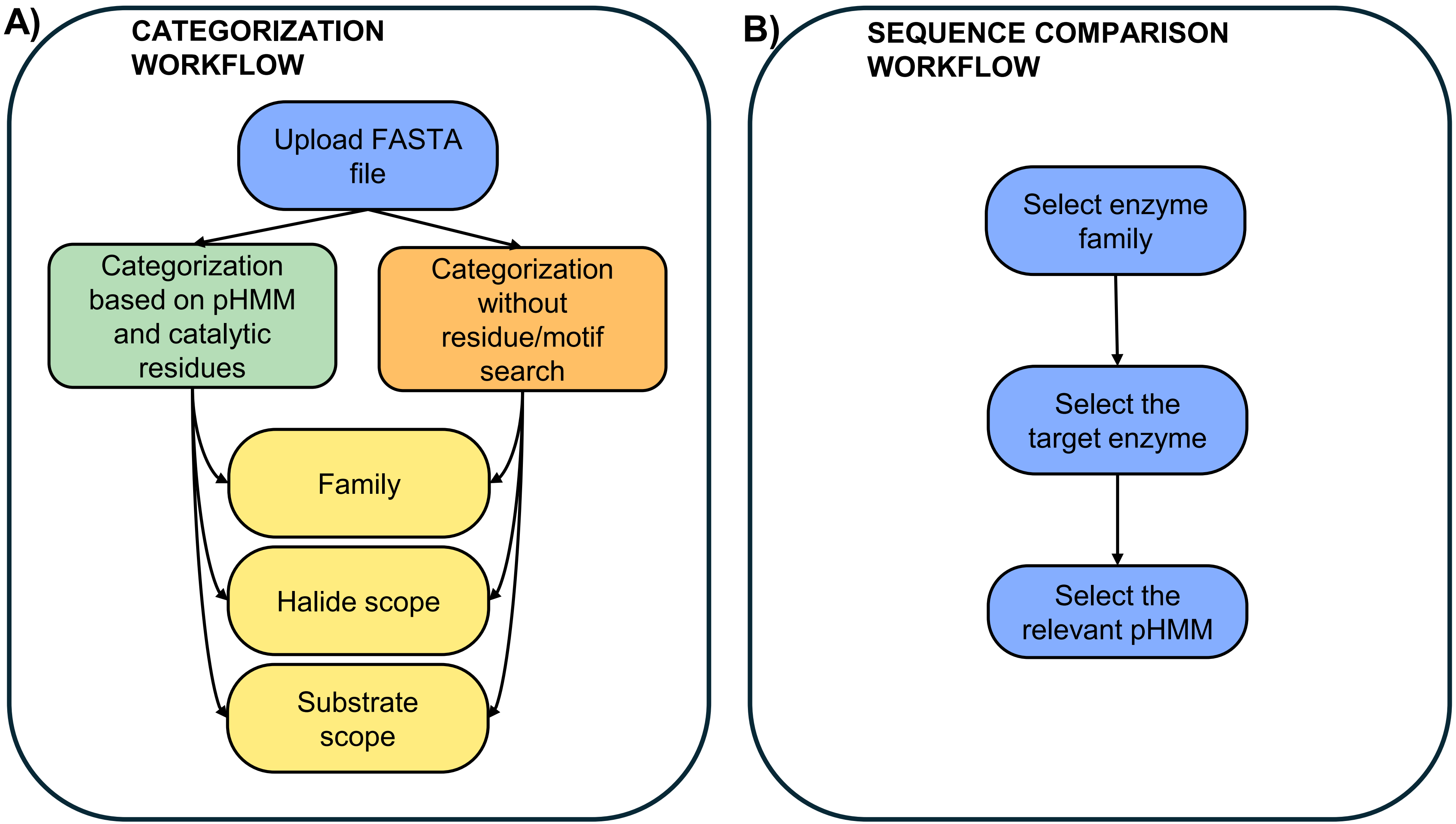
A)
CATEGORIZATION WORKFLOW
Upload FASTA file
Categorization based on pHMM and catalytic residues
Categorization without residue/motif search
Family
Halide scope
Substrate scope
B)
SEQUENCE COMPARISON WORKFLOW
Select enzyme family
Select the target enzyme
Select the relevant pHMM

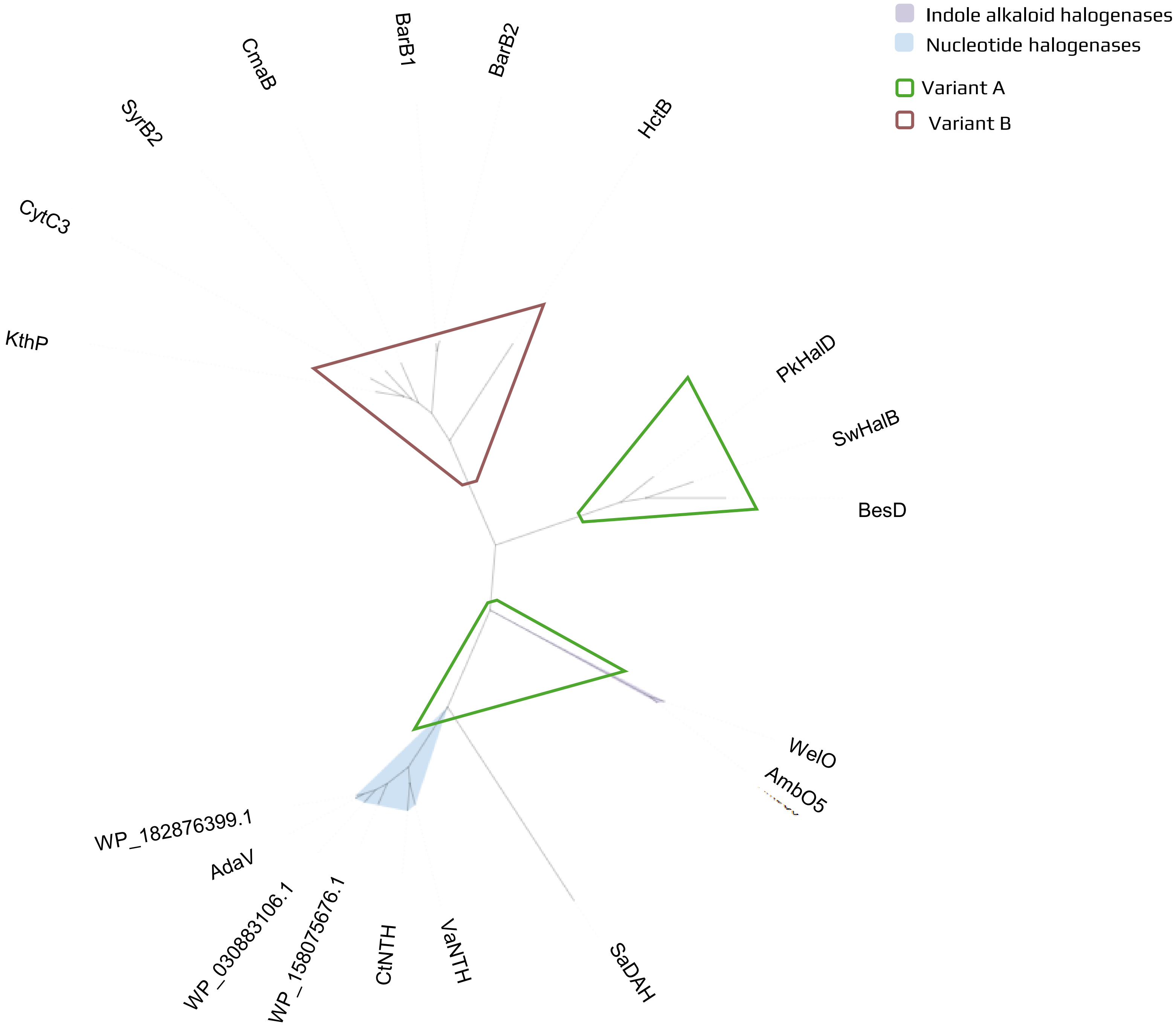

Indole alkaloid halogenases
Nucleotide halogenases
Variant A
Variant B
CmaB
BarB1
BarB2
SyrB2
HctB
CytC3
KthP
PkHalD
SwHalB
BesD
WelO
AmbO5
WP_182876399.1
AdaV
WP_030883106.1
WP_158075676.1
CtNTH
VaNTH
SaDAH
