## Supplemental Information for "Computational pipeline reveals nature’s untapped reservoir of halogenating enzymes"

**RECOMMENDED pHMM CUTOFFS**

| Enzyme Family | pHMM | Recommended cutoff | MSA (for building the pHMM) |
|---|---|---|---|
| **SAM-dependent** | Non-fluorinases | 250 | Whole sequence |
| | Fluorinases | 150 | Whole sequence |
| **NHFe-dependent** | Indole alkaloid (variant A) | 600 | Whole sequence |
| | Nucleotide (variant A) | 200 | Cut |
| | Small amino acids (variant A) | | Whole sequence |
| | Variant B | 200 | Whole sequence |
| **Dimetal-carboxylate** | General | No cutoff | Whole sequence |
| **Flavin-dependent** | General | No cutoff | Cut |
| | Tryptophan-5 | 350 (for mibH-like sequences); 850 | Whole sequence |
| | Tryptophan-6/7 | 750 | Whole sequence |
| | Orsellinic acid-like | 500 | Whole sequence |
| | Tyrosine/Hpg-like | 305 | Cut |
| | Pyrrolic | 400 | Cut |
| **Vanadium-dependent** | General | | Whole sequence |
| | Selective chloroperoxidases | 600 | Cut |
| | Non-selective chloroperoxidases | 700 | Cut |
| | Bromoperoxidases | 500 | Whole sequence |
| | Iodoperoxidases | 700 | Whole sequence |
| **Copper-dependent** | General | No cutoff | Whole sequence |

***Supplementary Table 1 Recommended pHMM cutoffs***

The cutoffs can be used to help identify close homologs and conventional sequences. The thresholds were defined by crossvalidation and by testing the closest homologs with different activity, substrate, regio-or substrate-selectivity against the target pHMM. The general pHMMs which are used to detect members of the given family without substrate or detailed activity prediction do not have recommended cutoffs and we suggest to rely on the conserved motifs and catalytic residues solely.

Database implemented in Geneious Prime 2024.03.25 searched by InterProScan: CDD, Coils, Gene3d, HAMAP, MobiDBLite, Panther, PfamA, Phobius, PIRSF, PRINTS, PrositePatterns, PrositeProfiles, SFLD, SignalP, SignalP_EUK, SignalP_GRAM_NEGATIVE, SignalP_GRAM_POSITIVE, SMART, SuperFamily, NCBIfam, TMHMM

**GUIDE FOR THE RESIDUE AND MOTIF COLLECTION**

**Radical substitution**

**Non-heme iron alphaketoglutarate-dependent halogenases**

**Sequence motifs.** Non-heme Iron alphaketoglutarate (NHFe)-dependent halogenases carry out nucleophilic substitution and are shown to act on aliphatic compounds (The Role of Chloride in the Mechanism...; Hillwig and Liu 2014; Song et al. 2019). Differentiating them from their hydroxylase and dioxygenase close homologs is supported by the catalytic triad. We see HXD/E in hydroxylases and dioxygenases, and HXG/A in halogenases (Neugebauer et al. 2019). It is confirmed by experiments that the single residue is the key for switching between halogenation and hydroxylation reactions (Blasiak et al. 2006; Kim et al. 2020). Although not part of the catalytic triad, Ser189 in WelO5 (nucleotide halogenase) seems to be the key to the halogenation reaction over hydroxylation (Zhang et al. 2020). This information is not generalizable to the whole family, as the halogenation activity of a substrate-and stereo-selective enzyme from 2'-chloropentostatin biosynthesis (Zhai et al. 2022), AdaV is due to Glu196, Asn203 and Val269 (Zhai et al. 2022).
**Activity.** As seen for flavin-dependent halogenases (FDHs), they can also be placed into variant A and variant B groups. Variant A enzymes acting on free standing substrates and variant B enzymes acting on a substrate tethered to an acyl carrier protein (ACP).In some cases the ACPs locate spatially close to the halogenase, like in the instance of HctB (halogenase from hectochlorin biosynthesis), where they form a multi-enzyme complex (Pratter et al. 2014), and therefore we can hypothesize that the halogenase belongs in the variant B group. Although in most cases neither the sequence nor the structural motifs are known for predicting the mechanism of action, enzymes in both groups have several described residues that can guide researchers to predict their substrate-specificity and chemoselectivity, but not their variant A or variant B nature.
Indole alkaloid halogenases, such as WelO5 or AmbO5, act on free-standing substrates (Hillwig and Liu 2014). Many small, aliphatic amino acid accepting enzymes, e.g., SwHalB, PkHalD, and BesD, and the nucleotide decorating enzymes like AdaV (acting on dAMP), CtNTH, and VaNTH (acting on dGMP) also belong to this group (Hillwig and Liu 2014; Ni et al. 2024; Zhai et al. 2022).
Among the variant B group, we will find CmaB from cyclopropyl biosynthesis (Vaillancourt et al. 2005), KtzD from kutzneride biosynthesis (Biosynthesis of (−)-(1S,2R)-Allocoron...), SyrB2 from syringomycin biosynthesis (Matthews et al. 2009), BarB1/2 from bartoloside biosynthesis (Galonić et al. 2006), and CytC3 from soil *Streptomyces* sp. (Ueki et al. 2006) acting on amino acids. Two enzymes showing cryptic halogenation reactions, CurA from curamicin biosynthesis and JamE from jamaicamide biosynthesis (Khare et al. 2010; Gu et al. 2009), are also in this group. A fatty acid halogenase, HctB from the hectochlorin biosynthesis, acts on the substrate tethered to a carrier protein as well (The Role of Chloride in the Mechanism...). In-depth structural characterization of the fatty acid acting halogenase HctB (Pratter et al. 2014) and the enzyme decorating piperazyl (KthP) (Jiang et al. 2011) is missing, making it difficult to develop substrate-specific annotation pipelines for fatty acid and piperazyl-acting halogenases in this family.

**Dimetal-carboxylate halogenases**

**Sequence motifs.** Dimetal-carboxylate halogenases are a group described in 2017 (Wang et al. 2025). There were no established practices for mining for them. However, we might have some support to differentiate them from the diiron oxygenases, e.g., N-oxygenase AurF (Simurdiak et al. 2006). The putative nine catalytic residues (Glu103, Gln135, Glu136, His139, Glu283, His310, Asp313, Glu314, His317) are all necessary for chlorination activity (Wang et al. 2025). Furthermore, Asp282 and Glu252 may affect catalytic activity without having an impact on the metal acquisition (Wang et al. 2025). Therefore, both two motifs that contain the mentioned catalytic residues are relied on when looking for halogenases in this group. AurF homologs can be differentiated from the halogenase members as it contains Asp135 in place of Gln135 (Wang et al. 2025). Therefore, two small motifs, QExxH and HxxDExxH, can be seen in genome mining practices to recognize CylC and its homologs (Wang et al. 2025).
**Activity.** Dimetal-carboxylate halogenases are a predicted group that received experimental support for their diiron cofactor and catalytic mechanism recently (Wang et al. 2025). The first characterized enzyme is CylC from the cylindrocyclophane biosynthetic gene cluster (Nakamura et al. 2017). The potential role of iron was suggested by its presence in copurified CylC samples (Nakamura et al. 2017). Since then, it was revealed that this family is more frequently found in biosynthetic gene clusters than NHFe-dependent halogenases and might also compete with the occurrence of flavin-dependent halogenases in cyanobacteria (Eusebio et al. 2021). However, the successful expression of homologs such as NocO (halogenase from nocuolactylate biosynthesis) requires the coexpression of the ACP (Wang et al. 2025).

**Copper-dependent halogenases**

**Sequence motifs.** The group of copper-dependent halogenases is represented by one enzyme (ApnU) from the atpenin biosynthesis (Chiang et al. 2025). While the catalytic residues to differentiate between the halogenases and the hydroxylase homologs in this family are not known yet, two conserved motifs, containing His residues and a Cys residue mark the copper-dependent nature of the enzyme (Chiang et al. 2025). Mutation of the His residues to Ala or the Cys residue to Ser result in the loss of activity of the enzyme, proving the cardinal presence of the two motifs (Chiang et al. 2025).

**Activity.** ApnU from atpenin gene cluster is capable of decorating an unactivated carbon center with pseudohildes and halides (Chiang et al. 2025). The copper-dependent halogenase showed bromination and iodination activity besides chlorination and does not utilize fluoride (Chiang et al. 2025).

**Nucleophilic substitution**

**SAM-dependent halogenases**

**Sequence motifs.** Fluorinases have been identified primarily in *Actinomycetes* (Eustáquio et al. 2008; Deng et al. 2006). A 23-residue loop in the conventional fluorinases sets them apart from the other halogenase homologs which are unable to utilize fluoride. However, an unconventional fluorinase from *Archaea* lacks this structural motif (Pardo et al. 2022). On a sequence level, the mining for these enzymes is often based on the catalytic residues (Asp16, Tyr77, Asp210, and Asn215 in FlA$^{MA37}$) (Pardo et al. 2022). A C-terminal small motif (RNAA or YYGG in the archeal fluorinase) can help as well to predict the fluorination activity of the enzyme. Three residues, Tyr70, Trp129, Gly131 might indicate the lack of fluorination activity and can be utilized to engineer enzymes towards better catalytic efficiency (Eustáquio et al. 2008). These residues are replaced by Thr77, Phe156 and Ser158 in the fluorinase (flA) of *Streptomyces cattleya* (Eustáquio et al. 2008).
**Activity.** At the moment, S-Adenosyl-L-Methionine (SAM)-dependent halogenases are the only group of enzymes where we know fluorination in nature. The labeling of fluorinases and chlorinases can also be misleading, as the type of halogenation depends on the laboratory conditions (Deng et al. 2006). An example for that is flA, an enzyme which carries out a reverse reaction. Therefore, the detection of its chlorination activity only works in coupled-enzyme assays with L-amino acid oxidase or adenyl acid deaminase that could shift the reaction from substrates to products (Deng et al. 2006). The substrate scope of these enzymes is less explored, as most studies focus on SAM (Eustáquio et al. 2008; Deng et al. 2006; Schaffrath et al. 2003). Only one study focused on exploring their substrates and reported activity on 2'd-SAM (Cobb et al. 2006).

**Electrophilic substitution**

**Flavin-dependent halogenases**

**Sequence motifs.** Conventional FDHs can be recognized by their flavin-binding site (GxGxxG) (Crowe et al. 2021; Dym and Eisenberg 2001) and a broader sequence motif that marks the tunnel in the structure (Fx.Px.Sx.G) (Crowe et al. 2021). Another motif often seen is WxWxI(P), which is hypothesized to prevent the enzyme from carrying out monooxygenation. However, this motif can be missing from unique members of the family. Such is the case with Bmp5, a decarboxylative brominase acting on pyrrole in the pentabromopseudilin biosynthesis (Agarwal et al. 2014).
Several confirmed catalytic residues enable more in-depth categorization. Bmp2, also from the pentabromopseudilin pathway, shows three residues which hold the key for tetrahalogenation (Tyr302, Phe306, and Ala345) (El Gamal et al. 2016). Moreover, a catalytic Lys residue is often looked for to determine whether the enzyme is acting on tryptophan (Yeh et al. 2007; Lingkon and Bellizzi 2020) and might be cardinal for indole-accepting halogenases (Neubauer et al. 2018). However, it's worth noting that some members of the group containing the residue do not accept tryptophan and are instead acting on indole (Neubauer et al. 2020).

**Activity.** FDHs rely on a reduced flavin cofactor to generate a hypohalite used for halogenation. The reduced flavin can come as the product of a flavin reductase or in single-component enzymes; it can be generated by the halogenase (Gäfe and Niemann 2023). The regioselectivity of FDHs is often mentioned (Chankhamjon et al. 2016; Song et al. 2019; Mori et al. 2019; Domergue et al. 2019), especially when it comes to aromatic substrates (Crowe et al. 2021). However, AoiQ (Liu et al. 2021), CmlS (Podzelinska et al. 2010), and PloK (Ugai et al. 2020) can act on aliphatic substrates. While the family is generally categorized under the enzymes showcasing electrophilic substitution mechanism, a radical pathway might be considered for these members (Chankhamjon et al. 2016). There are also examples of cryptic halogenation reactions. In other families we tend to see cyclization reactions being induced or helped by a halogen atom, in marinopyrrole A and pentabromopseudilin biosynthesis, biaryl coupling is achieved with the help of a halogen (Adak and Moore 2021).

**Vanadium-dependent halogenases**

**Sequence motifs.** For VHPOs, so far, no conserved motifs have been established in genome mining practices. However, catalytic residues have been described across enzymes in this family with different halide-specificities and in different taxon. We see conserved Lys and Ser residues in selective VCPOs, where Lys is replaced by a Threonine in the close homolog hydroxylase, NapH3 (Chen et al. 2022).
The catalytic residues have also been described for bromoperoxidases, including the active sites and the intra-and intermolecular bridges containing Cys residues (Wischang et al. 2012).

**Activity.** Vanadium-dependent haloperoxidases (VHPOs), similarly to FDHs, act on negatively charged positions on the substrate. Despite them having a reputation for being promiscuous, several enantioselective VCPOs have been revealed in *Streptomyces*. This includes NapH1 and NapH4 from the napyradiomycin biosynthesis (Winter et al. 2007), LvcH involved in lavanducyanin (Regioselective Halogenation of Lavand...), Mcl24 and Mcl40 from merochlorin biosynthesis (Kaysser et al. 2012), and MarH1 and MarH3 showing cryptic halogenation reactions on marinone intermediates (Murray et al. 2018).

### GUIDE FOR RESIDUE/MOTIF BASED CATEGORIZATION

The most accessible medium when looking for novel enzymes is the protein sequence. Even though the detection, annotation, and mining of these enzymes often happen based on sequence motifs and known catalytic residues, there was no systematic way of executing these tasks across several halogenase families.

We collected the functionally important residues and conserved motifs to support the precise categorization of halogenating enzymes. This collection serves as support for the categorization done by a computational workflow including a library of family-, substrate-, and variant-specific pHMMs and a repository of the conserved motifs and catalytic residues.

The precise annotation is further assisted by a manually curated database, consisting of more than 100 reviewed entries holding information about the halide-acceptance, results of mutagenesis studies and structural analysis.

The workflow enables the efficient mining and categorization of halogenases and facilitates the detection of known motifs and catalytic residues that are crucial for activity.

**SAM-dependent halogenases with distinct C-terminal motif.** While there are several catalytic residues described in fluorinases, this categorization leans on the C-terminal motif which is RNAA in the bacterial sequences and YYGG in the archeal sequences. Moreover, the search is extended to identify sequences with RNGG and YYAA. Including new combinations of the established pattern allows the identification of novel candidates. The additional features help the search of catalytic residues from known fluorinases and a well described chlorinase.

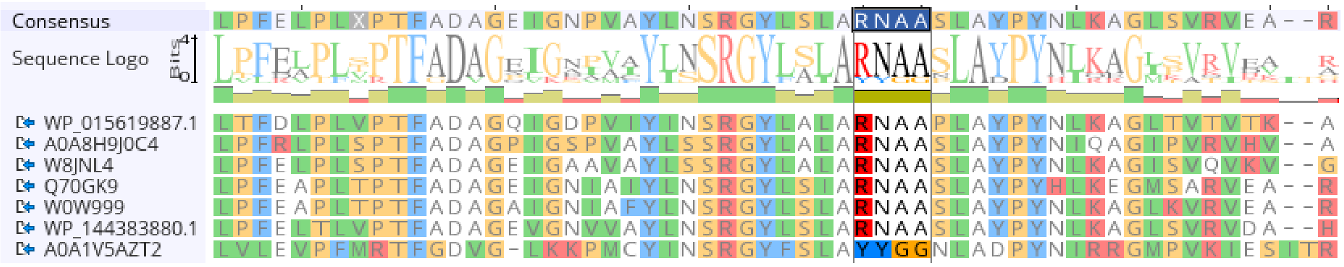

***Supplementary Figure 1 Alignment of the C-terminal of conventional and unconventional halogenases***

MAFFT multiple sequence alignment of conventional (WP_015619887.1, A0A8H9J0C4, W8JNL4, Q70GK9, W0W999, WP_144383880.1) fluorinases with RNAA motif and the unconventional archeal fluorinase with YYGG motif (A0A1V5AZT2) (Jiang et al. 2024)

**Variant B NHFe-dependent halogenases show inconsistency in the catalytic triad.** The HXG/A catalytic triad is often relied on to distinguish between NHFe halogenases and dioxygenases or hydroxylases. While it is in a highly conserved position in variant A enzymes, it shows inconsistencies in the variant B group. A cryptic halogenase JamE in jamaicamide biosynthesis and a fatty acyl halogenase HctB in hectochlorin biosynthesis lack the catalytic G/A residue based on the alignment. During the categorization, the workflow relies on this small motif for the variant A enzymes. However, besides the triad, a HxSxP pattern is also searched for. While this pattern is not based on experimental data, it could help highlight unique enzymes that hit the variant B profile for NHFe halogenases.

The first layer of categorization happens based on the mentioned short patterns in specific regions of the alignment; there are additional features to identify catalytic residues described in variant B NHFe-dependent halogenases. Therefore, we cover both variant A and variant B enzymes.

**Mining for dimetal-carboxylate carboxylate enzymes rely on two motifs.** While the pHMM for this family proved to be specific for the halogenases and the oxygenase or hydroxylase homologs from SwissProt did not hit the profile (Supporting Information), there is further annotation based on the presence of the motif containing the catalytic Glu residue for halogenases and the second motif with H residues that are also present in AurF. The workflow allows comparison to two known enzymes in this family. The structures of CylC and NocO have been studied before (Wang et al. 2025) and can be used to search for the putative catalytic residues in the target sequence.

**Abundance of described catalytic residues to categorize FDHs.** The scientific literature provides several described catalytic residues that can serve as starting points to infer substrate-and regio-specificity (Yeh et al. 2007) or predict tetrahalogenation activity (El Gamal et al. 2016). InterPro consists of two FDH profiles now. The Flavin-dependent halogenase profile from PFAM (PF04820) and the Flavin-dependent tryptophan halogenase profile from the Protein Information Resource (PIR) (PIRSF011396). The curation of substrate-and regio-specific pHMMs help with more accurate activity predictions as the InterPro profiles can hit phenol-like substrate decorating enzymes as well (Supporting Information).

We can use the search for catalytic residues and conserved motif features to carry out in-depth categorization. The hits against the pyrrole-specific pHMM profile can be searched for three positions (Phe306, Ala345 and Tyr302 in bmp2; Val329, Trp369 and Ser325 in Mpy16 and other mono/dihalogenases (El Gamal et al. 2016)). This step helps predict polyhalogenation activity.
Catalytic residues described in BorH (from borregomycin BGC) and MalA (from malbrancheamide biosynthesis) can be used to decide about indole-acceptance (Lingkon and Bellizzi 2020). The workflow allows the comparison to FasV, a recently sescribed multi-site halogenase (Hu et al. 2025) among several conventional tryptophan-and pyrrole-decorating FDHs.

**No established motif when mining for VHPOs.** Unlike in case of flavin-dependent or dimetal-carboxylate halogenases, there are no established motifs that are used for enzyme-mining practices for VHPOs. The catalytic Lys and Ser residues serve as steppingstones to distinguish enantioselective chloroperoxidase. However, these residues are not conserved in bromo-or iodoperoxidases. The workflow allows the comparison of the target sequence(s) to known iodoperoxidases from *Zobellia galactanivorans* and bromoperoxidases from *Acaryochloris marina*, *Ascophyllum nodosum* and *Corallina pilulifera* to provide further guidance for inferring activity.
The workflow checks for broader motifs in VBPO pHMM hits which contain amino acids forming the active site and Cys residues which are responsible for intra-and intermolecular bridges (Wischang et al. 2012).

### PHYLOGENETIC TREE OF NHFE-DEPENDENT ENZYMES

MSA was generated with MAFFT followed by phylogenetic tree construction with FastTree in Geneious Prime 2024.0.4

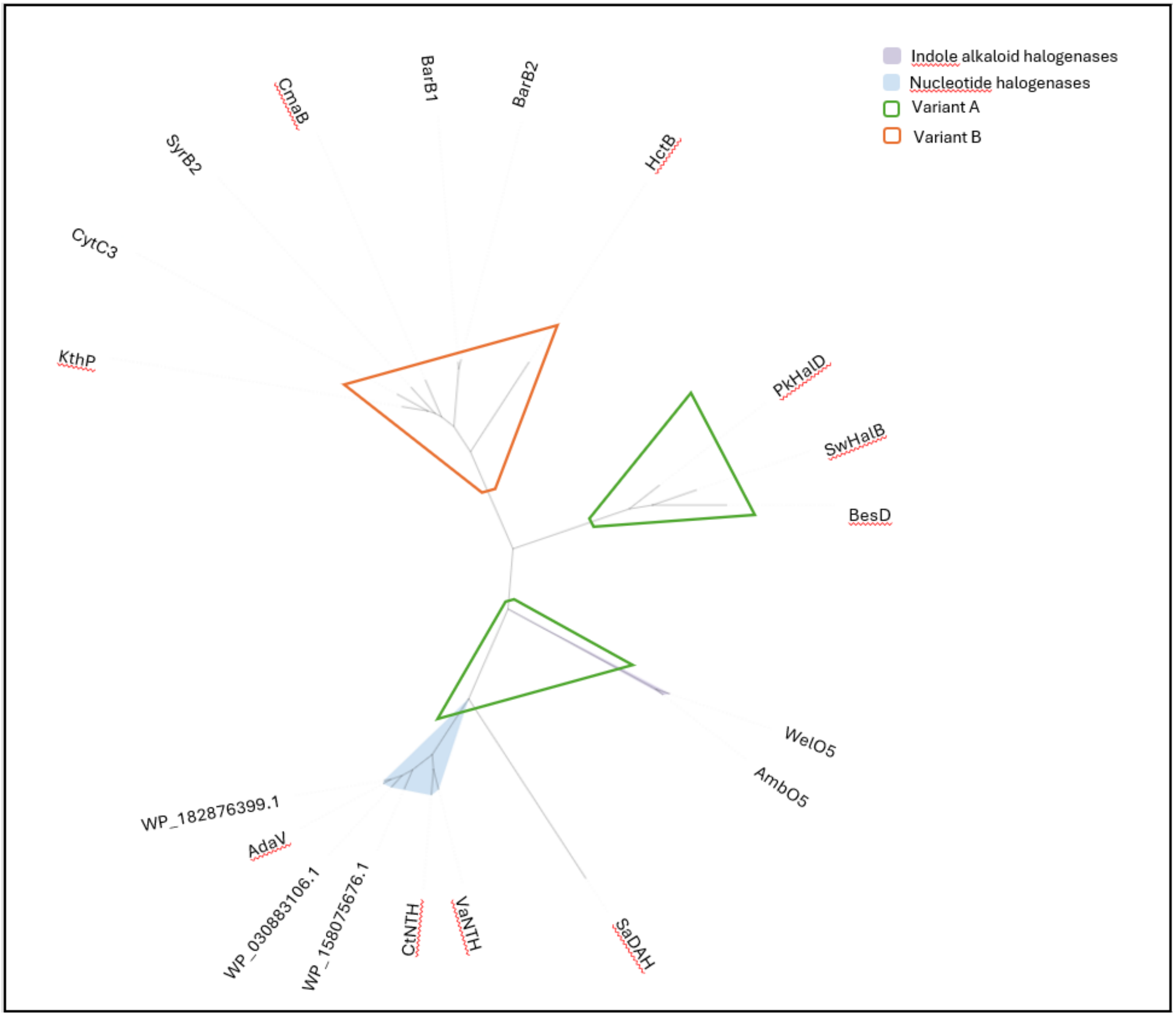

***Supplementary Figure 2 Phylogenetic tree of NHFe-dependent enzymes***
The workflow supports the substrate prediction of NHFe-dependent enzymes based on homology, as they cluster according to substrate and variance (variant A or B) on the phylogenetic tree. AmbO5 (AKP23998.1), WelO5 (5TRQ), BesD (6NIE), SwHalB (SDN46247.1), PkHalD (WP_046063366.1), HctB (Q1EDB4), BarB2 (Q8GAQ8), BarB1 (Q8GAQ9), CmaB (A0A8T0C957), SyrB2 (2FCU), CytC3 (GJB), KthP

(W7T5C7), WP_182876399.1 (dAMP), AdaV (AKQ99303.1, dAMP), WP_030883106.1 (dAMP), WP_158075676.1 (dAMP), CtNTH (WP_217394979.1, dGMP), VaNTH (WP_204007738.1, dGMP), SaDAH (QJD15032.1).

### UniRef50 ANALYSIS

**Putative SAM-dependent fluorinases in an undescribed cluster.** The SAM-dependent non-fluorinase and fluorinase models revealed several clusters unconnected to the known halogenases. The general workflow for the SAM-dependent family searches for YYGG and RNAA, known C-terminal motifs in bacterial and in the archeal fluorinases. We also extended the search for RNGG and YYAA. Searching for the C-terminal motifs allowed us to identify two needles in the haystack. A large cluster without any known chlorinase or fluorinase holds two sequences with the RNAA motif. Both are bacterial sequences, one from an unclassified bacterium (A0A0S8JKK5), another from *Planctomycetota* (A0A2D6A2E4). We used the features for comparing to known enzymes when it came to the known chlorinases and looked for the catalytic residues from flA, flA(PtaU1), flA1 and flA4. However, these residues seem to be specific to the known cluster as it didn't give any match from groups not connected to the known enzymes.
When the chlorinase pHMM hits were compared to the known *Salinospora tropica* catalytic residues. However, we could not identify not-validated enzymes with these residues.

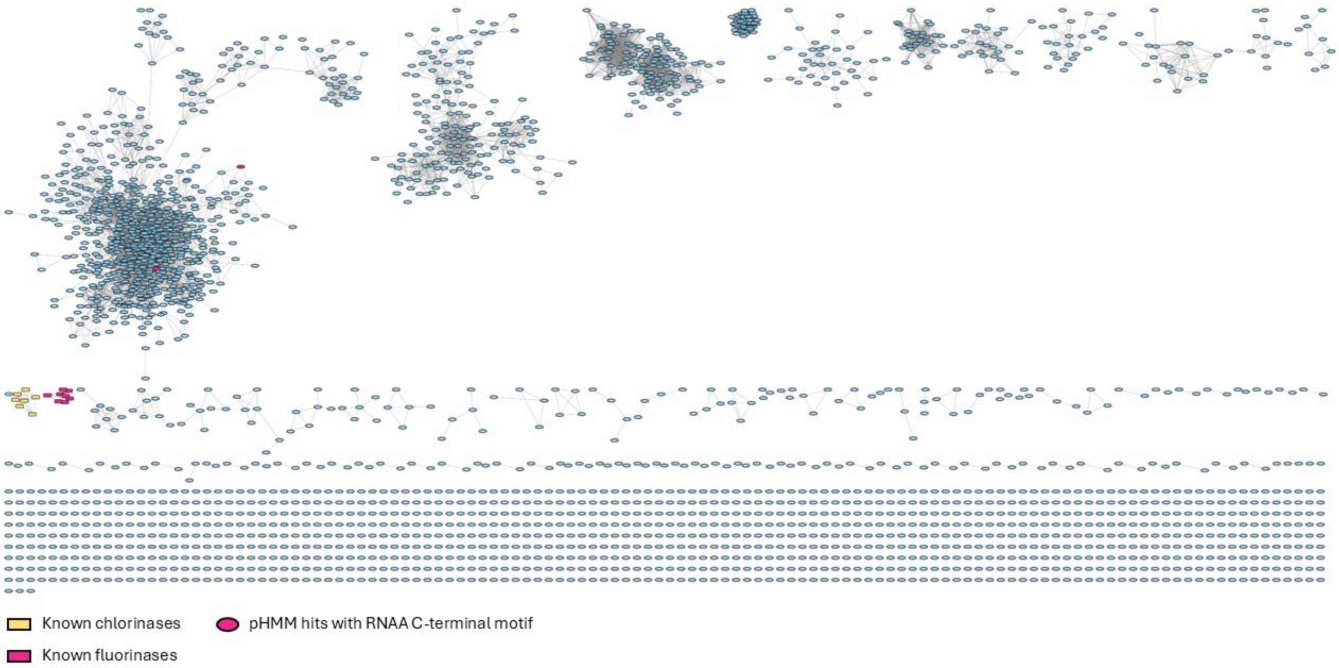

***Supplementary Figure 3 SAM-dependent enzymes***
The network shows the currently known halogenases in two small clusters. Separated from them is a large group revealed with two enzymes possessing the C-terminal RNAA motif

**Known unconventional FDHs contain the tunnel-lining motif.** The workflow highlighted the presence of the Fx.Px.Sx.G pattern in unconventional FDHs and the network shows two undescribed close homologs with the same motif, connected to the known FDHs.
The pHMM curated from whole sequence alignments of AetF, VatD, JamD and PhmJ picked up 193 sequences from UniRef50, including AetF. The application of regular expressions in the computational workflow allowed the systematic detection of the conventional Fx.Px.Sx.G motif. However, the range in the pHMM alignment which searches for the motif is quite long and the motif is broad. This can result in false positives, if we do not consider the conserved putative flavin-binding motif in the hits. The network highlights enzymes possessing this motif that happen to show GxGxxA pattern at the N-terminal, possibly flavin-binding site as well. Due to their unique ability of halogenating terminal alkynes (Lukowski et al. 2023), which is not characteristic of this enzymes family based on what we know so far, the workflow can accelerate the mining for new biocatalysts.

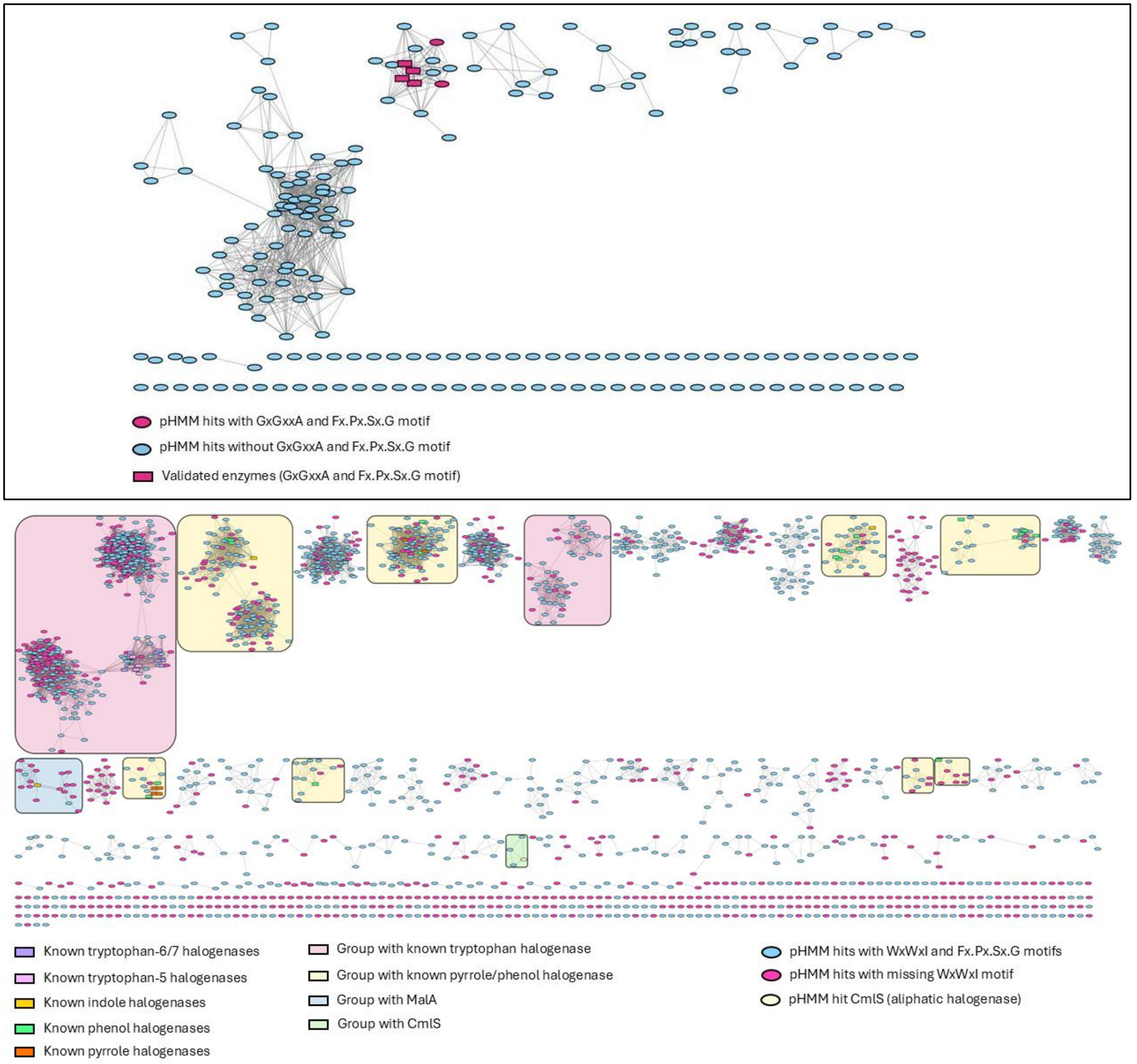

***Supplementary Figure 4 SSN of conventional FDHs***

Clusters with known FDHs scatter in the network, separated by clusters with halogenase candidates with the Fx.Px.Sx.G motif, but many of these showing inconsistencies with the WxWxI motif.

Likely due to the highly conserved flavin-binding motif, flavin-dependent monooxygenases and oxidoreductases often hit the tryptophan halogenase profile in InterPro and lead to their misannotation as halogenase. The FDH profiles exclude the conserved flavin-binding region from the MSA that the pHMM was built from to make them more halogenase-specific. However, using the pHMM for the general detection of the halogenase members of this family still picked up 14545 sequences. Searching for the WxWxI and Fx.Px.Sx.G motif helped to narrow down the dataset to 1960 sequences. As the WxWxI motif can be missing from sequences with unique activity (e.g. Bmp5), these proteins are highlighted in the network. The curated general FDH profile is also suitable for identifying sequences that act on aliphatic molecules. We see that CmlS is among the hits even though it was not part of MSA the model was built from.

**Known variant A enzymes are connected to a variant B cluster among NHFe-dependent halogenases.** Even though the pHMMs seemed to separate variant A and variant B enzymes well, the indole alkaloid halogenases are loosely connected to a cluster containing variant B hits without any described sequence in UniRef50. Deciding whether these are true variant B enzymes or they mean a new group of unconventional variant A sequences requires further experimental validation, which was out of the scope of this study.

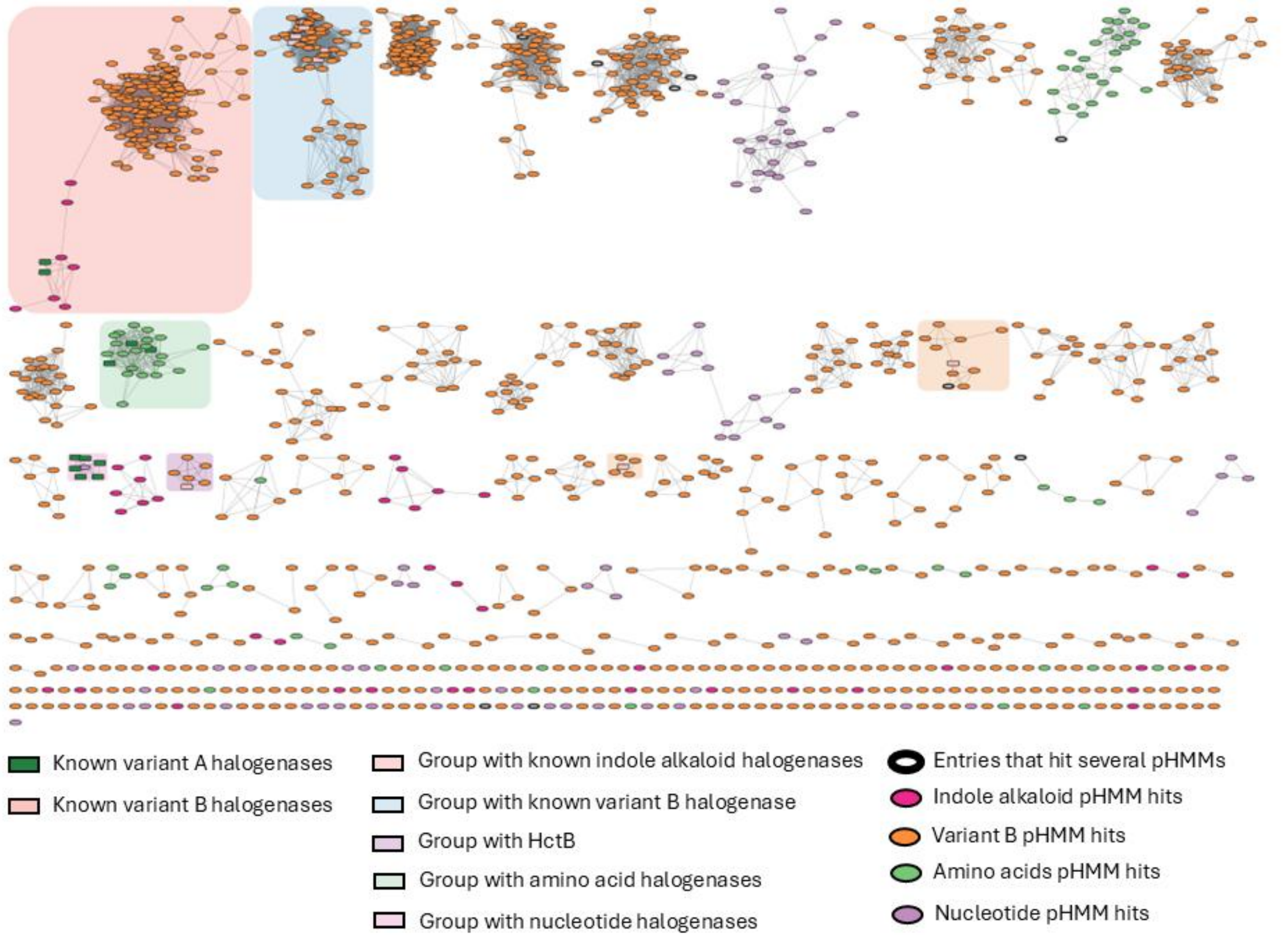

***Supplementary Figure 5 SSN of NHFe-dependent enzymes***
*NHFe-dependent enzymes reveal a large space waiting for experimental validation. The SSN highlights the selective nature of the variant A- and variant B-and substrate-specific pHMMs. Sequences which hit more than one profile, usually the variant B and indole alkaloid, but sometimes two variant A profiles are represented by bold, empty nodes.*

The small clusters, belonging to the variant B category, holds the known cryptic halogenases and could point to biocatalysts responsible for cyclopropane ring formation, like CurA or Jame. It was very rare that one sequence would be a hit to several profiles. This highlights the selective nature of the pHMMs and could mean novel activity for the enzymes belonging to multiple pHMM.

**Dimetal-carboxylate enzymes.** Known dimetal-carboxylate enzymes belong to the same cluster in the SSN with several homologous sequences possessing the ExxQExxH and HxxDExxH motifs characteristic of the halogenases in the family. Some of the enzymes seem to possess only the HxxDExxH motif, which can indicate a similar function to AurF, a 4-aminobenzoate N-oxygenase. Furthermore, when searching for the HxxDExxH motif, some of the sequences, including BrtJ (halogenase from bartoloside biosynthesis) didn't return any hits. This was intriguing because looking at the MSA, BrtJ shows both motifs. The pHMM alignment showed that BrtJ has several deletions before the second motif. The searched region was adjusted to exclude the deletions and the workflow became suitable for identifying the second motif in BrtJ and its homologs as well.
A previous study claimed that dimetal-carboxylate halogenases might be represented in greater numbers compared to NHFe-dependent and Flavin-dependent halogenases (Eusebio et al. 2021). This was not reflected in the UniRef50 dataset. The reason might be the homology-reduction or the distribution of cyanobacterial sequences or the strictness of the pHMM. However, miscellaneous nodes and separate clusters from the one containing the known sequences did not show the presence of both motifs and only in few cases did the second motif occur

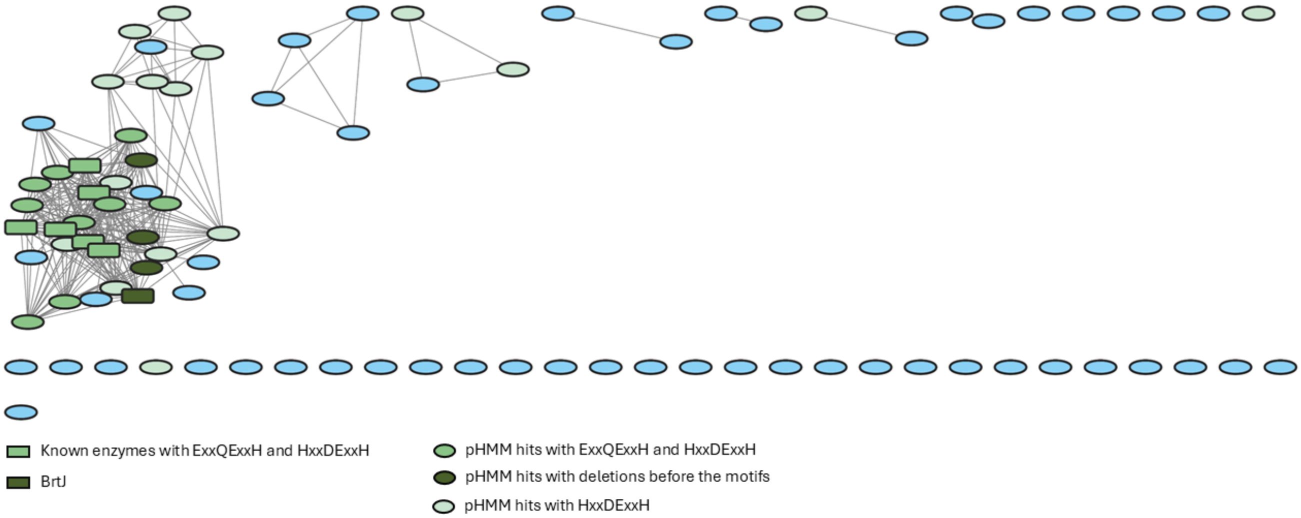

***Supplementary Figure 6 SSN of dimetal-carboxylate enzymes***

The SSN shows known enzymes. BrtJ and its homologs are shown in dark green, as the search for the second motif needed adjustments due to the deletions in the alignment of the pHMM. The network shows all the known enzymes and putative halogenases in one cluster.

**Copper-dependent enzymes.** The enzyme, ApnU from atpenin biosynthesis (Chiang et al. 2025) can guide us in the mining process for copper-dependent halogenases. The presence of two HxxHC motifs is prevalent among copper-dependent enzymes (including hydroxylases and the recently explored halogenase) as the SSN shows.

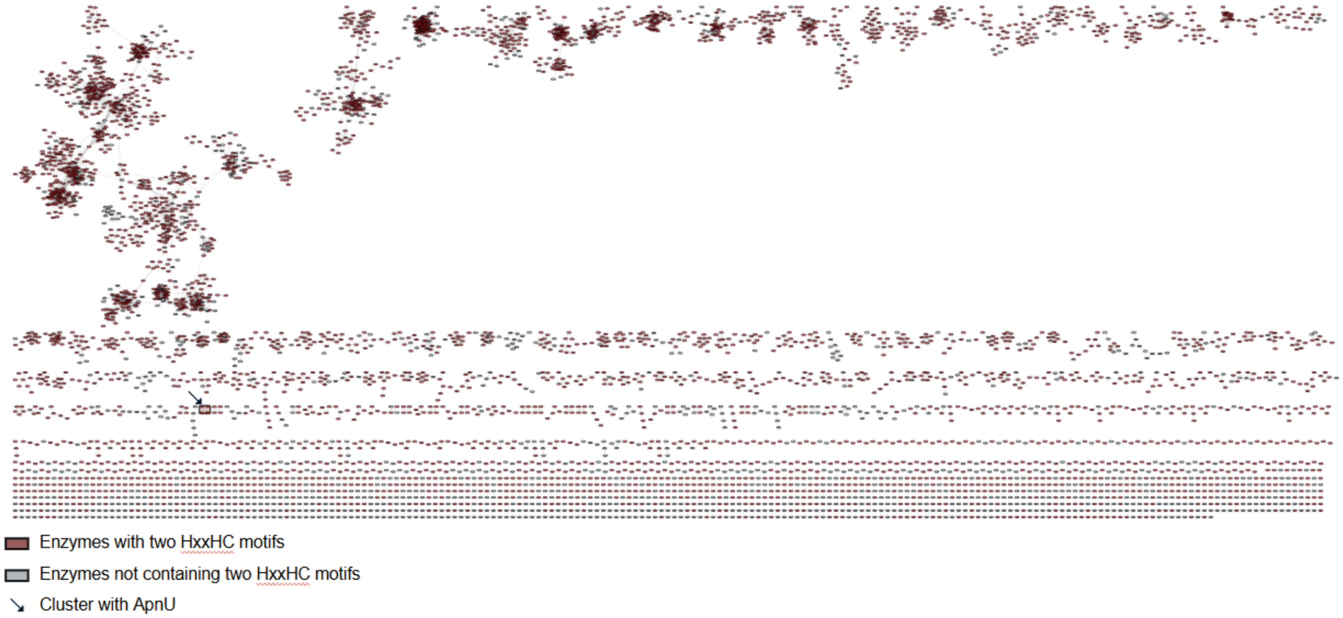

***Supplementary Figure 7 SSN of copper-dependent enzymes***

The SSN highlights enzymes with two HxxHC motifs. The arrow points to the cluster containing the recently described copper-dependent halogenase from atpenin biosynthesis (Chiang et al. 2025)

**COMMERCIAL SUPPLIERS FOR THE MCD ASSAY of *Rhoba*VHPO**

Synthetic genes were ordered from twist, restriction enzymes were supplied by NEB, TherfmoFisher provided the media, sigma Aldrich supplied antibiotics, IPTG and small molecule chemicals.

1. Pratter, S. M., Light, K. M., Solomon, E. I., & Straganz, G. D. (2014). The role of chloride in the mechanism of O2 activation at the mononuclear nonheme Fe (II) center of the halogenase HctB. *Journal of the American Chemical Society*, *136*(26), 9385-9395.
2. Hillwig, M. L., & Liu, X. (2014). A new family of iron-dependent halogenases acts on freestanding substrates. *Nature chemical biology*, *10*(11), 921-923.
3. Ni, J., Zhuang, J., Shi, Y., Chiang, Y. C., & Cheng, G. J. (2024). Discovery and substrate specificity engineering of nucleotide halogenases. *Nature Communications*, *15*(1), 5254.
4. 46
5. Neugebauer, M. E., Sumida, K. H., Pelton, J. G., McMurry, J. L., Marchand, J. A., & Chang, M. C. (2019). A family of radical halogenases for the engineering of amino-acid-based products. *Nature chemical biology*, *15*(10), 1009-1016.
6. Blasiak, L. C., Vaillancourt, F. H., Walsh, C. T., & Drennan, C. L. (2006). Crystal structure of the non-haem iron halogenase SyrB2 in syringomycin biosynthesis. *Nature*, *440*(7082), 368-371.
7. Kim, C. Y., Mitchell, A. J., Glinkerman, C. M., Li, F. S., Pluskal, T., & Weng, J. K. (2020). The chloroalkaloid (−)-acutumine is biosynthesized via a Fe (II)-and 2-oxoglutarate-dependent halogenase in Menispermaceae plants. *Nature Communications*, *11*(1), 1867.
8. Zhang, X., Wang, Z., Gao, J., & Liu, W. (2020). Chlorination versus hydroxylation selectivity mediated by the non-heme iron halogenase WelO5. *Physical Chemistry Chemical Physics*, *22*(16), 8699-8712.
9. Zhai, G., Gong, R., Lin, Y., Zhang, M., Li, J., Deng, Z., ... & Zhang, Z. (2022). Structural insight into the catalytic mechanism of non-heme iron halogenase AdaV in 2’-chloropentostatin biosynthesis. *ACS Catalysis*, *12*(22), 13910-13920.
10. Hillwig, M. L., & Liu, X. (2014). A new family of iron-dependent halogenases acts on freestanding substrates. *Nature chemical biology*, *10*(11), 921-923.
11. Ni, J., Zhuang, J., Shi, Y., Chiang, Y. C., & Cheng, G. J. (2024). Discovery and substrate specificity engineering of nucleotide halogenases. *Nature Communications*, *15*(1), 5254.
12. Zhai, G., Gong, R., Lin, Y., Zhang, M., Li, J., Deng, Z., ... & Zhang, Z. (2022). Structural insight into the catalytic mechanism of non-heme iron halogenase AdaV in 2’-chloropentostatin biosynthesis. *ACS Catalysis*, *12*(22), 13910-13920.
13. Vaillancourt, F. H., Yeh, E., Vosburg, D. A., O'Connor, S. E., & Walsh, C. T. (2005). Cryptic chlorination by a non-haem iron enzyme during cyclopropyl amino acid biosynthesis. *Nature*, *436*(7054), 1191-1194.
14. Neumann, C. S., & Walsh, C. T. (2008). Biosynthesis of (−)-(1 S, 2 R)-allocoronamic acyl thioester by an FeII-dependent halogenase and a cyclopropane-forming flavoprotein. *Journal of the American Chemical Society*, *130*(43), 14022-14023.
15. Matthews, M. L., Krest, C. M., Barr, E. W., Vaillancourt, F. H., Walsh, C. T., Green, M. T., ... & Bollinger Jr, J. M. (2009). Substrate-triggered formation and remarkable stability of the C− H bond-cleaving chloroferryl intermediate in the aliphatic halogenase, SyrB2. *Biochemistry*, *48*(20), 4331-4343.
16. Galonić, D. P., Vaillancourt, F. H., & Walsh, C. T. (2006). Halogenation of unactivated carbon centers in natural product biosynthesis: trichlorination of leucine during barbamide biosynthesis. *Journal of the American Chemical Society*, *128*(12), 3900-3901.
17. Ueki, M., Galonić, D. P., Vaillancourt, F. H., Garneau-Tsodikova, S., Yeh, E., Vosburg, D. A., ... & Walsh, C. T. (2006). Enzymatic generation of the antimetabolite γ, γ-dichloroaminobutyrate by NRPS and mononuclear iron halogenase action in a streptomycete. *Chemistry & Biology*, *13*(11), 1183-1191.
18. Khare, D., Wang, B., Gu, L., Razelun, J., Sherman, D. H., Gerwick, W. H., ... & Smith, J. L. (2010). Conformational switch triggered by α-ketoglutarate in a halogenase of curacin A biosynthesis. *Proceedings of the National Academy of Sciences*, *107*(32), 14099-14104.
19. Gu, L., Wang, B., Kulkarni, A., Geders, T. W., Grindberg, R. V., Gerwick, L., ... & Sherman, D. H. (2009). Metamorphic enzyme assembly in polyketide diversification. *Nature*, *459*(7247), 731-735.
20. Pratter, S. M., Ivkovic, J., Birner-Gruenberger, R., Breinbauer, R., Zangger, K., & Straganz, G. D. (2014). More than just a halogenase: modification of fatty acyl moieties by a trifunctional metal enzyme. *ChemBioChem*, *15*(4), 567-574.
21. Jiang, W., Heemstra Jr, J. R., Forseth, R. R., Neumann, C. S., Manaviazar, S., Schroeder, F. C., ... & Walsh, C. T. (2011). Biosynthetic chlorination of the piperazate residue in kutzneride biosynthesis by KthP. *Biochemistry*, *50*(27), 6063-6072.
22. Wang, M. L., Glasser, N. R., Nair, M. A., Krebs, C., Martin Bollinger Jr, J., & Balskus, E. P. (2025). Biochemical Studies of a Cyanobacterial Halogenase Support the Involvement of a Dimetal Cofactor. *Biochemistry*, *64*(10), 2173-2180.
23. Simurdiak, M., Lee, J., & Zhao, H. (2006). A New Class of Arylamine Oxygenases: Evidence that p-Aminobenzoate N-Oxygenase (AurF) is a Di-iron Enzyme and Further Mechanistic Studies. *ChemBioChem*, *7*(8), 1169-1172.
24. Wang, M. L., Glasser, N. R., Nair, M. A., Krebs, C., Martin Bollinger Jr, J., & Balskus, E. P. (2025). Biochemical Studies of a Cyanobacterial Halogenase Support the Involvement of a Dimetal Cofactor. *Biochemistry*, *64*(10), 2173-2180.
25. Nakamura, H., Schultz, E. E., & Balskus, E. P. (2017). A new strategy for aromatic ring alkylation in cylindrocyclophane biosynthesis. *Nature Chemical Biology*, *13*(8), 916-921.

26. Eusebio, N., Rego, A., Glasser, N. R., Castelo-Branco, R., Balskus, E. P., & Leão, P. N. (2021). Distribution and diversity of dimetal-carboxylate halogenases in cyanobacteria. *BMC genomics*, *22*(1), 633.
27. Wang, M. L., Glasser, N. R., Nair, M. A., Krebs, C., Martin Bollinger Jr, J., & Balskus, E. P. (2025). Biochemical Studies of a Cyanobacterial Halogenase Support the Involvement of a Dimetal Cofactor. *Biochemistry*, *64*(10), 2173-2180.
28. Pardo, I., Bednar, D., Calero, P., Volke, D. C., Damborsky, J., & Nikel, P. I. (2022). A nonconventional Archaeal fluorinase identified by in silico mining for enhanced fluorine biocatalysis. *ACS catalysis*, *12*(11), 6570-6577.
29. Eustáquio, A. S., Pojer, F., Noel, J. P., & Moore, B. S. (2008). Discovery and characterization of a marine bacterial SAM-dependent chlorinase. *Nature Chemical Biology*, *4*(1), 69-74.
30. Deng, H., Cobb, S. L., McEwan, A. R., McGlinchey, R. P., Naismith, J. H., O'Hagan, D., ... & Spencer, J. B. (2006). The fluorinase from Streptomyces cattleya is also a chlorinase. *Angewandte Chemie (International ed. in English)*, *45*(5), 759.
31. Eustáquio, A. S., Pojer, F., Noel, J. P., & Moore, B. S. (2008). Discovery and characterization of a marine bacterial SAM-dependent chlorinase. *Nature Chemical Biology*, *4*(1), 69-74.
32. Deng, H., Cobb, S. L., McEwan, A. R., McGlinchey, R. P., Naismith, J. H., O'Hagan, D., ... & Spencer, J. B. (2006). The fluorinase from Streptomyces cattleya is also a chlorinase. *Angewandte Chemie (International ed. in English)*, *45*(5), 759.
33. Schaffrath, C., Deng, H., & O'Hagan, D. (2003). Isolation and characterisation of 5'-fluorodeoxyadenosine synthase, a fluorination enzyme from Streptomyces cattleya. *FEBS letters*, *547*(1-3), 111-114.
34. Cobb, S. L., Deng, H., McEwan, A. R., Naismith, J. H., O'hagan, D., & Robinson, D. A. (2006). Substrate specificity in enzymatic fluorination. The fluorinase from Streptomyces cattleya accepts 2'-deoxyadenosine substrates. *Organic & biomolecular chemistry*, *4*(8), 1458-1460.
35. Crowe, C., Molyneux, S., Sharma, S. V., Zhang, Y., Gkotsi, D. S., Connaris, H., & Goss, R. J. (2021). Halogenases: a palette of emerging opportunities for synthetic biology–synthetic chemistry and C–H functionalisation. *Chemical Society Reviews*, *50*(17), 9443-9481.
36. Dym, O., & Eisenberg, D. (2001). Sequence-structure analysis of FAD-containing proteins. *Protein Science*, *10*(9), 1712-1728.
37. Agarwal, V., El Gamal, A. A., Yamanaka, K., Poth, D., Kersten, R. D., Schorn, M., ... & Moore, B. S. (2014). Biosynthesis of polybrominated aromatic organic compounds by marine bacteria. *Nature chemical biology*, *10*(8), 640-647.
38. El Gamal, A., Agarwal, V., Diethelm, S., Rahman, I., Schorn, M. A., Sneed, J. M., ... & Moore, B. S. (2016). Biosynthesis of coral settlement cue tetrabromopyrrole in marine bacteria by a uniquely adapted brominase–thioesterase enzyme pair. *Proceedings of the National Academy of Sciences*, *113*(14), 3797-3802.
39. Yeh, E., Blasiak, L. C., Koglin, A., Drennan, C. L., & Walsh, C. T. (2007). Chlorination by a long-lived intermediate in the mechanism of flavin-dependent halogenases. *Biochemistry*, *46*(5), 1284-1292.
40. Lingkon, K., & Bellizzi III, J. J. (2020). Structure and Activity of the Thermophilic Tryptophan-6 Halogenase BorH. *ChemBioChem*, *21*(8), 1121-1128.
41. Neubauer, P. R., Widmann, C., Wibberg, D., Schröder, L., Frese, M., Kottke, T., ... & Sewald, N. (2018). A flavin-dependent halogenase from metagenomic analysis prefers bromination over chlorination. *PLoS One*, *13*(5), e0196797.
42. Neubauer, P. R., Pienkny, S., Wessjohann, L., Brandt, W., & Sewald, N. (2020). Predicting the Substrate Scope of the Flavin-Dependent Halogenase BrvH. *ChemBioChem*, *21*(22), 3282-3288.
43. Gäfe, S., & Niemann, H. H. (2023). Structural basis of regioselective tryptophan dibromination by the single-component flavin-dependent halogenase AetF. *Biological Crystallography*, *79*(7), 596-609.
44. Chankhamjon, P., Tsunematsu, Y., Ishida-Ito, M., Sasa, Y., Meyer, F., Boettger-Schmidt, D., ... & Hertweck, C. (2016). Regioselective Dichlorination of a non-activated Aliphatic Carbon Atom and Phenolic Bismethylation by a multifunctional fungal flavoenzyme. *Angewandte Chemie International Edition*, *55*(39), 11955-11959.
45. Song, R., Shi, H., Zhu, J., Wang, H., & Shen, Y. (2019). A single-component flavoenzyme catalyzed regioselective halogenation of pyrone in the biosynthesis of venemycins. *ACS Chemical Biology*, *14*(12), 2533-2537.
46. Mori, S., Pang, A. H., Thamban Chandrika, N., Garneau-Tsodikova, S., & Tsodikov, O. V. (2019). Unusual substrate and halide versatility of phenolic halogenase PltM. *Nature communications*, *10*(1), 1255.
47. Domergue, J., Erdmann, D., Fossey-Jouenne, A., Petit, J. L., Debard, A., de Berardinis, V., ... & Zaparucha, A. (2019). Xszen FHal, a novel Tryptophan 5-halogenase from Xenorhabdus szentirmaii. *AMB Express*, *9*(1), 175.
48. Crowe, C., Molyneux, S., Sharma, S. V., Zhang, Y., Gkotsi, D. S., Connaris, H., & Goss, R. J. (2021). Halogenases: a palette of emerging opportunities for synthetic biology–synthetic chemistry and C–H functionalisation. *Chemical Society Reviews*, *50*(17), 9443-9481.
49. Liu, M., Ohashi, M., Hung, Y. S., Scherlach, K., Watanabe, K., Hertweck, C., & Tang, Y. (2021). AoiQ catalyzes geminal dichlorination of 1, 3-diketone natural products. *Journal of the American Chemical Society*, *143*(19), 7267-7271.
50. Podzelinska, K., Latimer, R., Bhattacharya, A., Vining, L. C., Zechel, D. L., & Jia, Z. (2010). Chloramphenicol biosynthesis: the structure of CmlS, a flavin-dependent halogenase showing a covalent flavin–aspartate bond. *Journal of molecular biology*, *397*(1), 316-331.

51. Ugai, T., Minami, A., Tanaka, S., Ozaki, T., Liu, C., Shigemori, H., ... & Oikawa, H. (2020). Biosynthetic machinery of 6-hydroxymellein derivatives leading to cyclohelminthols and palmaenones. *ChemBioChem*, *21*(3), 360-367.
52. Chankhamjon, P., Tsunematsu, Y., Ishida-Ito, M., Sasa, Y., Meyer, F., Boettger-Schmidt, D., ... & Hertweck, C. (2016). Regioselective Dichlorination of a non-activated Aliphatic Carbon Atom and Phenolic Bismethylation by a multifunctional fungal flavoenzyme. *Angewandte Chemie International Edition*, *55*(39), 11955-11959.
53. Adak, S., & Moore, B. S. (2021). Cryptic halogenation reactions in natural product biosynthesis. *Natural product reports*, *38*(10), 1760-1774.
54. Chen, P. Y. T., Adak, S., Chekan, J. R., Liscombe, D. K., Miyanaga, A., Bernhardt, P., ... & Moore, B. S. (2022). Structural basis of stereospecific vanadium-dependent haloperoxidase family enzymes in napyradiomycin biosynthesis. *Biochemistry*, *61*(17), 1844-1852.
55. Wischang, D., Radlow, M., Schulz, H., Vilter, H., Viehweger, L., Altmeyer, M. O., ... & Hartung, J. (2012). Molecular cloning, structure, and reactivity of the second bromoperoxidase from Ascophyllum nodosum. *Bioorganic chemistry*, *44*, 25-34.
56. Winter, J. M., Moffitt, M. C., Zazopoulos, E., McAlpine, J. B., Dorrestein, P. C., & Moore, B. S. (2007). Molecular basis for chloronium-mediated meroterpene cyclization: cloning, sequencing, and heterologous expression of the napyradiomycin biosynthetic gene cluster. *Journal of Biological Chemistry*, *282*(22), 16362-16368.
57. Baumgartner, J. T., & McKinnie, S. M. (2024). Regioselective Halogenation of Lavanducyanin by a Site-Selective Vanadium-Dependent Chloroperoxidase. *Organic Letters*, *26*(27), 5725-5730.
58. Kaysser, L., Bernhardt, P., Nam, S. J., Loesgen, S., Ruby, J. G., Skewes-Cox, P., ... & Moore, B. S. (2012). Merochlorins A–D, cyclic meroterpenoid antibiotics biosynthesized in divergent pathways with vanadium-dependent chloroperoxidases. *Journal of the American Chemical Society*, *134*(29), 11988-11991.
59. Murray, L. A., McKinnie, S. M., Pepper, H. P., Erni, R., Miles, Z. D., Cruickshank, M. C., ... & George, J. H. (2018). Total synthesis establishes the biosynthetic pathway to the naphterpin and marinone natural products. *Angewandte Chemie International Edition*, *57*(34), 11009-11014.
60. Jiang, Y., Yao, M., Niu, H., Wang, W., He, J., Qiao, B., ... & Yuan, Y. (2024). Enzyme engineering renders chlorinase the activity of fluorinase. *Journal of Agricultural and Food Chemistry*, *72*(2), 1203-1212.
61. Chiang, C. Y., Ohashi, M., Le, J., Chen, P. P., Zhou, Q., Qu, S., ... & Tang, Y. (2025). Copper-dependent halogenase catalyses unactivated C− H bond functionalization. *Nature*, *638*(8049), 126-132.
